# What is it like to be a blob? A distributed, decentred and context-sensitive neurobiology of consciousness

**DOI:** 10.64898/2026.09.02.745341

**Authors:** Jeremy I Skipper, Christopher Timmermann, Rosalind McAlpine, Krisztina Jedlovszky, Magdalena Jaglinska, Greg Cooper, George Blackburne

## Abstract

Consciousness is often studied by asking whether particular brain regions or systems are privileged. We tested a distributed, decentred and context-sensitive alternative using 579 consciousness-focused neuroimaging studies. Semantic clustering of titles and abstracts identified seven domains whose corrected meta-analytic maps showed limited overlap and no invariant cross-domain core. These maps occupied 18.0% of grey matter while extending across 91.0% of brain regions. After controlling for reporting topography, distributed cognitive, sensorimotor and organismic-affective maps reconstructed the residual consciousness landscape better than spatial surrogates. Organismic-affective maps contributed most, but functional contributions varied contextually across domains. Theory-associated regional loci added no additional meaningful prediction. Information carried by the functional families was both redundant and synergistic. Coactivation-network and hypergraph analyses revealed a shared network repertoire but context-dependent higher-order coalitions. These results argue against an invariant core, supporting a distributed neurobiology of consciousness in which specialised systems are weighted and combined differently across contexts.

## 1 Introduction

The search for the ‘neural correlates of consciousness’ has often focused on which brain regions or systems matter most. Global workspace theory proposes that information becomes conscious when it gains access to a shared workspace and is made broadly available to otherwise specialised processors [1]. Its neuronal counterpart suggests that this workspace extends across prefrontal, parietotemporal and cingulate cortex, with prefrontal ignition and broadcasting particularly prominent [2–4]. Integrated information theory identifies consciousness with whole-system causal organisation irreducible to its parts and emphasises a posterior cortical ‘hot zone’ [5, 6]. Recurrent-processing theories emphasise local feedback, particularly in sensory cortices [7], whereas higher-order theories posit representations of first-order states, often linked to dorsolateral prefrontal and putative metacognitive systems [8]. Some proposals privilege the thalamus [9], brainstem and other subcortical structures [10–12]. None is a simple one-region theory, and each includes distributed operations. Yet their neural implementations are often adjudicated by whether a territory is privileged, necessary or sufficient, most visibly in the front-versus-back debate [13], formalised by an adversarial comparison of prefrontal and posterior predictions [14]. The unresolved issue is not whether local structures matter, but whether any territory retains explanatory privilege across the contexts in which consciousness has been studied.

A less locus-centred account shifts emphasis from privileged loci to distributed, differentiated systems whose organisation can vary across contexts. Dennett and Kinsbourne’s multiple-drafts model rejected a Cartesian theatre in which distributed discriminations are assembled for a central observer [15, 16]. Parallel processes instead acquire graded influence over cognition and behaviour, later expressed as ‘fame in the brain’ [16–18]. Temporo-spatial theory attributes consciousness to intrinsic spatial and temporal dynamics rather than any region [19]. Tononi and Edelman’s dynamic-core hypothesis proposed successive differentiated, re-entrantly integrated thalamocortical coalitions [20]. Relatedly, central-thalamic stimulation shows how a focal intervention can reorganise distributed cortical dynamics [21]. At a more circumscribed scale, Zeki proposed partly dissociable micro-consciousnesses across specialised visual systems [22]. At the cellular scale, apical amplification offers a context-sensitive gate [23]. Taken together, this literature separates local causal importance from anatomical privilege. A local mechanism may gate or reorganise a conscious state without itself constituting a fixed neural locus. Conscious states may instead be realised by context-specific combinations of distributed functional systems rather than by a fixed network unique to consciousness.

A harder challenge for a distributed account is subjectivity, because the first-person character of experience might seem to privilege brain systems that represent and regulate the organism. This is explicit in Parvizi and Damasio, who ground core consciousness and self in homeostatic regulation and brainstem somato-sensing structures [10, 24, 25]. Building on Damasio and related upper-brainstem accounts [11], Solms identifies affect as elemental consciousness and upper-brainstem systems as its source [26]. More broadly, Seth’s interoceptive predictive-coding model spans brainstem, subcortical, insular, cingulate and orbitofrontal systems, but gives anterior insula a key role in conscious presence [27, 28]. While these accounts give particular loci privileged roles, the processes they invoke to ground subjectivity are more distributed and differentiated than this emphasis suggests. Meta-analyses of interoception and autonomic regulation implicate distributed cortical and subcortical systems [29, 30], while a synthesis spanning interoceptive, exteroceptive and higher-level self-processing found overlapping but differentiated insular, cingulate, medial prefrontal and temporoparietal systems [31]. Meta-analyses of bodily self-consciousness likewise identify distributed and partly distinct systems for bodily integration, ownership and agency [32–34]. Affective processing is also distributed across cortical-subcortical systems [35]. A systematic review of electrical stimulation found disturbances of bodily self-consciousness across several cortical and medial-temporal sites [36].

Together, this evidence weakens anatomical privilege without establishing context sensitivity, since subjectivity could still depend on a relatively stable repertoire of bodily-affective systems. We propose instead that an adequate account of subjectivity must accommodate context. On this stronger version of the distributed account, bodily and affective processing makes a context-sensitive contribution to organism-relative significance within a wider functional architecture. Bodily condition and value then situate processing relative to homeostatic set points, allostatic demands and learned valuations without constituting consciousness themselves, and their influence can vary across states. We call this *organismic-affective anchoring* (but see [37]).

Dissociations between selfhood and awareness offer one way to assess whether organismic-affective anchoring is fixed or contextually weighted. Accounts grounding subjectivity in bodily regulation often treat this regulation as the basis of elementary selfhood, most explicitly in Damasio’s proto-self and core self [24, 25, 38]. Selfhood, however, spans embodied, perspectival, agentive, autobiographical, narrative and socially reflective processes [38–40]. For tractability, we group these deliberately coarsely into embodied and narrative forms [39, 41], the former roughly spanning bodily, perspectival and agentive processes and the latter autobiographical, narrative and socially reflective processes. Out-of-body experiences, dreaming, psychedelic self-loss, advanced contemplative practice and *minimal phenomenal experience* suggest that bodily self-location, ordinary bodily engagement, self-boundaries or narrative identity may attenuate or dissociate while awareness persists [41–48]. These cases do not show that physiological regulation disappears or that brainstem or insular systems become inactive. They instead suggest that organismic-affective anchoring supports different combinations of embodied and narrative self-related processes across conscious states rather than one invariant package. This is the pattern predicted by a distributed, context-sensitive account, on which the weighting of bodily, affective and narrative processes should vary with the state and content of experience rather than recur as a fixed self-related core.

Beyond subjectivity and selfhood, it remains difficult to evaluate whether consciousness depends on privileged territories or a context-dependent distributed architecture. Most consciousness experiments use one paradigm and modality, usually vision, so theory, method and anatomy can covary [49]. Few follow participants across visual, bodily and affective experiences to ask whether correlates remain invariant or shift with context, while supramodal work remains early and concentrated on perception [50]. Even the adversarial comparison spanned laboratories and imaging modalities but remained constrained within suprathreshold visual perception [14]. It did not test whether any territory remains privileged across conscious contents and states.

Reported effects may also include attention, report, decision, memory, motor preparation, or other prerequisites and consequences of experience despite efforts to control them [51, 52]. Because many recur across paradigms, they should increase cross-domain overlap. A privileged neural core of consciousness should likewise recur across domains. Common overlap would therefore remain ambiguous, but limited commonality would weigh against both an invariant core and dominance by one generic set of task-related processes. Testing this requires cross-domain synthesis, yet existing meta-analyses remain partitioned by domain, including visual awareness [53, 54], disorders of consciousness [55], hypnosis [56], meditation [57], psychedelics [58], bodily self-consciousness [34] and self-referential judgement [59, 60]. These syntheses identify reproducible systems within their domains but do not ask whether the wider literature converges on an invariant core.

Crucially, a distributed alternative to an invariant core does not imply indiscriminate whole-brain activation. A *distributed* account predicts selective convergence within particular contexts and broad anatomical participation when contexts are considered together. It also predicts limited exact overlap, because different contents and states recruit different combinations of specialised systems. We use *decentred* to mean that no coarse candidate locus adequately accounts for the literature as a whole, while allowing for local specialisation, causal gatekeepers and high-centrality regions. *Context-sensitive* means that neural organisation varies across research contexts rather than taking one fixed form.

Anatomical distribution is only one signature of a distributed, context-sensitive architecture. Such an architecture should also vary in the functional composition of convergent maps and the multi-region configurations associated with each domain. An invariant-core account may allow domain-specific additions but predicts a common functional and regional component across domains. A context-sensitive account instead predicts systematic variation in relative functional weightings and multi-region coalitions. Aggregating across contexts should then produce a composite that no single context reproduces, rather than reveal a hidden common core.

We tested a distributed, decentred and context-sensitive account of consciousness through seven predictions arising from the considerations above. These span anatomical, functional and network levels. We evaluated them in 579 coordinate-based neuroimaging studies whose titles or abstracts treated consciousness, awareness, conscious content or access, or changes in conscious state as substantive scientific targets. These studies were grouped into semantic domains, data-driven clusters of related consciousness literatures that served as the research contexts for all analyses.

At the anatomical level, the first prediction is selective voxelwise convergence but broad distribution across systems when semantic domains are considered together. This distinguishes a distributed architecture from indiscriminate wholebrain involvement and a compact privileged territory. Accordingly, when all domains are combined, the implicated voxels should span a larger anatomical territory than an equally sized set of voxels concentrated within a single compact region.

The second prediction is reproducible semantic domain-specific convergence with only partial overlap, because different contexts recruit different combinations of specialised regions. The corpus should also resist compression into one dominant spatial component expressing a common pattern across studies.

At the functional level, the third prediction is that the consciousness landscape should retain interpretable structure beyond generic neuroimaging reporting topography. Prespecified maps spanning executive-control (E), mnemonic/conceptual-social (M), perceptual/sensorimotor (S) and organismic-affective (O) functions should reconstruct the residual landscape better than spatially autocorrelated surrogates. Coarse regional predictors representing theory-associated loci, like the posterior ‘hot zone’, should add little beyond these functional maps.

The fourth prediction is that functional weighting should vary across semantic domains. Organismic-affective maps test organism-relative contributions, while embodied and narrative self-related proxies test whether aspects of selfhood make partly separable contributions.

On a decentred, context-sensitive account, consciousness-related information in brain maps should be distributed across functional families rather than concentrated in one of them. The fifth prediction therefore asks whether this information is redundant, when different functional families independently carry overlapping information about the consciousness landscape, or synergistic, when information about the landscape emerges only from considering multiple families jointly. We expected both, with greater joint information for the observed landscape than for matched reference landscapes, but no prediction about which would predominate [61, 62].

At the network level, the sixth prediction is that corrected consciousness territories should be embedded in broad coactivation networks rather than function as isolated loci.

Broad network embedding alone would still be compatible with one invariant architecture. The seventh prediction is therefore that complete multi-region coalitions should vary across domains rather than form one fixed configuration.

## 2 Results

### 2.1 Studies formed seven semantic domains

We constructed a broad coordinate-based functional neuroimaging corpus from NeuroStore and Neurosynth Compose [63]. A title-and-abstract search was followed by a two-tier screen in which three large language models independently assessed each title and abstract. The screen retained studies judged directly informative about consciousness and suitable for coordinate meta-analysis. Random 10% samples from both screening tiers were inspected as face-validity checks. The resulting corpus comprised 579 studies, 1,252 contrasts and 17,700 foci (Supplementary Table 1). Across the corpus, the most frequent eligible terms in titles and abstracts centred on ‘cortex’ and ‘consciousness’ (Fig. 1b).

**Figure 1.**
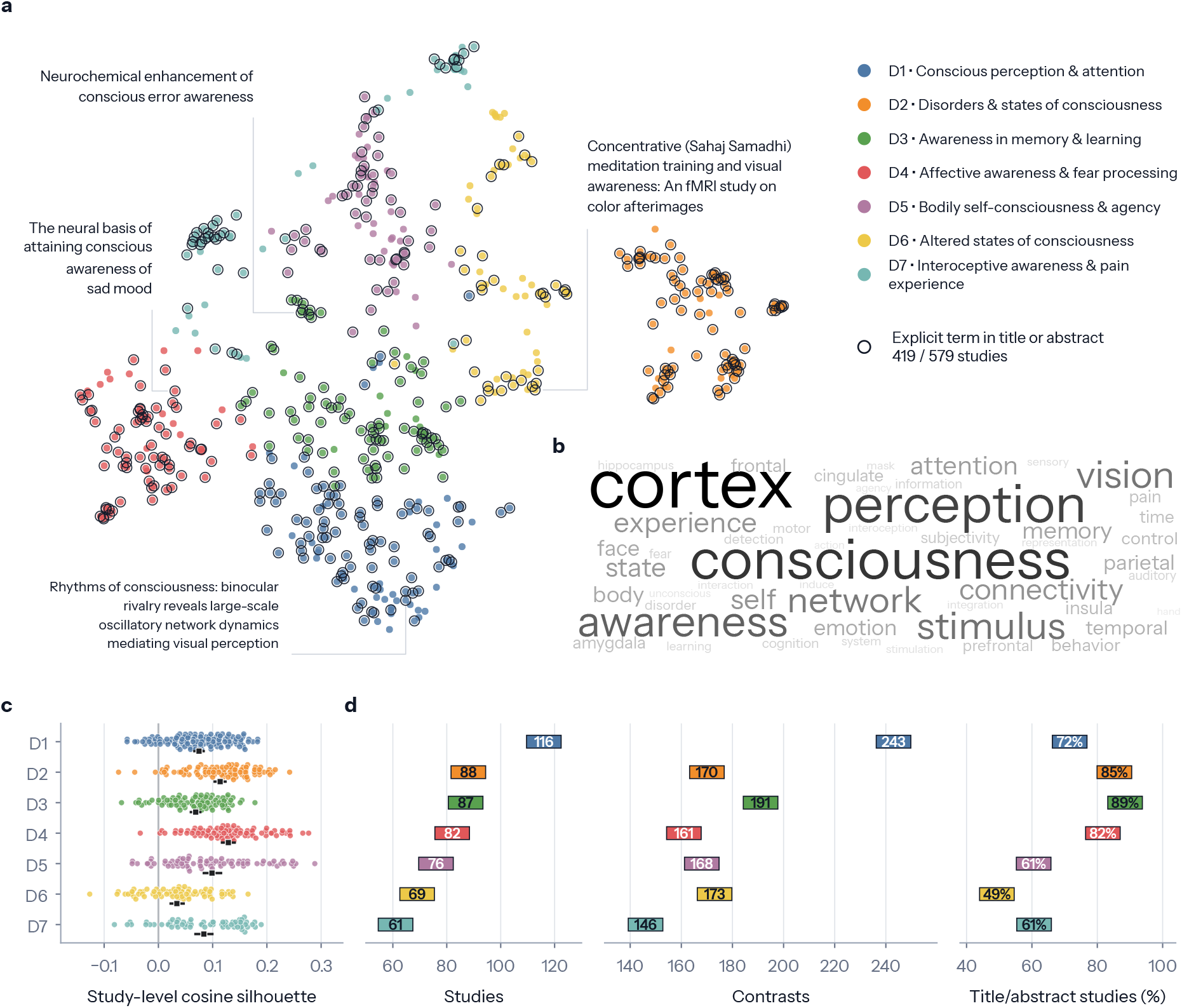
Semantic organisation and composition of the consciousness literature. **a**, Two-dimensional UMAP display of title and abstract embeddings from the 579-study corpus, visualising the semantic organisation of the literature. Each point is one study, colour identifies domain, and nearby points indicate greater semantic similarity in the projection. The four linked titles are illustrative, not cluster centroids or prevalence estimates. **b**, Fifty most frequent eligible terms across titles and abstracts, summarising the lexical content of the corpus; word size indicates corpus frequency. **c**, Study-level cosine silhouettes assessing how sharply the semantic domains separate in the original embedding space. Dots show individual studies, black squares domain means and horizontal lines 95% bootstrap intervals. **d**, Domain-level study counts, contrast counts and percentages of studies containing a direct consciousness or awareness term in the title or abstract, summarising domain composition and evidence volume. D1–D7 follow the labels in **a** and carry the same meaning in Figures 3, 4 and 6.

To test whether consciousness-related brain maps varied with research context, we clustered studies by the semantic similarity of their titles and abstracts without imposing categories in advance. Clustering stability analyses supported a seven-domain solution spanning a broad range of consciousness-related behaviours and states (Fig. 1a). Study-level cosine silhouettes in the original embedding space showed only weak separation (Fig. 1c), consistent with the domains’ shared focus on consciousness. Across domains, 49–89% of studies contained a direct consciousness or awareness term in the title or abstract (Fig. 1d). The results below therefore concern related consciousness literatures rather than unrelated topics, and the domains provide a useful scaffold rather than a unique taxonomy (Supplementary Note 1 and Supplementary Tables 1 and 2).

### 2.2 Descriptive landscape was anatomically and functionally broad

Before testing corrected convergence, we visualised the unthresholded whole-corpus ALE map as a descriptive summary of where reported foci were concentrated across the literature. It extended across much of the brain (Fig. 2a; Supplementary Note 2 and Supplementary Table 3).

**Figure 2.**
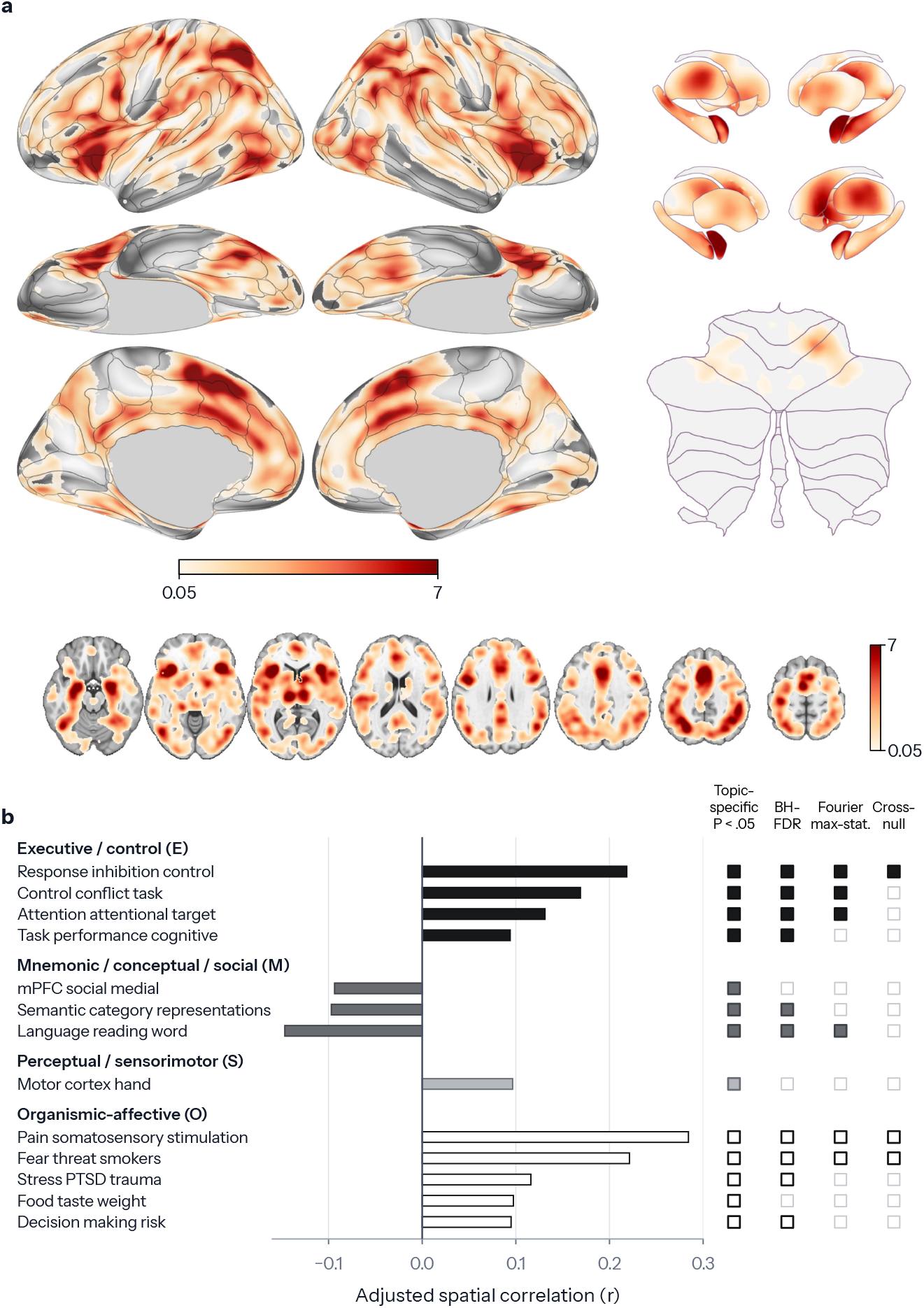
The descriptive consciousness landscape is anatomically and functionally broad. **a**, Continuous unthresholded whole-corpus activation likelihood estimation map from 579 studies, 1,252 reported contrasts and 17,700 foci, summarising the overall anatomical landscape. **b**, Reporting-density-adjusted spatial correlations for 13 associations that passed the topic-specific spatial test, assessing whether this landscape retains interpretable functional structure beyond generic reporting topography. Associations are grouped into the four broad functional families used in later analyses. Bars show Pearson *r*. Complete results for all 50 topics, including the undisplayed consciousness-adjacent topic, are reported in Supplementary Note 3 and Supplementary Table 4.

Because this breadth could partly reflect where neuroimaging studies tend to report effects, we asked whether the landscape retained interpretable functional structure after accounting for generic reporting topography. We compared its spatial pattern with 50 meta-analytic functional maps from NeuroSynth [64], after excluding source studies that overlapped with the target corpus or contained direct consciousness or awareness title terms.

Before reporting-density adjustment, all 50 topics survived family-wise correction. After adjustment, 25 passed topic-specific spatial tests, 16 survived FDR, 10 survived the primary family-wise correction and 4 survived both spatial nulls. Fig. 2b shows 13 associations that passed the topic-specific spatial test, spanning the four functional families used in later analyses, namely executive-control, mnemonic/conceptual-social, perceptual/sensorimotor and organismic-affective functions. Complete 50-topic results are reported in Supplementary Note 3 and Supplementary Table 4.

### 2.3 Convergence was selective yet anatomically distributed

We next meta-analysed each semantic domain separately using activation likelihood estimation and corrected the complete seven-domain-by-voxel family jointly. Taken together, the corrected maps spanned cortical, subcortical and cerebellar territories distributed widely across the brain (Fig. 3a). Because ALE localisation-kernel width varies with sample size and source sample sizes were unavailable, we used a common assumed *N* = 20. Varying the assumed *N* from 10 to 80 left the seven-map architecture broadly stable (Supplementary Note 4 and Supplementary Table 5).

**Figure 3.**
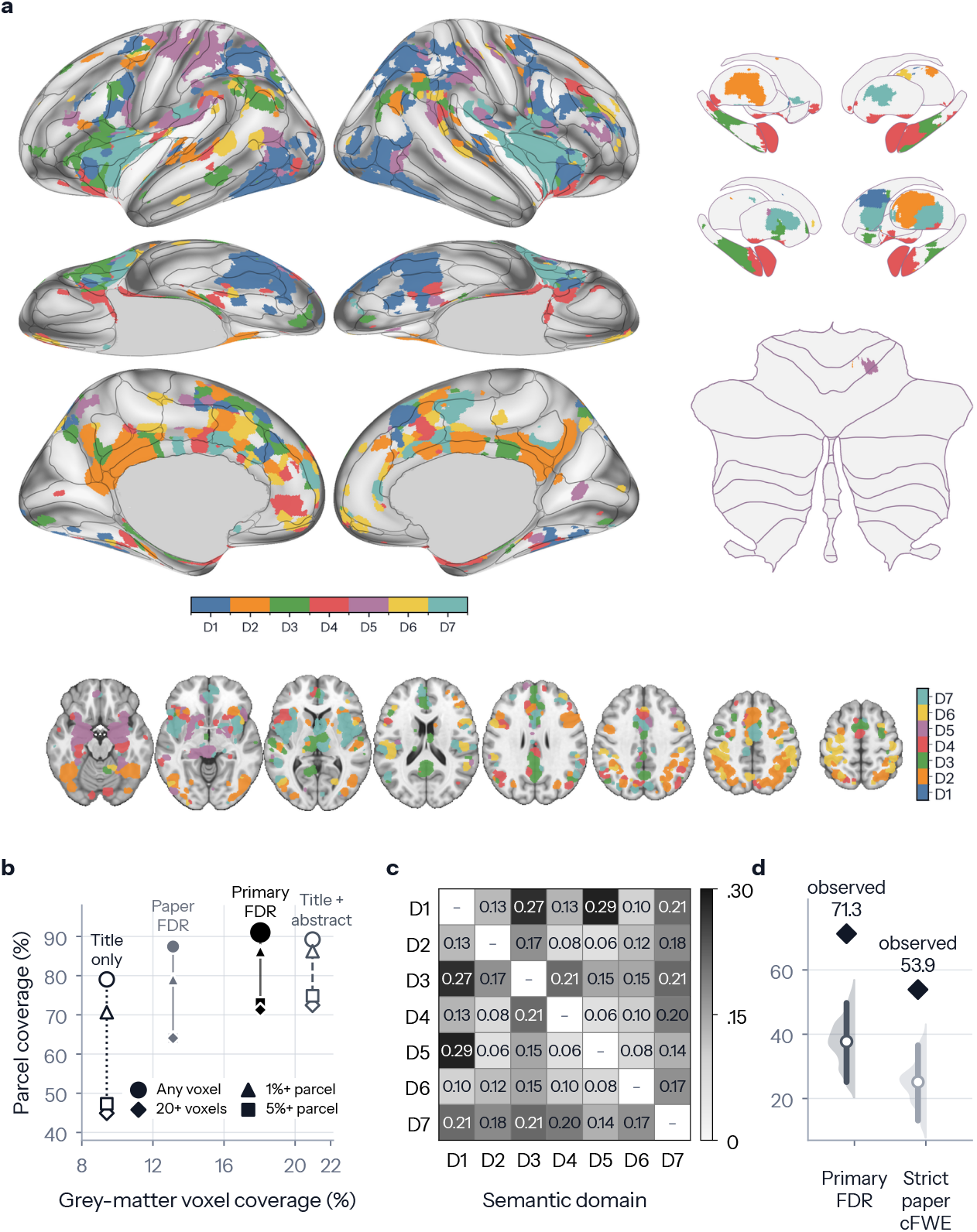
Corrected convergence is voxel-selective, anatomically widespread and differentiated across semantic domains. **a**, Corrected semantic-domain ALE maps, jointly controlled with FDR at *q <* .05, showing the spatial distribution of convergent territory across domains. Colour identifies domain rather than effect magnitude; where maps overlap, display colour follows the domain with the largest ALE value, and cross-domain overlap itself is shown in Figure 4. **b**, Grey-matter and parcel coverage across four seven-domain inference profiles and two whole-corpus lexical sensitivities, one restricted to studies containing direct consciousness or awareness terms in the title or abstract and the other to studies containing them in the title alone, contrasting local voxelwise extent with broader anatomical spread. Exact values are reported in Table 1. **c**, Pairwise Dice overlap among the primary corrected domain maps, quantifying shared corrected territory between domains. **d**, Parcel spread of the primary and strict corrected unions relative to 5,000 compact territories of identical voxel mass, testing whether broad parcel coverage follows from map size alone. D1–D7 follow the domain labels in Figure 1a.

To quantify anatomical distributedness, we compared the extent of corrected convergence in grey matter with its reach across atlas parcels. The seven-domain union occupied 18.0% of the grey-matter mask yet reached 91.0% of atlas parcels (Fig. 3b and Table 1). Convergence was therefore selective at the voxel level but broadly distributed anatomically. Individual domain maps were also broad, covering a median of 50.3% of parcels (range 44.3%-59.9%; Supplementary Note 5).

**Table 1.**
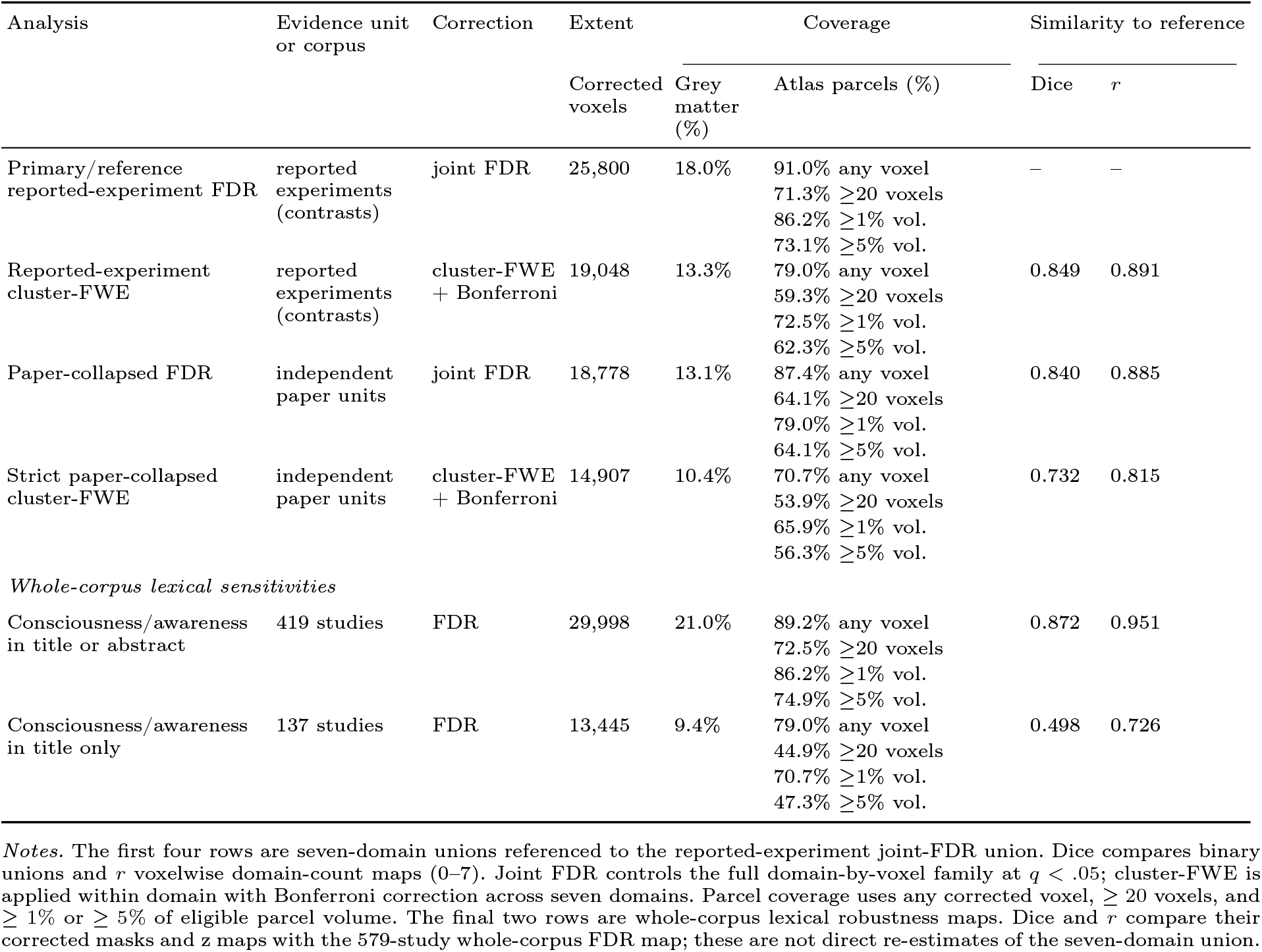
Corrected convergence across inference profiles and lexical sensitivities.

We tested this pattern across correction schemes, parcel-occupancy thresholds and lexical corpus restrictions. In every case, corrected evidence remained selective in grey matter while reaching most atlas parcels (Fig. 3b and Table 1). Parcel profiles were highly reproducible across random study divisions, and broad coverage survived removal of each domain in turn (Supplementary Notes 6 and 7, Supplementary Figure 1 and Supplementary Tables 6 and 7).

Broad parcel coverage could nevertheless reflect map size rather than spatial distribution. We therefore compared each observed union with 5,000 compact territories of identical voxel mass. Both the primary and stricter unions reached more parcels than their compact controls (Fig. 3d). A reporting-density-weighted randomised-coordinate control likewise showed a modest excess of parcel dispersion. The observed spread was therefore not reducible to one compact territory, though these controls do not exclude multifocal or network-localised accounts (Supplementary Note 8 and Supplementary Table 8).

A final control asked whether this degree of spread was specific to consciousness. It provided no evidence that consciousness was unusually parcel-distributed relative to matched heterogeneous cognitive-neuroimaging literatures. These controls support distributedness within the consciousness corpus without showing that it is unique to consciousness (Supplementary Note 9 and Supplementary Table 9).

### 2.4 Domain maps were differentiated without an invariant core

We next asked whether the distributed domain maps nevertheless converged on a common anatomical core. A total of 956 voxels were shared by at least four domains, but only 19 by all seven (Fig. 4a,b). Overlap across four or more domains involved insular and medial frontal/cingulate territories, with smaller thalamic involvement, before contracting to two nine-voxel components and one unassigned voxel (Supplementary Note 10).

**Figure 4.**
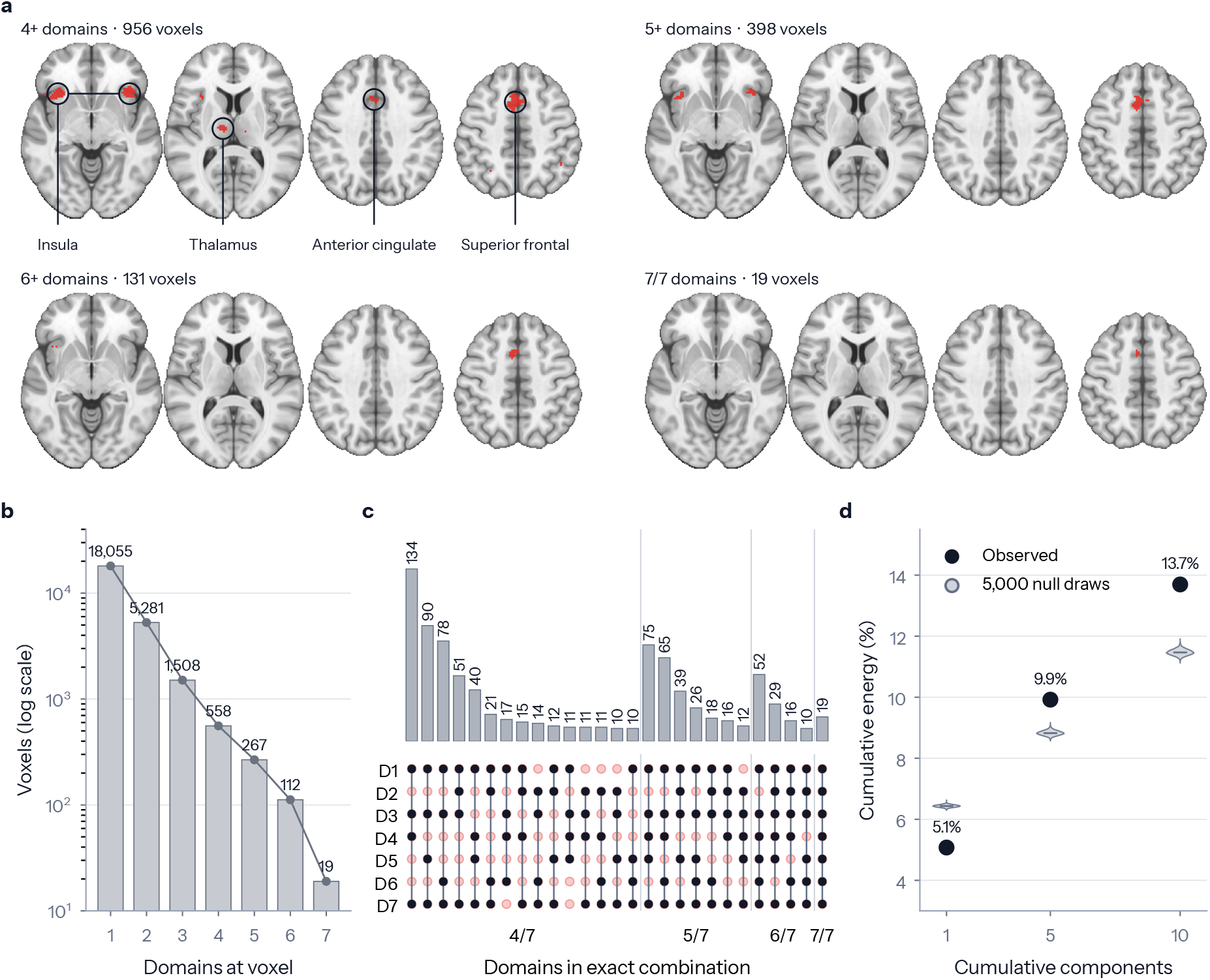
High-order overlap contracts sharply and contains changing domain combinations. **a**, Cumulative overlap across at least four through all seven corrected domain maps, showing shared territory at increasing levels of cross-domain overlap. **b**, Numbers of voxels present in exactly one through seven maps, summarising the distribution of domain membership across the corrected union. **c**, Exact four-through seven-domain combinations containing at least ten voxels, assessing whether shared territory reflects a stable subset of domains. Filled points identify included domains, pale open points omitted domains and bars show voxel counts. **d**, Cumulative activation energy from singular-value decomposition at *k* = 1, 5 and 10 components, compared with 5,000 sparse-map null draws to assess whether the corpus is dominated by a common spatial pattern. D1–D7 follow the domain labels in Figure 1a.

The 19-voxel intersection remained below the meaningful-core reference but was not unusually small under study-label permutation. Broader overlap involved changing combinations of domains (Fig. 4c), while the strict intersection did not recur at a fixed spatial location and recurrent overlap followed generic high-reporting-density territory. Thus, these shared loci are not interpretable as a consciousness-specific core (Supplementary Notes 11 and 12, and Supplementary Tables 10 and 11).

Sparse corrected overlap could still conceal one diffuse pattern expressed across studies. We therefore decomposed study-level activation maps with singular-value decomposition. The first component accounted for 5.1% of activation energy, compared with 9.9% and 13.7% across the first five and ten components (Fig. 4d). Null maps concentrated more energy in the leading component, whereas the observed top-five and top-ten totals were higher than expected; a reporting-density null gave the same pattern. Thus, neither thresholded overlap nor continuous study maps revealed one dominant spatial core.

However, the absence of an invariant core does not by itself show that domain maps vary systematically with context, because arbitrary partitions of a heterogeneous literature may also produce different maps. We therefore tested whether the semantic-domain maps were more differentiated than same-sized study-label partitions. Mean pairwise spatial overlap was 0.153, lower in the observed maps than in the partitions (*P* = 0.000999; Fig. 3c, Supplementary Note 13 and Supplementary Table 12). The domains were therefore collectively more differentiated than arbitrary partitions of the same literature. Changing four-through seven-domain combinations provided a descriptive counterpart to this differentiation (Fig. 4c).

Coordinate reporting volume per paper did not explain these domain differences. Complementary map-based split-half refits also retained the domains’ collective map identities, although the altered-states domain was less reliable (Supplementary Notes 14 and 6, Supplementary Figure 1 and Supplementary Tables 13 and 6).

### 2.5 Functional maps reconstructed the consciousness landscape

We next asked whether distributed functional maps could reconstruct the continuous consciousness landscape beyond generic neuroimaging reporting topography. Successful out-of-sample prediction would show systematic alignment with combinations of familiar functions rather than a spatial pattern unique to consciousness. Using spatially blocked cross-validation, we fitted ridge models with 28 NeuroSynth maps constructed after excluding target-corpus studies and grouped a priori into four theory-guided families. These spanned executive-control (E), mnemonic/conceptual-social (M), perceptual/sensorimotor (S) and organismic-affective (O) functions, together termed EMSO (Supplementary Table 14).

Reporting density alone explained total heldout *R*^2^= 0.7320. Adding the complete set of functional maps increased total performance to *R*^2^= 0.7830, an absolute increment of Δ*R*^2^= 0.0510. Of the variance remaining after foldwise density adjustment, EMSO explained residual *R*^2^= 0.1902 (95% spatial-block bootstrap interval 0.1400–0.2373, Fig. 5a,c).

**Figure 5.**
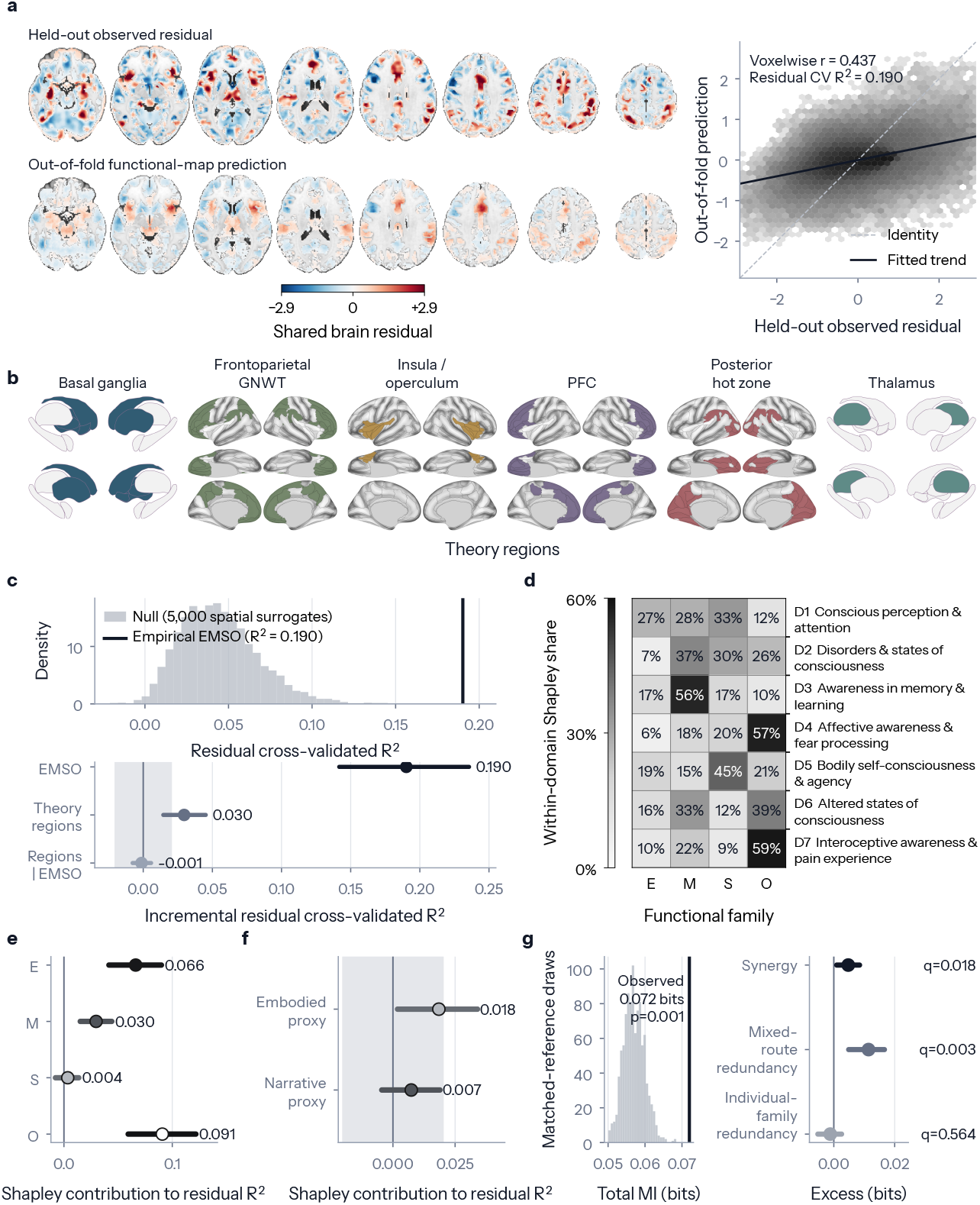
Distributed functional maps reconstruct the consciousness landscape and vary across contexts. **a**, Held-out reporting-density-adjusted target and out-of-fold functional-map prediction, assessing reconstruction of the residual consciousness landscape. The inset compares observed and predicted voxel values. **b**, Six theory-associated regional loci used to test whether theoretically privileged territories add prediction beyond distributed functional maps. The fitted block also included brainstem. **c**, Upper, residual cross-validated *R*^2^for the observed target relative to 5,000 spatial surrogates, testing target specificity. Lower, incremental residual *R*^2^for functional maps and regional predictors, comparing their conditional contributions. **d**, Descriptive within-domain Shapley shares, assessing contextual variation in functional weighting. **e**, Whole-corpus Shapley allocations, comparing conditional contributions of the four functional families. **f**, Secondary Shapley allocations for embodied and narrative self-related proxies, assessing partly separable contributions. **g**, Left, total mutual information relative to matched reference targets, testing whether the families jointly carry above-reference information. Right, excess synergy, mixed-route redundancy and individual-family redundancy, characterising how that information is shared. Complete estimates are in Supplementary Tables 14 and 16.

To test target specificity, we applied the same model to 5,000 spatial surrogates with approximately matched autocorrelation [65]. No surrogate matched the observed residual *R*^2^= 0.190 (empirical *P* = 0.0002; Fig. 5c). The fixed functional maps therefore reconstructed the observed landscape better than spatially comparable surrogates, without implying that EMSO is uniquely optimal.

To determine which families drove reconstruction, an exact Shapley decomposition averaged each family’s marginal contribution across all 16 combinations. Organismic-affective made the largest conditional contribution, followed by executive-control and mnemonic/conceptual-social functions. The perceptual/sensorimotor contribution was smaller and its interval crossed zero (Fig. 5e and Supplementary Table 14).

Because these families were correlated, their Shapley values represent conditional allocations rather than independent effects. Organismic-affective was less collinear with the other maps. This lower collinearity may have contributed to, but did not determine, its leading allocation (Supplementary Note 15 and Supplementary Table 15).

Finally, we asked whether the functional maps merely recapitulated seven theory-associated anatomical territories, like a posterior ‘hot zone’ (Fig. 5b and Supplementary Note 16). Fitted alone, these regional predictors explained *R*^2^ = *−*0.0296 beyond density. When added after EMSO, the regional predictors produced an increment of Δ*R*^2^= −0.0012 (95% interval −0.0077– 0.0057), entirely within the prespecified equivalence bounds (Fig. 5c). Thus, they carried limited predictive information alone but none that was practically meaningful beyond the functional maps.

### 2.6 Functional contributions varied across contexts

To ask whether the same functional composition recurred across consciousness contexts, we repeated the prediction and decomposition for each semantic domain (Fig. 5d). The profiles did not reproduce one common composition. M was largest in the disorders/states and memory/learning domains, S in perception/attention and bodily self/agency, and O in affective/fear, altered states and interoception/pain. E was not the largest family in any domain.

Because these domain-level profiles were descriptive, we tested their differences formally at the paper level. Four-dimensional functional profiles from 577 independent paper or sample units differed across the seven domains (pseudo-*F* = 17.10, multivariate *R*^2^= 0.153, *P <* 0.0001), without evidence that the result reflected unequal within-domain dispersion.

At the whole-corpus level, the organismic-affective family made the largest conditional contribution, consistent with organismic-affective anchoring within a wider architecture rather than a privileged organismic source (Fig. 5e). It comprised maps related to reward, bodily need, threat, affect, stress, valuation and pain, functions that are organism-relative but not necessarily explicitly self-referential. We next sought converging evidence for a context-sensitive rather than invariant organismic/self-related architecture by asking whether broad embodied and narrative self-related proxies contributed as one package or were partly separable. Neither analysis directly measures organismic significance or selfhood, but their convergence tests whether organism-relative, embodied and narrative contributions are differently weighted and partly separable.

A secondary partition compared seven narrative maps (speech perception, language, memory, semantic representation, autobiographical scene construction and social cognition) with seven embodied maps (bodily need, motor/action, body-space, affect, pain and multisensory processing). After removing all 14 from their parent EMSO families, the embodied proxy made a modest, more reliable contribution (Δ*R*^2^= 0.0185, 95% interval 0.0018–0.0340), whereas the narrative proxy was smaller and its interval included zero (Δ*R*^2^= 0.0074, −0.0045–0.0189; Fig. 5f and Supplementary Table 14). The embodied proxy partly overlapped organismic-affective functions but extended beyond them. Because both sets were removed before restoration, this was not an orthogonal three-way comparison, but showed only that the embodied proxy contributed more reliably than the narrative proxy beyond the same remaining maps.

### 2.7 Information was both redundant and synergistic

The continuous EMSO reconstruction showed that the four functional families jointly predicted the residual landscape, but not whether their information was redundant, synergistic or both. We therefore classified locations as above or below the landscape median and measured how much information each family, alone and in combination, carried about that classification. Together, they carried more information about the observed consciousness landscape than about matched meta-analytic reference landscapes (0.0720 versus 0.0570 bits; *P* = 0.000999; Fig. 5g). The excess of 0.0150 bits was modest, but greater than expected from the matched reference landscapes (Supplementary Note 17).

The decomposition showed excess synergy, information available only from combinations (0.0047 bits, *q* = 0.017982). When resolved by combination size, only three-family synergy showed positive excess over the references. Individual-family redundancy, where the same information was available separately from two or more individual families, did not exceed the references. By contrast, mixed-route redundancy, where the same information was available through alternative routes including at least one multi-family route, was elevated (0.0114 bits, *q* = 0.002997; Fig. 5g). Thus, the functional maps carried synergistic information alongside redundant routes to the same target (Supplementary Table 16).

After conditioning on the other three families, organismic-affective and mnemonic/conceptual-social maps retained additional above-reference information (Fig. 5g and Supplementary Table 16). These conditioned increments are not pure measures of unique information because they can also contain synergy involving the focal family. They therefore provide upper bounds on unique information.

### 2.8 Focal territories shared a broad network repertoire

Corrected convergence identifies where foci recur, but not the broader systems with which those territories are repeatedly reported. We therefore used each domain’s corrected territory as a seed for corpus-internal meta-analytic coactivation modelling, identifying regions repeatedly co-reported with the seed across experiments.

The seven seeds selected 441–640 contrasts each, and their corrected networks covered 81.4%-87.4% of atlas parcels (Supplementary Data). Despite sparse overlap among focal convergence territories (Fig. 3c and Fig. 4a–c), the seven network maps shared a broad anatomical repertoire (Fig. 6a).

**Figure 6.**
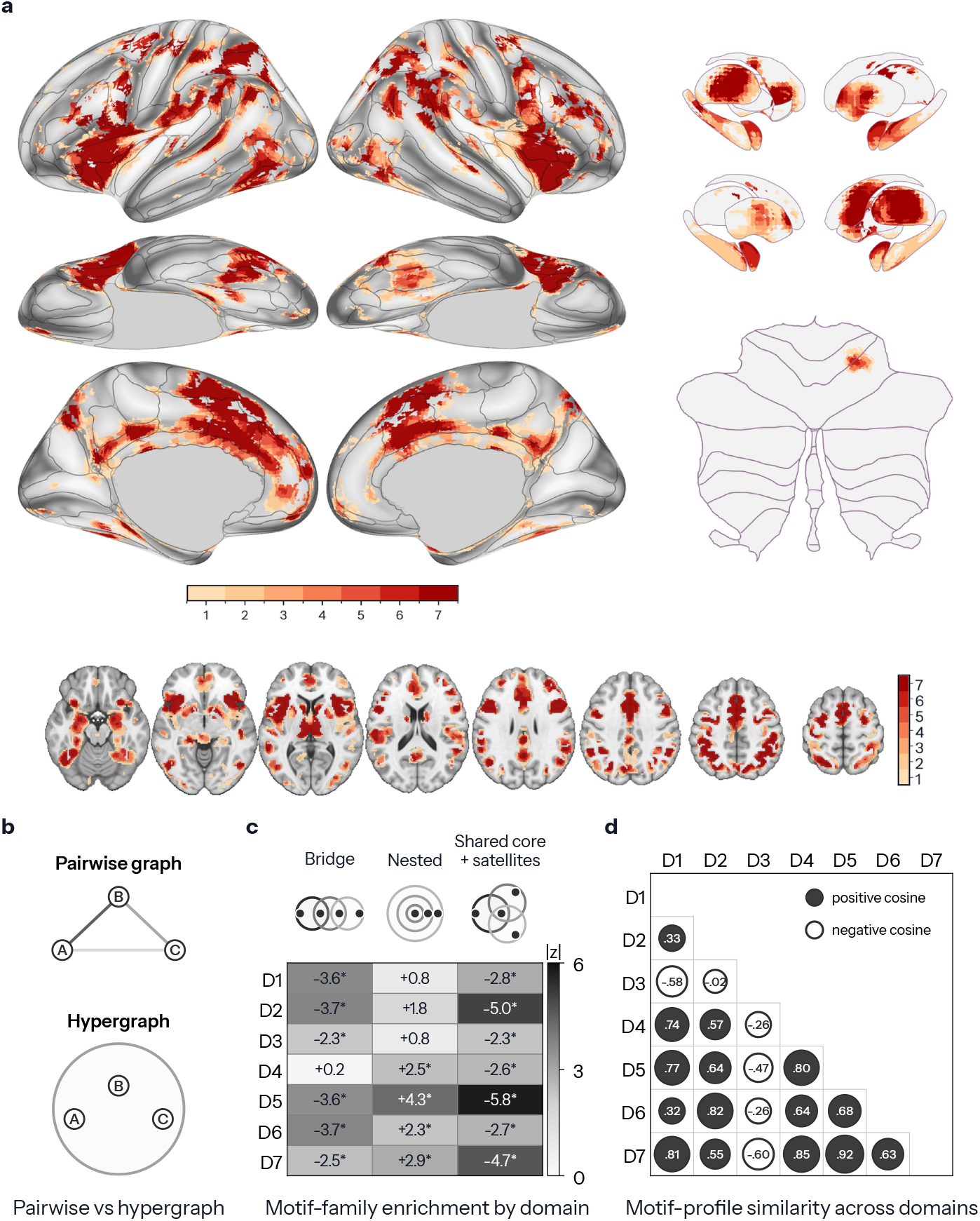
A broadly shared network repertoire is organised into context-dependent higher-order coalitions. **a**, Overlap across seven corrected corpus-internal meta-analytic coactivation networks, each seeded by a domain’s corrected convergence territory, assessing whether focal territories share a broader network repertoire. Colour indicates the number of domain networks containing each voxel. **b**, Schematic contrasting pairwise co-reporting with a hyperedge preserving complete within-contrast coalitions. **c**, Signed enrichment *z* scores by domain relative to matched randomised hypergraphs for the three motif families with corrected effects. Replicated coalitions had none and are omitted, but remained in the 28-test correction. Asterisks mark BH-FDR *q <* .05. **d**, Pairwise cosine similarities among complete 26-motif profiles, providing a descriptive comparison of detailed coalition structure across domains. D1–D7 follow the domain labels in Figure 1a.

### 2.9 Coalition structure varied across domains

The shared coactivation repertoire did not establish one invariant network configuration because seed-based maps do not preserve which regions occur together within individual contrasts. A distributed architecture predicts that such coalitions should recur non-randomly, while context sensitivity predicts that their form should vary across domains. To test this, we represented each contrast as a hyperedge linking all reported parcels [66] (Fig. 6b). We counted 26 recurring relationships among triplets of hyperedges and compared each domain with 5,000 randomised hypergraphs preserving how often each parcel appeared and the number of parcels per contrast. Sixteen domain-by-motif effects survived family-wise correction, and 17 of 28 domain-by-family tests survived FDR correction, spanning three of the four structural forms (Supplementary Data).

Relative to these nulls, bridge chains, where one coalition links two otherwise separate coalitions, were depleted in six domains. Nested expansions, where one coalition is nested within a broader one and extended by a third, were enriched in four domains. Shared cores with satellites, which contain a common core and private extensions, were depleted in all seven domains (Fig. 6c). These terms describe deviations from the null rather than absolute prevalence. No individual motif met the cross-domain conservation criterion, and conservation was not tested at the broader family level. Broad structural tendencies nevertheless recurred, while the complete 26-motif profiles varied across domains (mean pairwise cosine similarity 0.376, Fig. 6d). Thus, a broadly shared regional repertoire was organised into different higher-order configurations across semantic domains (Fig. 6c,d).

Separately, we mapped parcel roles within motifs, classifying parcels as core, intermediate/bridge or satellite. These descriptive roles varied across domains after adjustment for parcel reporting frequency (Supplementary Figure 2, panels a–g). These maps are descriptive and do not establish that any parcel preferentially occupied a role (Supplementary Note 18).

## 3 Discussion

Across anatomical, functional and network levels, this synthesis revealed a distributed, decentred and context-sensitive neurobiology of consciousness built from functionally specialised systems. Seven related semantic domains provided the contexts for these findings (Fig. 1). Anatomically, convergence was locally selective yet widespread and resisted reduction to a compact territory or dominant spatial pattern (Figs. 2–4 and Table 1). Functionally, distributed maps reconstructed the landscape beyond reporting density and better than spatial surrogates, while loci associated with consciousness theories added no practically meaningful prediction and functional contributions varied across domains (Fig. 5a–f). The maps also carried redundant and synergistic information (Fig. 5g). At the network level, distinct focal territories were embedded in a broadly shared repertoire whose higher-order coalitions varied across domains (Fig. 6a,c,d). Overall, the evidence supports neither a privileged locus nor undifferentiated whole-brain activation, but rather specialised systems weighted and combined differently across contexts.

Spatial controls qualified the distributedness claim. The observed landscape was more parcel-distributed than compact or reportingdensity-weighted randomised-coordinate fields, but matched heterogeneous neuroimaging literatures showed comparable spread (Supplementary Note 8 and Supplementary Note 9). The principal conclusion is therefore not that consciousness engages more of the brain than other complex domains, but that consciousness-related evidence is composed differently across perceptual, mnemonic, affective, bodily, interoceptive, altered-state and disorders-of-consciousness contexts.

Beyond distributedness and sparse cross-domain overlap, we showed that organisation changed systematically with context. Differentiation exceeded arbitrary partitions, while higher-order domain combinations, functional profiles and higher-order coalitions varied across domains (Fig. 3c, Fig. 4c, Fig. 5d and Fig. 6c,d). Because the semantic domains were only weakly separable, these differences are better understood as graded shifts among related literatures rather than categorical kinds (Fig. 1c and Supplementary Note 1).

Several findings bear more directly on existing accounts of consciousness, beginning with the distinction between local causal importance and anatomical privilege. The seven candidate territories for leading theories of consciousness carried limited predictive information alone and added no practically meaningful information beyond the distributed functional maps (Fig. 5b,c and Supplementary Note 16). This challenges the sufficiency of these territories, and of the theories that privilege them, as general accounts of the neurobiology of consciousness, not their relevance. Perturbational and lesion-network evidence supports the same distinction. Central-thalamic stimulation can restore distributed cortical organisation during anaesthesia, human thalamic stimulation associated with recovery engages a network disrupted across other consciousness-impairing conditions, and cortical lesions causing loss of consciousness map to a distributed brainstem-linked circuit [67–69]. A region can therefore exert substantial causal influence without constituting a privileged substrate for consciousness.

Organismic-affective functions add a more specific constraint. Within the correlated functional maps, those associated with bodily need, affect, threat, reward, stress, valuation and pain received the largest conditional allocation (Fig. 5e). Two results nevertheless distinguish this leading contribution from a core. It was not fixed across contexts. Organismic-affective maps led only in the affective/fear, altered-states and interoception/pain domains, mnemonic or perceptual maps led elsewhere, and paper-level functional profiles differed across domains (Fig. 5d). Anatomically, the territory most often shared across domains, spanning anterior insula, medial frontal/cingulate cortex and, at lower thresholds, thalamus, is compatible with organismic-affective involvement, but it followed generic high-reporting-density territory, did not recur at a fixed location and nearly vanished in the strict seven-way intersection (Fig. 4a–c).

This is consistent with organismic-affective anchoring as a context-sensitive contribution within a wider architecture rather than a privileged source of consciousness. It therefore retains a central insight, shared by subjective-frame, interoceptive-inference, affect-centred and proto-self accounts, that conscious processing is situated relative to the condition and value of the organism, while relaxing the requirement that consciousness originate in a privileged organismic system [25, 26, 28, 70]. The anchor is thus a frequently, though not invariably, recruited frame of organism-relative significance rather than a foundation on which every conscious state is built.

Together, the organismic-affective ranking and self-related proxy analysis suggest that organismic-affective, embodied and narrative contributions need not covary as one invariant package. This interpretation accords with multidimensional accounts of selfhood and with states in which dimensions of selfhood attenuate while awareness persists, as during psychedelic or meditative self-loss [41, 44–46, 48]. Recent 5-MeO-DMT, nondual meditation and jhāna (deep meditative absorption) studies pair profound alterations of selfhood with large-scale reorganisation of distributed brain dynamics, providing tractable models of how the functional architecture of consciousness changes as ordinary selfhood attenuates [71–75].

The target-information analysis showed excess synergy and mixed-route redundancy across functional families relative to matched reference landscapes (Fig. 5g). This combination parallels information-decomposition work in which synergy supports integration and emergent information available only jointly, whereas redundancy supports robustness, maintenance and distributed sharing of the same information [61, 62]. Although our decomposition concerns spatial prediction rather than neural information flow, it suggests a compatible architecture in which specialised contributions are both combined and available through overlapping routes.

The network analyses revealed a similar balance between shared architecture and contextual reconfiguration. A broadly shared coactivation repertoire was organised into different higher-order coalitions across domains (Fig. 6c,d). This parallels network-neuroscience findings that diverse cognitive states largely preserve a common network architecture while selectively reconfiguring its interactions, including hypergraph evidence for both task-general and task-specific higher-order organisation [76, 77]. Consciousness-related contexts may likewise differ less in the systems available than in how a shared repertoire is assembled.

Several limitations qualify these interpretations. First, study inclusion was decided from titles and abstracts by three-model voting. The first author inspected a random 10% of records at each screening tier, but the complete corpus, ambiguous records and contrast provenance were not adjudicated from full texts. This trade-off is common in large-scale automated neuroimaging synthesis [64]. Second, although the original studies often attempted to control attention, report, decision, memory, motor preparation or related processes, reported coordinates cannot determine whether the associated activity was constitutive of experience or reflected prerequisites and consequences that remained despite those controls [51, 52]. However, to the extent that the same prerequisites or consequences recur across paradigms, they should increase cross-domain commonality rather than produce the lower-than-expected overlap, differing functional profiles and varying coalition structures observed across domains. Third, the semantic domains compare different literatures rather than contexts manipulated within the same participants. Population, modality, task, source measurement, sample size, preprocessing and smoothing were not completely classified (Supplementary Note 14 and Supplementary Table 13). Methodological and theory-linked choices may shape domain effects [49], but they do not explain away the selective yet broad convergence within every domain. Population and paradigm are themselves part of the consciousness contexts compared here, but their independent contributions cannot be separated in this design.

Individual analyses carry additional interpretive limits. The functional and self-related analyses use prespecified, theory-guided partitions of correlated maps. These groupings are informative, but they are not a definitive taxonomy of brain function and do not directly measure organismic significance or selfhood. The regional predictors are likewise coarse operationalisations rather than complete implementations of the theories they approximate. The target-information analysis describes static spatial relations among fixed templates and a meta-analytic target. Meta-analytic coactivation describes co-reporting across experiments, whereas hypergraphs preserve co-reporting within contrasts. Neither measures connectivity or causal interaction within brains. The present study therefore addresses the architecture represented across the neuroimaging literature, not the temporal or causal mechanism of an individual conscious episode.

These limitations aside, our results strongly motivate a distributed, decentred and context-sensitive coalition account. The constituent systems need not be unique to consciousness. Instead, the content, character and organism-relative significance of a conscious episode depend on how systems supporting perception, cognition, action and organismic regulation are coordinated into a context-specific coalition. The coalition is not a fixed consciousness network. Its composition should change predictably with what is experienced, how the organism is engaged and how consciousness is experimentally probed [49]. Interoceptive challenge should increase organismic-affective weighting. Report and metacognitive demands should increase executive-control and mnemonic/conceptual-social weighting. Visually guided flow should increase perceptual and action weighting while reducing narrative weighting. Self-loss or minimal phenomenal experience should reduce narrative or ordinary embodied weighting without abolishing experience. Local perturbations should have different consequences depending on which coalition they reorganise. The observed mixture of redundancy and synergy suggests that overlapping routes may stabilise some aspects of a state, while synergistic combinations may support structure unavailable to any one system.

Multiple-drafts and dynamic-core theories are the closest existing matches to our proposal because they reject a fixed theatre and allow organisation to vary across contents and states [15–18, 20]. Temporo-spatial theory is broadly aligned in emphasising spatially nested organisation rather than a fixed substrate [19]. Integrated information theory likewise requires differentiated yet integrated activity, but the present account does not require an invariant posterior substrate [5, 6]. Recurrent processing and predictive coding offer mechanisms for coordinating content, expectations and organismic significance [7, 28]. Global workspace broadcasting and higher-order representation may support access, report and reflective awareness when present rather than defining every conscious state [3, 4, 8]. Thalamic, brainstem, insular and cellular mechanisms may gate coalition formation without serving as universal generators. A single regional and functional configuration that remained sufficient across carefully matched contexts would count against the account.

These findings shift the central question from where consciousness is to how specialised systems are weighted and assembled across contents, states and forms of selfhood. The neurobiology of consciousness lies not in an invariant core but in the distributed coalitions through which conscious states take form.

## 4 Methods

### 4.1 Corpus construction

We assembled the corpus from a local NeuroStore/Neurosynth Compose database containing 32,411 base studies associated with reported stereotactic coordinates [63]. We deliberately searched broadly and imposed specificity during screening. A wide-net title-and-abstract search covered explicit consciousness terminology, established awareness paradigms, global and altered states, and bodily or self-consciousness. Neither coordinate count nor the number of reported foci influenced study inclusion. Publications were deduplicated by DOI, then PMID, then NeuroStore base-study identifier. The complete search expression, deduplication procedure and screening inputs are provided in the Supplementary Information and analysis code.

The two screening tiers answered different questions. Tier I established whether a study was substantively about consciousness. OpenAI GPT5.5, Claude Opus 4.8 and Gemini 3.1 Pro Preview independently assessed each title and abstract. A study qualified only when consciousness, awareness, conscious content or access, subjective experience, or a change in conscious state was itself the object of the scientific question, comparison, manipulation or finding. Merely studying participants who were conscious during scanning was not sufficient. Eligible routes included explicit studies of consciousness or awareness, direct manipulations of perceptual awareness or conscious access, global-state studies involving anaesthesia, sleep or disorders of consciousness, altered-state studies involving psychedelics, hypnosis or meditation when tied to subjective experience, and bodily or self-consciousness. Masking, rivalry, attentional-blink, blindsight, threshold-perception and no-report paradigms qualified only when the abstract explicitly related the relevant comparison to awareness. This criterion distinguished evidence specifically brought to bear on consciousness from the generic neural activity reported when a conscious participant performs a task. A study advanced only when all three models returned `include = yes`.

Tier II was run on the 1,094 records supported by at least two Tier I models and established whether an otherwise relevant study supplied functional coordinates suitable for coordinate-based meta-analysis. A study was excluded when the available coordinates came exclusively from structural or anatomical analyses, lesion or lesion-symptom mapping, a single case or small case series, or secondary rather than original empirical reports. Functional PET and fMRI results were eligible, including activation, resting-state, connectivity, component and graph analyses; mixed studies were retained when they contained at least one eligible functional result. Unclear cases were retained. A study was excluded at Tier II when at least two models returned `methods_exclude = yes`. Final corpus inclusion required unanimous Tier I support and fewer than two Tier II exclusion votes. Full prompts, decision fields, supporting evidence, model settings and vote tables are provided with the repository.

The wide-net search yielded 3,388 candidates. Tier I yielded 1,094 records supported by at least two models, including 695 with unanimous support. Tier II excluded 116 records within the unanimous set, leaving 579 studies. The distribution of inclusion routes and agreement between models is reported in the Supplementary Information. A small number of retained records lacked the representation required for particular downstream analyses and were excluded only from those analyses.

The first author also inspected a randomly selected 10% of Tier I records and a separate randomly selected 10% of Tier II records. These informal face-validity checks found the classifications sufficiently accurate to continue. They did not alter the screening rules, and no separate coded validation dataset or agreement statistics were retained.

### 4.2 Semantic embedding, stability and labels

To organise the literature by research context without imposing categories, we represented each study by its title and abstract. Text was embedded with BAAI/bge-large-en-v1.5 [78], yielding a 1,024-dimensional vector normalised to unit length.

UMAP reduced the embeddings to ten dimensions for clustering, while a separate two-dimensional projection was used only for visualisation [79]. A nearest-neighbour graph was built in the ten-dimensional space and partitioned with Leiden community detection [80]. Full UMAP and Leiden settings are provided in Supplementary Methods.

Because both UMAP and Leiden are parameter-dependent, we evaluated stability across UMAP seeds 13, 42 and 73, neighbourhood sizes 15, 30 and 50, and Leiden resolutions 0.5, 1.0 and 1.5. We also clustered a 30-nearest-neighbour graph built directly from the original embeddings, bypassing UMAP. Agreement among solutions was assessed with the adjusted Rand index and normalised mutual information. Seven and eleven clusters tied as the modal counts, so the lower-count rule selected seven. Among seven-cluster solutions, the partition with the highest mean adjusted Rand agreement across all stored candidates used seed 42, 50 neighbours and resolution 1.5. This was the scaffold used in subsequent analyses (Supplementary Table 1).

To assess how sharply the selected domains separated, we analysed the original embedding space rather than the two-dimensional display. We quantified within-domain dispersion, distances between domain centroids, nearest-centroid margins and cosine silhouettes, with percentile intervals from 2,000 cluster-stratified bootstrap samples. These descriptive results are shown in Fig. 1c and reported in Supplementary Note 1 and Supplementary Table 2.

Clusters were fixed before they were named. For each cluster, term frequency–inverse document frequency summaries and illustrative titles in fixed study-identifier order were used to generate independent label proposals from the same three model families. A single adjudicating model (OpenAI GPT-5.5), presented with the three proposals in anonymised and shuffled order so that their source could not be inferred, selected or minimally combined the proposals, with no human override. The final labels are shown in Fig. 1a.

### 4.3 Functional decoding of the descriptive landscape

To ask whether the descriptive consciousness landscape had interpretable functional structure, we compared it with the 50 maps from the NeuroSynth v0.7 topic model [64]. Each topic map combined 10-mm multilevel kernel density maps from the source studies, weighted by how strongly each study loaded on that topic [35]. The source model contained 14,371 studies.

To reduce circularity, we reconstructed these maps after removing 303 source studies that overlapped with the target corpus and 106 whose titles directly referred to consciousness or awareness. After overlap between these exclusions, 14,035 studies remained. Matching rules and the complete exclusion lists are provided in Supplementary Methods.

The target was the continuous, unthresholded whole-corpus ALE map. We calculated its voxelwise Pearson correlation with each topic map across the shared whole-brain mask. Because neighbouring voxels are spatially dependent, significance was evaluated against 10,000 phase-randomised versions of the target that preserved its spatial power spectrum while disrupting anatomical alignment. The largest absolute topic correlation in each draw provided two-sided family-wise error control across the full topic set. Topic-specific spatial probabilities, FDR across topics, Bonferroni correction and the maximum-statistic result were retained as graded evidence tiers.

As a robustness analysis, we repeated the reporting-density-adjusted 50-topic comparison with 10,000 volumetric BrainSMASH target surrogates and the same two-sided max-*T* correction [65]. Because these surrogates were substantially smoother than the observed target, making this null conservative for correlations with fixed smooth topic maps, only associations surviving both nulls were classified as robust across null construction; Fourier-only associations were left unresolved rather than rejected (Supplementary Note 3 and Supplementary Table 4).

A further control asked whether these associations reflected functional similarity or merely the locations in which neuroimaging studies most often report effects. We constructed a reporting-density map from all coordinate-bearing studies in NeuroStore, removed its spatial pattern from both the target and topic maps and recomputed their correlations. The resulting associations describe functional correspondence beyond generic reporting topography. The same reporting-density map was used as the nuisance baseline in the predictive analyses below.

### 4.4 Coordinate meta-analysis using ALE

To estimate where reported coordinates converged across studies, we used activation likelihood estimation (ALE) in NiMARE [81–83]. Analyses used the NiMARE MNI152 2-mm grid. MNI coordinates were retained and Talairach coordinates were converted to MNI space. Coordinates without a recognised space label were treated as MNI in the primary analysis and omitted in a sensitivity analysis. The analysis mask combined the NiMARE mask with MNI152 grey-matter tissue maps and contained 142,985 voxels; exact coordinate-handling and tissue-threshold rules are provided in Supplementary Methods.

ALE represents each focus as a spatial probability distribution and estimates convergence across experiments. No numeric source sample sizes were available in the production metadata, so all contrasts used the configured value *N* = 20. This held localisation uncertainty constant rather than representing actual enrolment.

To test sensitivity to this common-kernel assumption, we repeated the whole-corpus map and both reported-experiment and paper-collapsed seven-domain profiles with common assumed sample sizes of *N* = 10, 20, 40 and 80, spanning FWHM kernels of 8.63–10.00 mm. A same-kernel *N* = 20 repeat estimated Monte Carlo variability, and strict cluster-FWE refits at *N* = 10 and *N* = 80 tested the extremes (Supplementary Note 4 and Supplementary Table 5).

Each semantic domain was meta-analysed separately, with every eligible coordinate-bearing contrast treated as one experiment. Because the primary claim concerned convergence across the complete set of domains, all voxels in all seven maps were treated as one inferential family. Benjamini–Hochberg false-discovery-rate correction was applied at *q <* .05 before the corrected maps were combined. A separate unthresholded whole-corpus ALE map served as the descriptive landscape. As a stricter sensitivity, we also applied cluster-level family-wise error correction within each domain (cluster-forming *p <* .001; 10,000 iterations), followed by Bonferroni correction across the seven maps.

Because some papers contributed several eligible contrasts, both correction schemes were repeated after pooling all eligible coordinates within paper and removing duplicates. This produced 579 independent paper units from 1,252 contrasts and 17,700 foci, retaining 16,711 unique coordinates. A publication-level assessment asked whether domains differed simply in how much coordinate information each paper contributed, using the numbers of retained foci, source contrasts and removed duplicates (Supplementary Note 14).

To distinguish a common core from context-specific convergence, we counted how many corrected domain maps contained each voxel and calculated pairwise Dice and Jaccard overlap. The strict seven-way intersection was assessed in two ways. First, its extent was compared with a prespecified meaningful-core bound equal to one tenth of the smallest corrected domain map (298 voxels). Second, 1,000 study-label permutations preserved domain sizes, refitted all seven maps and repeated the joint correction to provide a reference for the intersection’s extent (Supplementary Note 11). The same permutations provided the reference for mean pairwise Dice; pair-specific and continuous-map analyses are reported in Supplementary Note 13.

Finally, five whole-corpus sensitivity refits tested the effects of removing the three non-human studies, removing contrasts with uncertain coordinate-space labels, applying validated coordinate-provenance corrections, restricting the corpus to studies with explicit consciousness or awareness terminology in the title or abstract, and restricting it to studies with such terminology in the title alone. Each sensitivity used the same grid and mask and was corrected at *q <* .05. Because these refits corrected one whole-corpus map rather than the seven domain maps jointly, their coverage estimates were treated as robustness checks rather than direct re-estimates of the primary union (Supplementary Table 1 and Table 1).

### 4.5 Parcel spread and spatial controls

To quantify anatomical spread, we mapped the corrected ALE results to the grey-matter-filtered FreeSurfer Destrieux aparc.a2009s plus aseg atlas in MNI space [84]. Intersecting the supplied atlas with the common analysis mask retained 167 parcels, comprising 148 cortical, 17 subcortical and two cerebellar-cortex parcels. The atlas labelled 67.0% of the analysis mask. Parcel analyses used only labelled in-mask voxels and did not restrict voxelwise ALE analyses. We assessed coverage using four criteria, namely any corrected voxel, at least 20 corrected voxels, and at least 1% or 5% of the parcel’s eligible volume. The proportional criteria tested whether broad coverage depended on isolated edge voxels or unequal parcel size (Supplementary Note 5). For matched-mass controls, we also calculated normalised Shannon entropy, with higher values indicating a more even distribution across parcels.

A large map can cross many parcels simply because of its size. We therefore asked whether one compact territory containing the same number of voxels could reproduce the observed spread. For each corrected union, 5,000 null draws generated a single compact, 26-connected territory of identical voxel mass within the analysis mask. The primary statistic was the number of parcels containing at least 20 voxels. Upper-tail probabilities used the plus-one rule [85]. This prespecified compact-locus test was applied to both the primary reported-experiment joint-FDR union and the stricter paper-collapsed cluster-FWE union.

#### Randomised-coordinate control

The compact-locus control does not reproduce the smoothness or multifocal structure produced by ALE. As a complementary test, we relocated all 17,700 foci according to whole-NeuroStore reporting density while preserving studies, experiments, focus counts and semantic-domain assignments. Each of 1,000 draws then reran the seven domain-specific ALE analyses. We compared normalised parcel entropy after selecting 16,675 atlas-assigned voxels from every field, matching the atlas support of the observed reported-experiment joint-FDR union rather than its complete voxel extent. A lower-mass sensitivity selected the strongest 14,000 atlas voxels, all lying within each draw’s joint-FDR union. Moran’s *I* assessed residual differences in smoothness. Complete spatial-fit diagnostics and sensitivity analyses are reported in Supplementary Note 8.

#### Matched-literature control

To ask whether this degree of distribution was distinctive of consciousness, we compared the paper-collapsed consciousness map with 1,000 samples from the wider NeuroSynth literature. Samples were matched exactly to the target on the number of papers and the complete distribution of foci per paper, after excluding records that overlapped with the target or contained direct consciousness or awareness title terms. Every experiment received the same assumed sample size, *N* = 20, so that ALE kernel width was identical across the target and controls. All maps underwent the same ALE, FDR and matched-mass procedures, and normalised parcel entropy was the primary statistic. Because only four primary controls contained at least 21,083 FDR-significant atlas voxels, we repeated the equal-mass comparison at *K* = 16, 000, which lay below the FDR extent of every target and control map. Every voxel selected in this sensitivity therefore survived FDR. Focus-matched topic literatures were retained as descriptive comparisons (Supplementary Note 9).

### 4.6 Spatial decomposition of study maps

Broad parcel coverage could still arise from one diffuse spatial pattern expressed weakly across many studies. To test this possibility, we constructed one modelled-activation map for each of the 576 studies with at least one in-mask focus. Each map was smoothed with a 10-mm Gaussian kernel and normalised to unit length so that studies reporting more foci did not dominate the analysis. The maps were assembled into a study-by-voxel matrix.

Singular-value decomposition ranked recurring spatial patterns by the share of activation energy they explained. We report the energy captured by the first component and cumulatively by the first five and ten components. A dominant common pattern would concentrate energy in the first component, whereas several recurring patterns would distribute it more broadly.

To determine whether this component structure exceeded that expected from sparse coordinate maps, we generated 5,000 null matrices. Each study retained its number of foci, smoothing and normalisation, but focus locations were resampled from the spatial density of the consciousness corpus. A sensitivity analysis instead sampled from whole-NeuroStore reporting density. The same decomposition was applied to every null matrix. Because a single-source account predicts greater concentration in the leading component, the first-component statistic was tested in the lower tail; the cumulative five- and ten-component statistics were tested two-sided.

### 4.7 Reliability, domain influence and coordinate validation

Two split-half analyses asked whether the broad anatomical profile and the semantic-domain maps were reproducible across independent subsets of studies. First, the corpus was divided randomly 1,000 times. Within each half, reported foci were assigned to atlas parcels, using the nearest valid parcel within 4 mm when necessary, and summed to form parcel-count profiles. Pearson correlations between the two profiles measured the reproducibility of the corpus-wide anatomical distribution.

Second, we created 100 complementary partitions within each semantic domain and reran the complete seven-domain ALE analysis in both halves. We compared every map in one half with every map in the other. The confirmatory statistic was the mean correlation between maps bearing the same domain label, evaluated against all 5,040 permutations of the labels in the second half. This tested whether the domain maps retained their collective identities across non-overlapping study samples. Thresholded-map, parcel, peak and identification measures were descriptive (Supplementary Note 6).

We also asked whether one exceptional domain concealed a large common core. Each domain was omitted in turn and the six-map intersection was compared with position-matched exclusions across 1,000 study-label permutations. Remaining-six union coverage was reported descriptively to show whether broad spread persisted after each exclusion, but it was not part of the inferential family (Supplementary Note 7 and Supplementary Table 7).

Finally, we documented coordinate-space labels and all analysis-specific representation exclusions. Contrasts that could not generate a valid modelled-activation map or parcel hyperedge were excluded only from analyses requiring that representation, not from the study-selection corpus.

### 4.8 Functional-map reconstruction and context dependence

#### Functional maps and reconstruction

We asked whether the continuous consciousness landscape could be reconstructed from a distributed set of familiar functions after generic neuroimaging reporting topography was removed. The target was the unthresholded whole-corpus ALE map in the shared 142,985-voxel mask. Predictors were 28 NeuroSynth topic maps selected before model fitting from the 50-topic set estimated after the source-study exclusions described above.

The maps were organised into four equal, theory-guided families informed by RDoC [86]. Executive-control (E) included working memory, task performance, inhibition, conflict and attention. Mnemonic/conceptual-social (M) included social cognition, memory retrieval, language, semantics, numerical cognition and autobiographical scene construction. Perceptual/sensorimotor (S) included auditory and speech perception, motor and action processes, body and spatial representation, and visual and multisensory functions. Organismic-affective (O) included reward, bodily need, threat, affect, stress, valuation and pain. Together, these families are termed EMSO. Each contained seven maps, listed in Supplementary Table 14d. All 28 maps entered the regression individually; family labels were used only to compare model subsets and allocate predictive contributions. Equal family size prevented the number of maps from mechanically favouring one family.

Twenty-one topics that did not describe a coherent psychological function and one consciousness-adjacent topic were excluded before fitting. The grouping served as an interpretable functional model rather than an official RdoC ontology. Supplementary Table 4 reports the complete inclusion and exclusion status of all 50 topics.

Models used ridge regression because the topic maps were correlated and regularisation stabilised their joint out-of-sample prediction [87]. Five-fold spatial-block cross-validation limited leakage between neighbouring voxels [88]. The 2-mm grid was divided into contiguous 8 *×* 8 *×* 8-voxel blocks, approximately 16-mm cubes, and assigned to five folds. The same folds were used for every model, and ridge penalties were selected in nested spatial folds. Within each fold, reporting density was fitted on training voxels and removed from the target and predictors before prediction in heldout voxels. Centring and scaling were likewise estimated from training data only.

We report total held-out *R*^2^for the raw target and residual held-out *R*^2^, the primary measure, after reporting-density adjustment. We fitted the density-only model and all 16 combinations of E, M, S and O. Exact Shapley values [89] averaged each family’s added predictive value across every subset and entry order.

Because the topic maps were correlated, we also quantified correlations among family-average templates and how well each family’s maps could be reconstructed from the remaining 21 predictors. Paper-level correlation diagnostics are reported in Supplementary Note 15. These analyses contextualised the Shapley allocations rather than providing independent contribution estimates.

Uncertainty was estimated from 5,000 spatial-block bootstrap samples. Residual Δ*R*^2^= 0.020 was prespecified as the smallest meaningful predictive increment.

#### Spatial target specificity

Cross-validation alone does not establish that the functional-map set is specific to the observed landscape, because smooth predictors can fit other smooth brain maps. We therefore generated 5,000 BrainSMASH surrogate targets with approximately matched spatial autocorrelation [65]. The observed target was separated into reporting-density and residual components, and BrainSMASH randomised the residual field.

For each surrogate, any remaining association with reporting density was removed, its mean and variance were matched to the observed residual, and the fixed reporting-density component was restored. Surrogates were then evaluated with the same mask, folds, nested tuning and 28 predictors as the observed map. The upper-tail empirical probability counted surrogates whose residual *R*^2^matched or exceeded the observed value, using the plus-one rule. Complete BrainSMASH settings, postprocessing and variogram diagnostics are provided in Supplementary Methods.

#### Context-dependent functional profiles

To ask whether functional composition varied across consciousness contexts, we first repeated the complete 16-subset reconstruction and Shapley analysis for each unthresholded semantic-domain ALE target. The resulting seven-by-four heat map was descriptive.

Formal inference used four family-average templates constructed separately from the individual predictors used above. Each eligible contrast received four spatial-similarity scores, which were averaged within paper or independent sample, yielding 577 paper-level profiles.

The four dimensions were standardised before a one-way Euclidean PERMANOVA [90] tested whether profiles differed across semantic domains. Domain labels were shuffled 10,000 times while preserving domain sizes. A companion dispersion test [91] assessed whether any effect reflected unequal within-domain spread, and the analysis was repeated without standardisation. This omnibus test concerned the four-dimensional profiles as a whole rather than individual heat-map cells or domain pairs.

#### Regional-locus comparison

We next asked whether the distributed set of functional maps merely recapitulated theory-associated anatomical territories. Seven binary predictors represented basal ganglia, brainstem, frontoparietal workspace, insula and operculum, prefrontal cortex, posterior hot zone and thalamus. The masks were defined a priori from the 167-label Destrieux/FreeSurfer atlas and entered jointly as one regional block. The associations were motivated by basal-ganglia circuit [92], upper-brainstem [10, 11], global workspace [4], interoceptive and insular [27, 93], higher-order prefrontal [8], posterior hot-zone [6, 13] and thalamic accounts [9, 21]. Regional-only, EMSO and combined models used the same target, reporting-density adjustment, folds, nested tuning and paired spatial-block bootstrap.

The primary statistic was the change in heldout residual *R*^2^when the regional block was added to the complete set of functional maps. Practical equivalence required the complete 95% paired-bootstrap interval to fall within [*−*0.020, 0.020]. Six regional loci are illustrated in Fig. 5b, and all seven binary masks, parcel definitions and overlap fields are supplied as Supplementary Data.

#### Self-related proxy analysis

Finally, a secondary partition asked whether embodied and narrative self-related functions contributed as one package or were partly separable. The narrative set contained seven maps related to speech and language, social cognition, memory, semantic representation and autobiographical scene construction. The embodied set contained seven maps related to bodily need, motor and action processes, body and spatial representation, affect, pain and multisensory processing.

These 14 maps were removed from their parent families to define the remaining maps, preventing any predictor from entering twice. We compared the remaining maps alone, with each proxy set separately and with both together using the same cross-validation, two-block Shapley decomposition and bootstrap. Complete topic membership is listed in Supplementary Table 14e.

### 4.9 Information sharing among functional families

The reconstruction analysis established how well the four functional families predicted the residual consciousness landscape, but not whether the information they carried was redundant, synergistic or both. We therefore treated the four fixed, reporting-density-adjusted EMSO family templates as sources and the similarly adjusted whole-corpus ALE map as the target across 142,985 grey-matter voxels. Following the empirical MMI-PID approach of Jansma and colleagues [94], each map was binarised at its median. Alternative high-state fractions of 40% and 60% were descriptive sensitivities.

The reference distribution comprised 1,000 target maps generated by relocating all 17,700 foci according to whole-NeuroStore reporting density while preserving studies, experiments, focus counts and the *N* = 20 ALE kernels. The source templates remained fixed, and the ALE and reporting-density-adjustment pipeline was rerun for every reference map. Before inference, the reference distribution had to pass prespecified spatial-quality gates relative to the observed target, namely mean six-neighbour same-state agreement within 5% and mean high- and low-state component counts within 50% (observed mismatches −0.81%, 8.28% and 15.57%). All 1,000 references passed and entered upper-tail plus-one inference.

To test how information was shared across the functional families, total mutual information served as an omnibus test of whether the joint family state carried more target information than the matched references. We then examined all non-empty source subsets, Shapley allocations, fully conditioned increments, concentration summaries and a four-source minimum-mutual-information partial information decomposition [94–96]. Synergy denoted information available only from combined families. Mixed-route redundancy denoted information available through two or more alternative routes, with at least one route combining families. Individual-family redundancy denoted information available separately from two or more individual families. The secondary decomposition tests used two-sided plus-one probabilities with Benjamini–Hochberg correction within each named family. Fully conditioned increments were interpreted as upper bounds on unique information because they also include synergy involving that family.

### 4.10 Meta-analytic coactivation networks

Corrected convergence identifies where foci recur, but not the broader systems in which those territories are reported. We therefore used meta-analytic coactivation modelling, or MACM, to estimate which regions were repeatedly co-reported with each seed across the literature [97, 98]. Each domain’s joint-FDR-corrected map served as a seed. Every contrast in the consciousness corpus containing at least one focus in that seed was selected, whole-brain ALE was rerun on the selected contrasts, and the resulting map was controlled with FDR at *q <* .05 within the common mask. Voxel and parcel coverage summarised the seven corpus-internal coactivation maps.

### 4.11 Context-dependent higher-order coalitions

Meta-analytic coactivation modelling shows which regions are repeatedly co-reported with a seed across experiments, but it does not preserve which regions occur together within an individual contrast. We therefore represented each coordinate-bearing contrast as a hyperedge linking all atlas parcels containing at least one focus. Foci were assigned directly to the atlas or to the nearest valid parcel within 4 mm; contrasts spanning fewer than two parcels were omitted. Exact duplicate parcel sets within a domain were counted once in the primary unweighted analysis. Complete assignment and dependence sensitivities are provided in Supplementary Methods.

We counted the 26 connected motifs formed by triplets of hyperedges defined by Lee, Ko and Shin [66]. For each semantic domain, 5,000 randomised hypergraphs preserved how often each parcel appeared and the number of parcels per contrast while randomising their co-occurrence. Motif counts were standardised against these nulls, and the maximum absolute statistic across motifs provided family-wise error control within each domain.

Before examining the results, motifs sharing broad topological features were grouped into bridge chains, shared cores with satellites, nested expansions and replicated coalitions. Domain-by-family tests were corrected together with Benjamini–Hochberg FDR. Paper-dependence sensitivities repeated the analysis after collapsing within paper and after repeatedly sampling one contrast per paper. Cross-domain conservation was assessed only for individual motifs and required a consistent direction and corrected evidence that survived these dependence controls. Descriptively, each motif’s deviation was transformed as 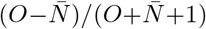, each domain’s resulting 26-element characteristic profile was L2-normalised, and all pairwise cosine similarities were calculated; no pairwise inference was performed (Fig. 6d).

As a descriptive anatomical follow-up, parcels were classified as core, intermediate/bridge or satellite according to their membership across the three hyperedges in each retained motif. Because these role frequencies were strongly related to how often a parcel was reported, role fractions were regressed on log hyperdegree and the residuals were mapped. Maps were displayed only above a minimum-support threshold and carried no parcel-level inference (Supplementary Figure 2, panels a–g, and Supplementary Note 18); complete maps and role summaries are supplied as Supplementary Data.

### 4.12 Use of artificial intelligence

Large language models were used for study screening and post-clustering label proposals as described above, and assisted with analysis coding, figure generation and manuscript drafting and editing. The authors made all scientific decisions and take responsibility for the analyses, interpretations and final manuscript. No language model was treated as an author.

### 4.13 Reproducibility

Analyses were configured in `config/publication.yaml` and run with Snakemake [99] through `workflow/Snakefile`. Machine-readable outputs populated manuscript values, maps, tables and figures. Versioned inputs, prompts, model outputs, commands, software environments and provenance records are provided with the public release, allowing the workflow to be reproduced without live language-model calls.

### 4.14 Data availability

Redistributable corpus metadata, classification decisions, statistical maps and source-data tables are available at https://github.com/jskipper/distributed-consciousness. Upstream datasets that cannot be redistributed are documented with retrieval instructions and identifiers.

### 4.15 Code availability

Analysis code, configuration files, prompts, archived model outputs and software environments are available at https://github.com/jskipper/distributed-consciousness.

## Acknowledgements

J.I.S. thanks Ursula Oliveira for her support, patience and enthusiasm for consciousness research.

## Author contributions

J.I.S. conceived the study, developed the methodology, curated the data, performed the analyses, prepared the figures and drafted the manuscript. C.T. contributed to conceptualisation, theoretical framing, interpretation and manuscript revision. R.M., K.J. and M.J. contributed to interpretation and manuscript revision. G.C. contributed to methodology, robustness and sensitivity analyses and revision of the title and manuscript. G.B. contributed to conceptualisation and methodology, including the information-theoretic analysis strategy, figure development and manuscript revision. All authors reviewed and approved the final manuscript.

