## Supplementary Information for "What is it like to be a blob? A distributed, decentred and context-sensitive neurobiology of consciousness"

Jeremy I Skipper[1,2*], Christopher Timmermann[1,2], Rosalind McAlpine[1,2,3], Krisztina Jedlovszky[1,2], Magdalena Jaglinska[1,2], Greg Cooper[1,3], and George Blackburne[1,2]

[1]UCL Centre for Consciousness Research, University College London, UK.
[2]Experimental Psychology, University College London, UK.
[3]Clinical, Educational and Health Psychology, University College London, UK.

#### Overview

The Supplementary Information provides additional methodological detail and results supporting the main analyses. Supplementary Methods follow the order of the main Methods. Numbered Supplementary Notes follow the order in which the corresponding analyses appear in the main Results. There are two Supplementary Figures and sixteen Supplementary Tables. Supplementary Data are supplied separately.

### Supplementary Methods

#### Corpus search and study screening

Candidate records were drawn from the local NeuroStore and Neurosynth Compose database used for the coordinate corpus. Retrieval was deliberately broad and used case-insensitive title-and-abstract matching across explicit consciousness terminology, perceptual-awareness paradigms, global and altered states, and bodily or self-consciousness. The complete expression and its source version are supplied with the analysis code. Publications were deduplicated by DOI, then PMID, then NeuroStore base-study identifier. Neither coordinate count nor the number of reported foci contributed to the screening decision.

Each candidate received an opaque study identifier before model screening. Tier I records contained only that identifier, title and abstract. Tier II records additionally contained available contrast labels. Original manuscript claims, expected outcomes and outputs from the other models were not included in the model-facing records. OpenAI GPT-5.5, Claude Opus 4.8 and Gemini 3.1 Pro Preview independently returned structured decisions, confidence ratings, exclusion reasons and supporting text. Provider prompts, structured-output schemas, settings and versioned provider outputs are supplied with the repository.

Tier I assessed whether consciousness was itself the scientific target. A record qualified only when consciousness, awareness, conscious content or access, subjective experience, or a change in conscious state formed part of the scientific question, manipulation, comparison, association or finding. Studies conducted in conscious participants were not eligible on that basis alone. Masking, rivalry, attentional blink, blindsight, threshold-perception and no-report designs qualified only when the abstract explicitly connected the relevant comparison to awareness. A record advanced to the final corpus only when all three models supported inclusion.

Tier II assessed whether an otherwise relevant record supplied coordinates suitable for functional coordinate-based meta-analysis. Structural or anatomical coordinates, lesion analyses, single cases or small case series, and secondary-source foci were exclusionary when at least two models agreed. Unclear cases and mixed studies containing at least one eligible functional result were retained. Final inclusion therefore required unanimous Tier I support and fewer than two Tier II exclusion votes.

The search produced 3,388 candidates. Tier I yielded 1,094 records supported by at least two models, including 695 with unanimous support. Tier II excluded 116 records from the unanimously supported set, leaving 579 studies. The retained relevance routes comprised 325 explicit-consciousness studies, 171 informative-difference studies and 83 state-informative studies. The first author independently inspected a random 10% of Tier I records and a separate random 10% of Tier II records. These informal face-validity checks found the classifications sufficiently accurate to continue and did not change either screening rule. The complete list of included study titles and semantic-domain assignments is supplied as Supplementary Data.

Analysis-specific exclusions were applied only when a downstream representation could not be formed. These included contrasts without valid coordinate maps or parcel hyperedges, three nonhuman studies removed in a human-only sensitivity, validated coordinate-version replacements and validated coordinate-provenance exclusions. None altered membership of the 579-study screening corpus.

#### Semantic embedding, stability and labels

Titles and abstracts were whitespace-normalised separately and combined in a fixed title-then-abstract format. Text was encoded in batches of 16 with BAAI/bge-large-en-v1.5 [1]. The resulting 1,024-dimensional float32 vectors were normalised to unit length. The exact matrix and ordered study index used in the analysis are supplied as Supplementary Data.

UMAP produced a ten-dimensional representation for clustering and a separate two-dimensional representation for display [2]. Both used cosine distance and a minimum distance of 0.1. The stability grid crossed seeds 13, 42 and 73, neighbourhood sizes 15, 30 and 50, and Leiden resolutions 0.5, 1.0 and

1.5. For each solution, an unweighted symmetrised nearest-neighbour graph was built in UMAP space and partitioned with the Leiden `RBConfigurationVertexPartition` quality function [3]. A direct 30-nearest-neighbour cosine graph was also constructed from the original normalised embeddings without UMAP.

Agreement among solutions was quantified with adjusted Rand index and normalised mutual information. Seven and eleven clusters each occurred in five grid solutions. The lower-count tie rule selected seven. Among seven-cluster candidates, the retained solution had the highest mean adjusted Rand agreement with all stored candidates, including the direct graph. It used seed 42, 50 neighbours and Leiden resolution 1.5. Final domain identifiers were ordered by decreasing size and then by the minimum fixed study-row index. The separate two-dimensional projection used the baseline construction with 30 neighbours and seed 42 solely for display.

The geometry of the retained domains was quantified in the original embedding space. We calculated within-domain distance to the normalised centroid, mean pairwise cosine distance, nearest-centroid margin, cosine silhouette and all 21 pairwise centroid distances. Percentile intervals used 2,000 cluster-stratified bootstrap samples with assignments held fixed; no inferential probabilities were calculated. Main Figure 1c displays the study-level cosine silhouettes, and Supplementary Note 1 and Supplementary Table 2 report the domain-level summaries.

Domains were labelled only after membership was fixed. Within each domain, TF-IDF used English and domain-specific stop words, unigrams and bigrams, a minimum document frequency of two and at most 8,000 features. The eight highest-scoring terms and eight illustrative titles in fixed study-identifier order were supplied to the three model families. A blinded adjudication selected or minimally combined their anonymous proposals. No human override was applied.

For main Figure 1a, open rings identify studies meeting the word-bounded title-or-abstract consciousness or awareness rule used for the lexical sensitivity; the four linked titles are illustrative examples. Figure 1b shows the 50 most frequent eligible terms across titles and abstracts after deterministic lemmatisation, declared concept mappings and general and domain-specific stop-word removal. Terms were weighted by corpus frequency and placed with a fixed seed. Figure 1c shows study-level cosine silhouettes from the original embedding space, and Figure 1d shows domain study counts, contrast counts and explicit-language percentages using fixed-width value markers. The complete term table, mappings, exclusion list and placement table are supplied as Supplementary Data.

#### NeuroSynth topic maps and functional decoding

Functional interpretation used the NeuroSynth v0.7 50-topic latent Dirichlet allocation model [4]. Each source study was converted to a 10-mm multilevel kernel density map and weighted by its loading on each topic [5]. Weighted study maps were summed to produce one continuous map per topic. Source studies matching the consciousness corpus by PMID, DOI or normalised title were removed. We also removed source studies whose titles contained direct consciousness or awareness terms. After accounting for overlap between these exclusions, 14,035 source studies remained.

The direct consciousness-adjacent topic remained in the descriptive 50-topic analysis but was excluded from the later functional-map reconstruction. This kept the descriptive screen complete while preventing a lexically circular predictor from entering the reconstruction model.

The target was the continuous unthresholded whole-corpus ALE map. Pearson correlations with each topic map were calculated across the common volumetric mask. Spatial dependence was evaluated with 10,000 three-dimensional Fourier phase-randomised targets that preserved the target power spectrum while disrupting anatomical alignment. Each draw contributed the largest absolute correlation across all 50 topics. This yielded a two-sided maximum-statistic family-wise reference. Topic-specific probabilities, Benjamini-Hochberg FDR, Bonferroni correction and Fourier maximum-statistic probabilities were retained as graded evidence tiers.

Generic neuroimaging reporting topography was estimated from all coordinate-bearing studies in the local NeuroStore database. Coordinates were placed on the common MNI grid and smoothed with the implementation's 1.5-voxel Gaussian parameter. The target and each topic map were separately regressed on this density field, and their residuals were correlated. The same density field served as the nuisance baseline in the predictive analyses.

An alternative spatial-null analysis repeated the reporting-density-adjusted comparison with 10,000 volumetric BrainSMASH target surrogates and the same two-sided maximum-statistic correction [6]. Each surrogate was residualised against reporting density, recentred and scaled to the observed residual target. Spatial fidelity was assessed in a separate set of 100 surrogates using variogram error and signed short-lag differences. The BrainSMASH fields were 17.13% smoother than the target at short lags and had a relative variogram RMSE of 0.3070. This mismatch is conservative for correlations with fixed smooth topic maps, so associations surviving only the Fourier family-wise test were treated as sensitive to null construction rather than rejected (Supplementary Note 3 and Supplementary Table 4).

Main Figure 2b displays all and only the 13 executive-control, mnemonic/conceptual-social, perceptual/sensorimotor and organismic-affective associations that passed the topic-specific spatial test, comprising four E, three M, one S and five O topics. The direct consciousness-adjacent topic remained in the complete 50-topic analysis but is not displayed. Complete results for all 50 topics are supplied as Supplementary Data. Descriptive anatomical follow-ups used 26-neighbour components and retained component extent, peak, corrected-z-weighted centre and parcel occupancy of the corrected whole-corpus map.

#### Coordinate normalisation, masks and ALE

Coordinate meta-analysis used NiMARE on the MNI152 2-mm grid [7]. MNI and MNI152 coordinates were retained. Talairach and TLRC coordinates were converted to MNI space by NiMARE. Unknown or missing coordinate-space labels were treated as MNI in the primary analysis and removed in the known-space sensitivity. The complete ledger contained 937 MNI contrasts with 13,640 foci, 188 Talairach contrasts with 1,935 converted foci and 127 contrasts with 2,125 foci treated as MNI.

The analysis mask intersected the NiMARE mask with MNI152 tissue maps. Voxels were retained when grey-matter probability was at least 0.2 and at least as large as white-matter probability. The resulting mask contained 142,985 voxels. The atlas-labelled support used for parcel analyses contained 95,846 voxels.

ALE represented each focus as a sample-size-dependent spatial probability distribution and estimated convergence across reported experiments [8, 9]. Numeric source sample sizes were unavailable, so every contrast used the common value $N = 20$. This imposed the same localisation-uncertainty kernel across contrasts rather than representing actual enrolment. A fixed-kernel sensitivity repeated the principal profiles with a common $N$ of 10, 20, 40 or 80 in each refit, alongside an independent same-kernel $N = 20$ Monte Carlo repeat using a different seed and strict cluster-FWE analyses at the two extremes (Supplementary Note 4 and Supplementary Table 5).

Each semantic domain was first analysed with one reported contrast as one experiment. The uncorrected probability maps for all seven domains were treated as one family. Benjamini-Hochberg correction was applied jointly across 1,599,381 domain-by-voxel tests at $q < .05$, after which the corrected maps were combined. A separate whole-corpus ALE map supplied the continuous descriptive landscape.

Because several papers contributed multiple contrasts, the joint-FDR and strict cluster-FWE analyses were repeated after pooling all eligible coordinates within paper. Coordinates were normalised, rounded to three decimal places and deduplicated before each paper was represented as one experiment. This produced 579 independent paper units from 1,252 source contrasts and 17,700 source foci, retaining 16,711 unique coordinates. Strict inference used a cluster-forming threshold of $p < .001$ and 10,000 Monte Carlo iterations within each domain, followed by Bonferroni correction across the seven maps.

Cross-domain overlap was summarised with the number of corrected maps containing each voxel, pairwise Dice and Jaccard coefficients and majority overlap defined as at least four of seven maps. The meaningful-core reference was one tenth of the smallest corrected domain map, rounded upward.

Study-label permutations preserved domain sizes, refitted all seven maps and repeated the joint correction. The strict intersection used lower-tail plus-one inference. The same permutations supplied references for mean pairwise Dice and continuous-map correlations. Pair-specific Dice deficits used position-standardised single-step maximum-statistic correction. Main Figure 4a–c shows cumulative, exact-count and exact-combination overlap; Supplementary Notes 10–13 report the anatomical and permutation analyses.

Main Figure 4c displays exact intersections involving four through seven domains when they contained at least ten voxels. This threshold controlled visual density and was not inferential. The complete 101-combination table is supplied as Supplementary Data.

Five whole-corpus refits tested scope and coordinate assumptions. The human-only analysis removed three nonhuman studies. The known-space analysis removed contrasts without recognised MNI or Talairach labels. The coordinate-provenance analysis applied the retained coordinate-version replacements and removed invalid or seed-only coordinates. Two lexical analyses retained studies with word-bounded consciousness or awareness terms in title or abstract and in title alone. Each lexical sensitivity fitted and corrected one whole-corpus ALE map, so its coverage was treated as a scope sensitivity rather than a direct re-estimate of the seven-domain union.

#### Parcel spread and spatial controls

Parcel analyses used the grey-matter-filtered FreeSurfer Destrieux aparc.a2009s plus aseg atlas in MNI volume space [10]. Intersecting the atlas with the common mask yielded 167 parcels, comprising 148 cortical, 17 subcortical and 2 cerebellar-cortex parcels. The atlas labelled 95,846 of the 142,985 eligible mask voxels. This incomplete tiling did not restrict voxelwise ALE inference.

Corrected voxels were assigned directly on the atlas grid. Nearest-parcel assignment within 4 mm was used only for the coordinate-profile split-half and hypergraph analyses. Coverage was calculated for any corrected voxel, at least 20 corrected voxels, at least 1% of eligible parcel volume and at least 5%. Additional summaries included normalised Shannon entropy, Gini concentration, effective parcel count and the share of assigned voxels in the most occupied parcel. Main Figure 3b and Supplementary Note 5 show the coverage criteria.

The compact-territory null asked whether one contiguous region with the observed voxel mass could reproduce the observed parcel spread. Each of 5,000 draws grew one 26-connected component of exact mass within the analysis domain. The primary statistic was the number of parcels containing at least 20 voxels. Upper-tail probabilities used the plus-one rule [11]. The test was applied to the primary reported-experiment joint-FDR union and the paper-collapsed cluster-FWE union and is displayed in main Figure 3d.

A complementary randomised-coordinate analysis relocated all 17,700 foci independently with replacement according to whole-NeuroStore reporting density plus a small pseudocount. The construction preserved studies, experiments, experiment-level focus counts, semantic-domain assignments and ALE kernels. Each of 1,000 draws reran the seven domain-specific ALE models. The evidence field was the voxelwise maximum of the seven uncorrected z maps. Normalised parcel entropy was compared after exact matching of atlas-assigned voxel mass at $K = 16,675$ voxels. A lower-mass sensitivity selected the strongest $K = 14,000$ atlas voxels, all lying within each draw's joint-FDR union. Moran's $I$ assessed residual smoothness differences (Supplementary Note 8 and Supplementary Table 8).

The matched-literature control compared the paper-collapsed consciousness map with 1,000 samples from the wider NeuroSynth literature. Samples matched the number of papers and the complete distribution of foci per paper after excluding records that overlapped with the target or contained direct consciousness or awareness title terms. All target and comparator papers used a common $N = 20$, identical ALE and FDR procedures, the same atlas and matched spatial mass. Normalised parcel entropy was the primary statistic. Because only four primary controls contained the complete target mass within their FDR extent, a second comparison used $K = 16,000$ voxels, which lay below the FDR extent of every map. Focus-matched functional-topic literatures and a separately constructed language corpus were descriptive companions (Supplementary Note 9 and Supplementary Table 9).

#### Spatial decomposition of study maps

One modelled-activation map was created for each of the 576 studies with at least one in-mask focus. The retained foci were smoothed with 10-mm FWHM Gaussian kernels, summed within study and normalised to unit length so that studies reporting more foci did not dominate the matrix. The resulting study-by-voxel matrix was decomposed with truncated singular-value decomposition using up to 30 components. Component energy was the squared singular value divided by the matrix Frobenius norm.

Two null priors sampled locations from either the smoothed focus density of the consciousness corpus or whole-NeuroStore reporting density. Both added a small pseudocount, preserved each study's focus count, smoothing and row normalisation, and contributed 5,000 null matrices. The leading-component statistic used lower-tail inference because a dominant common source predicts greater first-component concentration. Cumulative five- and ten-component statistics used two-sided probabilities. Main Figure 4d displays the complete saved null draws at $k = 1$, 5 and 10; intermediate draws were not retained.

#### Reliability, domain influence and coordinate validation

The coordinate-profile split-half analysis randomly divided the 579 studies 1,000 times. Foci were assigned to atlas parcels directly or to the nearest parcel within 4 mm, and parcel-count profiles were correlated across halves. This measured the stability of the corpus-wide coordinate distribution rather than the reproducibility of fitted ALE maps (Supplementary Note 6).

A separate map-based analysis constructed 100 complementary partitions within each semantic domain and reran the complete seven-domain ALE procedure in both halves. All 49 cross-half domain correlations were retained for every partition. The confirmatory statistic was the mean correlation between maps bearing the same domain label, evaluated against all $7! = 5,040$ global relabellings of the second-half domains. Thresholded-map overlap, parcel-set agreement, peak displacement and domain identification were descriptive (Supplementary Figure 1, Supplementary Note 6 and Supplementary Table 6).

The leave-one-domain-out analysis asked whether one exceptional domain concealed a large six-way intersection. Each domain was removed in turn and the intersection of the remaining six maps was compared with position-matched exclusions across 1,000 study-label permutations. Each position was standardised by its own null mean and standard deviation before maximum-statistic family control. Remaining-six union extent and the contribution of the omitted map were descriptive (Supplementary Note 7 and Supplementary Table 7).

Coordinate checks documented space labels and analysis-specific representation exclusions. Contrasts that could not generate a valid modelled-activation map or parcel hyperedge were excluded only from analyses requiring that representation.

#### Functional-map reconstruction

The reconstruction used 28 source-study-excluded NeuroSynth maps divided equally among executive-control, mnemonic/conceptual-social, perceptual/sensorimotor and organismic-affective families. Twenty-one topics that did not describe a coherent psychological function and the direct consciousness-adjacent topic were excluded. Individual maps entered the regression, while family membership defined model subsets and contribution allocations. Supplementary Table 4 reports the inclusion and exclusion status of all 50 topics, and Supplementary Table 14d lists the 28 selected-map memberships.

Ridge regression was used because the functional maps were correlated and regularisation stabilised prediction from overlapping predictors [12]. The 2-mm grid was divided into contiguous $8 \times 8 \times 8$-voxel blocks, approximately 16-mm cubes, which were assigned to five outer folds. Four inner spatial folds selected the ridge penalty from 0.1, 1, 10, 100, 1,000 and 10,000. Reporting density, predictor centring and scaling were estimated from training voxels only and applied to held-out voxels [13].

The density-only model and all 16 subsets of the four families were fitted. Performance was reported as held-out $R^2$ for the full target and residual held-out $R^2$ after reporting-density adjustment. Exact Shapley values averaged each family's added predictive value across every subset and entry order [14]. Uncertainty used 5,000 spatial-block bootstrap samples. A residual increment of $\Delta R^2 = 0.020$ was defined as the smallest practically meaningful increment.

Dependence among the functional maps was characterised with correlations among family-average templates and by estimating how well each family's maps could be reconstructed from the remaining 21 predictors. Contrast-level scores were also averaged within independent papers and used to calculate Pearson, Spearman, partial Pearson and partial Spearman matrices with paper-level bootstrap intervals. These analyses contextualised the Shapley allocations rather than providing independent contribution estimates (Supplementary Note 15 and Supplementary Table 15).

Spatial-target specificity used 5,000 BrainSMASH surrogate targets [6]. The ALE target was separated into reporting-density and residual components, and BrainSMASH randomised the residual field using Euclidean distances among MNI voxel centres. The implementation used 2,500 nearest neighbours, 500 sampled locations, 25 lag bins, an exponential kernel and delta values 0.5, 0.7, 0.9 and 1.0. Each surrogate was residualised again, recentred, scaled to the observed residual variance and recombined with the fixed density component. Surrogates were evaluated with the same mask, folds, nested tuning and predictors as the observed target. Variogram diagnostics compared the observed residual with the postprocessed surrogates; the result is displayed in main Figure 5c.

#### Context-dependent functional profiles

The descriptive context heat map repeated the complete 16-subset reconstruction and Shapley procedure for each unthresholded semantic-domain ALE target. It used all 28 individual topic maps and supplied no cellwise inference. Formal inference used four separately constructed family-average templates. Each coordinate-bearing contrast received four spatial-similarity scores, which were averaged within paper or independent sample, yielding 577 paper-level profiles. The descriptive allocations are shown in main Figure 5d.

The four dimensions were standardised before a one-way Euclidean PERMANOVA [15] tested whether profiles differed across semantic domains. Domain labels were permuted 10,000 times while domain sizes were preserved. The upper-tail probability used the plus-one rule. A companion dispersion analysis [16] assessed whether the result could be explained by unequal within-domain spread. The complete analysis was repeated without standardisation.

#### Regional-locus and self-related comparisons

Seven binary regional predictors represented basal ganglia, brainstem, frontoparietal workspace, insula and operculum, prefrontal cortex, posterior hot zone and thalamus. They were deterministic unions of parcels from the fixed 167-label Destrieux and FreeSurfer atlas and entered jointly as one regional block. Regional-only, functional-map and combined models used the same target, nuisance adjustment, folds, tuning and 5,000 paired spatial-block bootstrap samples. Practical equivalence required the complete 95% interval for the regional increment beyond the functional maps to lie within $[-0.020, 0.020]$. Main Figure 5b,c and Supplementary Note 16 describe the regional analysis.

The self-related partition contained seven narrative maps related to speech and language, social cognition, memory, semantic representation and autobiographical scene construction, and seven embodied maps related to bodily need, motor and action processes, body and spatial representation, affect, pain and multisensory processing. All fourteen maps were removed from their parent families before either set was restored, preventing predictor duplication. Four models compared the remaining maps alone, with the narrative set, with the embodied set and with both sets. The same nested cross-validation, paired bootstrap and exact two-block Shapley decomposition were used. Exact topic indices and curated functional glosses are provided in Supplementary Table 14e; main Figure 5f displays the allocations.

#### Information sharing among functional families

The four fixed family-average functional templates and the whole-corpus ALE target were residualised against reporting density and standardised. Each field was binarised with a greater-than-or-equal quantile rule. The primary threshold defined 50% of voxels as high, with 40% and 60% high-state fractions retained as descriptive sensitivities. Total mutual information tested whether the joint family state carried more target information than matched reference landscapes (main Figure 5g and Supplementary Note 17).

The reference distribution contained 1,000 target maps. In each draw, all 17,700 foci were relocated independently with replacement according to reporting density plus a small pseudocount while preserving 579 studies, 1,252 experiments, experiment-level focus counts and common $N = 20$ kernels. The ALE and residualisation pipeline was rerun for every reference. Spatial-quality criteria based on six-neighbour agreement and high-state and low-state component counts were applied before inference.

We examined all 15 non-empty source-subset mutual informations, four Shapley allocations, four fully conditioned increments, three concentration summaries and a four-source minimum-mutual-information partial information decomposition [17–19]. Synergy denoted information available only from combined families. Mixed-route redundancy denoted information available through two or more alternative routes, with at least one route combining families. Individual-family redundancy denoted information available separately from two or more individual families. Two-sided plus-one probabilities were corrected with Benjamini-Hochberg FDR within each named family. The decomposition contained 166 atoms, of which 78 had non-zero reference variance and entered atom-level inference. Fully conditioned increments were interpreted as upper bounds on unique information because they also include synergy involving the focal family (Supplementary Table 16).

#### Meta-analytic coactivation networks

Corrected convergence identifies where foci recur, but not the broader systems in which those territories are reported. We therefore used meta-analytic coactivation modelling, or MACM, to estimate which regions were repeatedly co-reported with each seed across the literature [20, 21]. Each domain's corrected territory served as a seed. Every contrast in the consciousness corpus containing at least one focus in that seed was selected. Seeds selecting fewer than ten contrasts would not have been analysed. Whole-brain ALE was rerun on the selected contrasts, and the resulting map was controlled with FDR at $q < .05$ within the common mask. Voxel and parcel coverage summarised the seven corpus-internal coactivation maps. MACM describes literature-level co-reporting rather than functional connectivity within individual brains (main Figure 6a).

For main Figure 6a, the seven corrected coactivation maps were binarised and summed voxelwise. Values from one through seven record how many maps contained each voxel. Nearest-neighbour or modal resampling preserved integer membership during rendering.

#### Context-dependent higher-order coalitions

MACM shows which regions accompany a seed across experiments, but it does not preserve which regions occur together within an individual contrast. Each coordinate-bearing contrast was therefore represented as a hyperedge linking all atlas parcels containing at least one focus (main Figure 6b). Foci were assigned directly to the atlas or to the nearest valid parcel within 4 mm. The 4-mm rule assigned 99.7% of in-mask foci. Hyperedges containing fewer than two parcels were omitted, and exact duplicate parcel sets within a domain were represented once in the primary unweighted analysis. Their multiplicity was retained for weighted and paper-dependence sensitivities.

We counted the 26 connected motifs formed by triplets of hyperedges defined by Lee, Ko and Shin [22]. For each domain, 5,000 randomised hypergraphs were generated with bipartite double-edge swaps after a burn-in of 20 and thinning of five. The null preserved each parcel's hyperdegree and every hyperedge's size. Individual motif inference used the maximum absolute standardised statistic across motifs within domain.

Motifs were grouped before analysis into broader structural families including bridge chains, shared cores with satellites, nested expansions and replicated coalitions. Domain-by-family tests were corrected together with Benjamini-Hochberg FDR. Dependence on papers contributing several contrasts was assessed with paper-collapsed hypergraphs and 5,000 repeated one-contrast-per-paper samples. Cross-domain conservation was evaluated for individual motifs only and required a consistent direction, corrected within-domain evidence and stability under the paper-dependence controls. Main Figure 6c displays the three families with corrected effects; the replicated-coalition family had none.

To describe variation in the complete motif structure, each domain's 26-element characteristic profile was formed as $(\text{observed} - \text{null mean})/(\text{observed} + \text{null mean} + 1)$ for each motif and normalised across motifs. Pairwise cosine similarity was calculated for all 21 domain pairs and displayed descriptively in main Figure 6d; no pairwise probabilities or uncertainty intervals were calculated.

For the descriptive role maps, parcels were classified as core, intermediate/bridge and satellite according to membership across the three constituent hyperedges. Role counts were divided by all same-domain instances of the relevant motif family containing the parcel. Because these conditional fractions remained related to hyperdegree, each was regressed on natural-log hyperdegree and the residual was mapped. A minimum-support threshold was used only for display, and no parcel-level inferential probability was calculated. The resulting maps are shown in Supplementary Figure 2, panels a–g, and described in Supplementary Note 18.

#### Reproducibility and provenance

The analyses were orchestrated with Snakemake [23] through `workflow/Snakefile`, using settings in `config/publication.yaml`. The public repository at https://github.com/jskipper/distributed-consciousness records software versions, input and output hashes, random seeds and commands required to regenerate the reported outputs. Archived prompts, model outputs and consensus tables allow screening and labelling to be reproduced without live language-model calls.

### Supplementary Notes

##### Supplementary Note 1 | High-dimensional geometry of the semantic domains.

**Rationale and analysis.** The seven semantic domains were intended as an organising scaffold rather than sharply separated natural kinds. We therefore quantified their geometry in the original 1,024-dimensional embedding space rather than the two-dimensional UMAP display. For each domain we calculated mean cosine distance to its normalised centroid, mean pairwise cosine distance, nearest-centroid margin and cosine silhouette. Positive margins indicate that a study is closer to its assigned centroid than to every alternative centroid. We also retained all 21 pairwise centroid distances. Percentile intervals used 2,000 cluster-stratified bootstrap samples with domain assignments held fixed.
**Results.** Mean cosine silhouette was 0.0867 with a bootstrap interval of 0.0816–0.0923, and mean nearest-centroid margin was 0.0250 with an interval of 0.0238–0.0264. 7.08% of studies were no closer to their assigned centroid than to another domain centroid, and 11.05% had negative silhouettes. The closest centroids were conscious perception and attention and awareness in memory and learning, with distance 0.0267. The farthest were disorders and states of consciousness and affective awareness and fear, with distance 0.0744. Altered states was the most diffuse domain, whereas affective awareness and fear was the most compact.

Related patterns were visible in the map-based reliability analysis. Altered states had the lowest continuous-map and thresholded-map reliability, while the more compact affective-awareness domain was among the most stable. With only seven domains, these associations are descriptive rather than independent tests. Domain-level summaries are reported in Supplementary Table 2, with pairwise centroid distances, study-level geometry and bootstrap summaries supplied as Supplementary Data.
**Interpretation.** The domains show weak, graded organisation with permeable boundaries. They are useful for comparing related consciousness literatures, but they should not be interpreted as mutually exclusive study classes, latent continua or biological mechanisms.

##### Supplementary Note 2 | Descriptive anatomy of the whole-corpus landscape.

**Rationale and analysis.** The continuous whole-corpus ALE map was used as the descriptive target for functional decoding and reconstruction. To characterise its anatomy independently of the seven-domain inference, we examined corrected support at $z > 1.645$ with 26-neighbour connectivity. For each component we retained its extent, peak coordinate, corrected-z-weighted centre and atlas assignment. We also enumerated corrected support inside and outside the shared semantic-analysis mask and across every eligible atlas parcel.
**Results.** The corrected map contained 48,918 voxels in 16 components. The largest component contained 48,523 voxels and included the whole-map peak at MNI [-34, 20, -2], in left anterior-insular and opercular territory. The corrected-z-weighted whole-map centre was [-1.03, -14.72, 19.14]. Its approximately midline location reflects the balance of broad bilateral support rather than a focal centre of convergence. Corrected occupancy was enumerated across all eligible parcels, with atlas-unassigned mask voxels retained separately. Component and parcel summaries are reported in Supplementary Table 3.

The strongest positive reporting-density-adjusted topic associations involved anterior cingulate, pain, fear, response inhibition, and control or conflict. Language was the strongest negative association. These rankings describe the functional profile of the same broad landscape and are not independent confirmations of the later reconstruction analysis. Complete component, parcel and topic inventories are supplied as Supplementary Data.

##### Supplementary Note 3 | Spatial-null sensitivity of functional topic decoding.

**Rationale and analysis.** The reporting-density-adjusted topic analysis used a three-dimensional Fourier phase null as its primary spatial reference. To determine whether corrected survival depended

on that construction, we repeated the same 50 correlations with 10,000 volumetric BrainSMASH target surrogates. The observed target and each topic map were residualised separately against reporting density. Each BrainSMASH surrogate was residualised again, recentred and scaled to the observed residual target. The maximum absolute correlation across all 50 fixed topic maps supplied a two-sided family-wise reference. A separate set of 100 surrogates assessed variogram fidelity.
**Results.** Observed correlations were essentially unchanged between the Fourier and BrainSMASH implementations, indicating that differences in corrected survival primarily reflected the null distributions. Twenty-five of 50 topics passed their topic-specific Fourier spatial test, sixteen survived FDR across topics, ten survived the Fourier maximum-statistic correction and four survived the BrainSMASH maximum-statistic correction. The four BrainSMASH survivors were also Fourier survivors and involved anterior cingulate, fear and threat, response inhibition and control, and pain or somatosensory processing. Supplementary Table 4 lists all 50 topics and their status at each evidence tier.

The BrainSMASH surrogates were 17.13% smoother than the target at short lags and had a relative variogram RMSE of 0.3070. This mismatch broadens the chance-correlation distribution for fixed smooth topic maps and is conservative in the present direction. The four common survivors are therefore robust across the two tested null constructions. The six topics surviving only the Fourier family-wise test remain sensitive to null specification rather than being invalidated by the BrainSMASH result. The retained Fourier outputs do not contain surrogate fields, so the two nulls cannot be ranked directly by spatial fidelity. Complete 50-topic estimates and surrogate diagnostics are supplied as Supplementary Data.

#### Supplementary Note 4 | Fixed-kernel ALE sensitivity.

**Rationale and analysis.** Numeric source sample sizes were unavailable, so every contrast in the primary ALE analyses used the common value $N = 20$. We tested whether the principal maps depended materially on that localisation-uncertainty assumption by repeating the analyses with a common $N$ of 10, 20, 40 or 80 while holding coordinates, semantic assignments, evidence units, masks and correction procedures fixed. These values produced FWHM kernels of 10.0026, 9.2412, 8.8360 and 8.6263 mm. The analysis assesses sensitivity across a plausible range of common kernels rather than recovering heterogeneous source sample sizes.

For the FDR sensitivity, we refitted the whole-corpus map and the reported-experiment and paper-collapsed seven-map families at each value, applying the joint procedure to each seven-map family. Strict sensitivity used a same-kernel $N = 20$ Monte Carlo repeat with a different seed and complete paper-collapsed cluster-FWE families at $N = 10$ and $N = 80$.
**Results.** Across the joint-FDR seven-domain profiles, semantic-union Dice ranged 0.89744–0.94646, the largest absolute parcel-coverage change was 3.493 percentage points, and median domain Dice ranged from 0.86380 to 0.92511. Sensitivity broadly tracked the difference in kernel width from the $N = 20$ reference. The corpus-wide anatomical result therefore remained stable across the tested range.

The same-kernel $N = 20$ Monte Carlo repeat had union Dice 0.99953, median domain Dice 1.00000 and minimum domain Dice 0.97526. At $N = 80$, altered states was the only domain with Dice below 0.70, falling to 0.47896. Its strict map contained 259 voxels compared with 359 in the $N = 20$ reference. This change was far larger than same-kernel Monte Carlo variation. The median $N = 80$ domain Dice remained 0.89855, and the minimum domain Dice at $N = 10$ remained 0.88315, so the instability was local rather than a general deterioration of the seven-map architecture.

Supplementary Table 5 reports the global and strict results, with complete domain-level profiles supplied as Supplementary Data. The precise strict altered-states anatomy is provisional under this sensitivity, while the broad corpus-wide pattern remains supported.

#### Supplementary Note 5 | Parcel-size-aware coverage.

**Rationale and analysis.** Any-voxel parcel coverage can be inflated by isolated boundary voxels, while a single absolute voxel threshold has different implications for parcels of unequal size. We therefore recalculated coverage for each corrected seven-domain union using four criteria. These were any corrected voxel, at least 20 corrected voxels, at least 1% of eligible parcel volume and at least 5%. Denominators were each parcel's labelled voxel count after intersection with the common analysis mask.
**Results.** The primary reported-experiment joint-FDR union occupied 152/167 parcels by the any-voxel criterion, 119 at 20 voxels, 144 at 1% and 122 at 5%. Broad coverage persisted across the reported-experiment and paper-collapsed joint-FDR and cluster-FWE profiles under every criterion. These results are displayed in main Figure 3b and tabulated in main Table 1.

The absolute and proportional criteria are complementary rather than uniformly ordered because parcel volumes differ. Their agreement shows that broad coverage did not depend on isolated edge voxels or on one threshold that ignored parcel size. Complete parcel-level occupancies, eligible volumes, fractions and denominator tables are supplied as Supplementary Data.

##### Supplementary Note 6 | Split-half reliability of semantic-domain maps.

**Rationale and analysis.** We assessed whether the seven semantic-domain topographies retained their collective identities across non-overlapping subsets of contributing studies. For each of 100 complementary partitions, studies were shuffled separately within domain and assigned without replacement to two halves. Each half-corpus passed independently through the primary seven-domain ALE and joint-FDR procedure. For every partition, all 49 ordered half-A by half-B continuous-map correlations were retained. The confirmatory statistic averaged the seven matched correlations. Its exact reference enumerated all $7! = 5,040$ global relabellings of the second-half domains.
**Results.** The matched mean was $r = 0.400$, compared with a reference mean of 0.263 and central limits of 0.217–0.334. The identity arrangement was the unique maximum, giving exact $P = 0.000198$. Supplementary Figure 1 shows the mean cross-half matrix, and Supplementary Table 6 reports domain-level summaries.

For 6 of the seven domains, the diagonal entry was the maximum in both its row and column. Altered states was the exception. Its median matched continuous-map correlation, thresholded-mask Dice and bidirectional identity were all lower than those of the more reliable domains. Across the full map family, continuous topographies were more stable than thresholded territories, parcel sets and peaks. Agreement-count maps had only moderate stability, which cautions against treating the exact high-order overlap location as perfectly reproducible.

A complementary 1,000-draw coordinate-profile check correlated nearest-parcel counts of reported foci rather than refitted ALE maps. It showed high reproducibility of the coarse corpus-wide anatomical profile but does not replace the map-based refits.
**Interpretation.** The semantic maps retained their collective identities across study halves, while the precise altered-states territory and thresholded high-order overlap were less stable. This is internal reliability within the retained corpus rather than an external replication.

##### Supplementary Note 7 | Leave-one-domain-out influence.

**Rationale and analysis.** We tested whether the sparse seven-way intersection concealed a large six-domain territory suppressed by one exceptional domain. Each domain was excluded in turn and the intersection of the remaining six maps was compared with position-matched exclusions across 1,000 study-label permutations. Because domain sizes differed, each exclusion was standardised against its own null mean and standard deviation before applying a maximum-statistic family test. Remaining-six union extent, voxels unique to the omitted map and omitted-map mass were descriptive companions.
**Results.** No exclusion revealed an unusually large six-domain intersection. Every observed six-way extent was below its position-specific null mean. The maximum observed standardised excess was -0.159, and the omnibus upper-tail result was $P = 0.9301$. The largest raw six-way extent was 71 voxels

after excluding disorders and states of consciousness, but its standardised excess was negative and its adjusted probability did not support an exceptional effect.

The remaining-six unions retained 83.8%–94.7% of the original union after every exclusion. Broad anatomical spread therefore persisted without any single domain. Supplementary Table 7 reports the inferential and descriptive results for all seven exclusions.

**Interpretation.** The sparse seven-way result does not conceal a large common territory blocked by one exceptional subliterature, and no single domain explains the broad union. The analysis remains a within-corpus robustness test and does not show that the seven domains are interchangeable.

##### Supplementary Note 8 | Randomised-coordinate distributedness control.

**Rationale and analysis.** The compact-territory null tests one connected region but does not reproduce the multifocal smoothness produced by ALE. We therefore generated complete ALE fields after relocating the observed coordinate evidence according to whole-NeuroStore reporting density. Each draw preserved studies, reported experiments, experiment-level focus counts, semantic-domain assignments and ALE kernels. The seven domain-specific ALE maps were refitted, and the voxelwise maximum of their uncorrected z maps supplied a common evidence field.

The primary comparison selected exactly 16,675 atlas-assigned voxels from the observed field and every null field, matching the atlas support of the observed reported-experiment joint-FDR union. The primary statistic was normalised Shannon entropy across 167 parcels. A lower-mass sensitivity selected the strongest 14,000 atlas voxels, all lying within each draw's joint-FDR union. Four topology summaries, including parcel count, component count and largest-component share, were descriptive.

**Results.** Observed primary entropy was 0.88624, compared with a null median of 0.87105 and central limits of 0.86255–0.87887. The two-sided probability was $P = 0.002$. The FDR-supported sensitivity gave the same qualitative conclusion. The four descriptive topology outcomes remained within their central null intervals.

The null fields were modestly smoother than observed under six-neighbour Moran's $I$. This difference was in the anti-conservative direction for a positive distributedness result. A bounded first-order location-shift sensitivity reduced the entropy excess but left the observation above the shifted central reference interval. This calculation was a sensitivity bound rather than a recalibrated probability.

**Interpretation.** The observed map showed a modest excess of parcel-distributedness relative to reporting-density-weighted randomised-coordinate ALE fields after matching spatial extent. The result does not show that this property is unique to consciousness. The matched-literature comparison addresses that question. Supplementary Table 8 reports the compact numerical results.

##### Supplementary Note 9 | Matched-literature distributedness control.

**Rationale and analysis.** A heterogeneous cognitive-neuroimaging literature may also produce broad convergence. We therefore compared the paper-collapsed consciousness map with 1,000 NeuroSynth samples matched to the target on paper count and the complete distribution of foci per paper. Records overlapping with the target or containing direct consciousness or awareness title terms were excluded. Target and comparator papers used the same $N = 20$ kernel, FDR procedure, atlas and exact spatial mass.

The primary comparison selected the 21,083 strongest corrected-z atlas voxels from every map, matching the FDR-significant atlas extent of the target. Because only four primary controls contained that many FDR-significant voxels, a second comparison used 16,000 voxels, which lay below the FDR extent of every target and control map. Every selected voxel therefore survived FDR in the second analysis. Normalised parcel entropy was primary. Parcel occupancy, component count and largest-component share were secondary.

**Results.** At the primary mass, consciousness entropy was 0.8869 compared with a null median of 0.8856 and central limits of 0.8767–0.8945. At the lower mass supported by every map, consciousness entropy was 0.8657 compared with a null median of 0.8662 and limits of 0.8554–0.8764. Both results lay near the centre of the matched-literature distributions, and the secondary spatial summaries were likewise central. Supplementary Table 9 reports both comparisons.

Twenty-eight focus-matched functional-topic literatures were retained as descriptive comparisons. Consciousness was broader than most of these more coherent subliteratures, while topic coherence was negatively related to parcel entropy. A separately constructed language corpus was also broad and differentiated across its subliteratures.

**Interpretation.** The evidence supports distributed organisation within the consciousness corpus but provides no evidence that consciousness is unusually parcel-distributed relative to matched heterogeneous neuroimaging literatures. The topic and language comparisons illustrate how heterogeneity and subliterature composition influence apparent spread.

##### Supplementary Note 10 | Observed high-order overlap anatomy.

**Rationale and analysis.** The agreement-count image records how many corrected semantic-domain maps contain each voxel. We enumerated every exact non-empty domain combination and formed cumulative masks at thresholds of at least four, five, six and seven domains. Six-neighbour components and Destrieux or FreeSurfer assignments were then summarised without additional selection or inference.

**Results.** Across the 25,800-voxel corrected union, 101 of the 127 possible non-empty domain combinations occurred. Exact agreement counts for one through seven maps were 18,055, 5,281, 1,508, 558, 267, 112 and 19 voxels. Main Figure 4b shows this complete count profile, while Main Figure 4c displays exact four- through seven-domain combinations containing at least ten voxels. The complete combination table is supplied as Supplementary Data.

Cumulative overlap contracted from 956 voxels in at least four maps to 398 in at least five, 131 in at least six and 19 in all seven. The at-least-four set comprised 34 components and was dominated by medial superior-frontal and anterior-cingulate and bilateral anterior-insular or opercular territories. It also contained a 19-voxel left-thalamic component and smaller right-thalamic involvement. Thalamic overlap was absent from sets shared by five or more domains.

Overlap across at least five maps contained large left anterior-insular or opercular, medial superior-frontal or cingulate, and right anterior-insular components. Overlap across at least six maps retained a left short-insular or opercular component, a medial superior-frontal or cingulate component, smaller right-insular territories and two singletons. The seven-way anatomy is described in Supplementary Note 11.

**Interpretation.** High-order overlap contracted sharply and changed in domain composition as the overlap threshold increased. The remaining anatomy therefore describes recurrent convergence zones rather than one stable domain combination or an invariant consciousness core.

##### Supplementary Note 11 | Magnitude and study-label reference for seven-domain overlap.

**Magnitude reference.** The substantive question was whether the exact cross-domain intersection constituted a spatially meaningful shared core. We used 298 voxels as a transparent magnitude reference. This was one tenth of the smallest corrected domain map, rounded upward. The observed intersection contained 19 voxels, corresponding to 0.0133% of the analysis mask and 0.639% of the smallest domain map. It remained below the reference under joint-FDR $q < 0.05$, joint-FDR $q < 0.10$, uncorrected $p < 0.001$ and uncorrected $p < 0.01$ definitions.

The intersection comprised two nine-voxel components and one atlas-unassigned singleton. One component involved left short-insular and opercular cortex. The other involved medial superior-frontal and mid-anterior-cingulate cortex. No atlas parcel was fully covered.

**Study-label reference.** To determine whether the intersection was unusually small for a seven-way partition of this literature, 1,000 permutations reassigned all studies without replacement to pseudo-domains preserving the observed size vector. Studies, experiments, coordinates and within-study focus structure were otherwise unchanged. Every draw refitted all seven ALE maps and repeated the joint correction across the complete domain-by-voxel family. The endpoint was the number of analysis-mask voxels significant in every corrected map, evaluated with a lower-tail plus-one probability.

The null distribution had mean 61.448, median 60, central limits of 4–132 and range 0–185 voxels. The observed intersection was below the null centre but not unusually small, with lower-tail $P = 0.118881$. Lower-order cumulative extents changed gradually from above the null centre at low overlap counts to below it at high counts. Those comparisons were descriptive. Supplementary Table 10 reports the cumulative reference and threshold sensitivity.

**Interpretation.** The result does not support a spatially substantial invariant core. The permutation result also shows that a 19-voxel intersection was not exceptionally small relative to arbitrary same-sized partitions, so context sensitivity is supported more strongly by the collective pairwise differentiation than by the seven-way extent alone.

##### Supplementary Note 12 | Spatial recurrence of high-order overlap.

**Rationale and analysis.** The study-label permutations also allowed us to ask a location question that differs from the extent tests. For each draw and each cumulative threshold from four through seven pseudo-domains, overlap-set membership was recorded separately for every voxel. We then calculated peak recurrence, territories present in at least 50% and 90% of draws, component anatomy, alignment with the observed overlap and association with whole-NeuroStore reporting density. The 50% and 90% criteria are descriptive recurrence summaries rather than significance thresholds.

**Results.** Peak recurrence values for overlap in at least four, five, six and seven maps were 100.0%, 100.0%, 98.2% and 72.1%. Four- and five-domain recurrence involved bilateral anterior-insular or opercular and medial superior-frontal or anterior-cingulate territories. The six-domain 90% territory was confined to left short-insular cortex. The seven-domain 50% territory was a 24-voxel left short-insular component, and no seven-way voxel reached 90% recurrence. A small thalamic component appeared only in the four-domain 50% territory.

The observed seven-way set and the 50% permutation-recurrent set shared 3 voxels. Mean recurrence across the observed seven-way voxels was 27.5%, with range 7.7%–56.6%. Every non-empty 50% and 90% recurrence territory lay in the top decile of whole-NeuroStore reporting density. Supplementary Table 11 reports the compact recurrence and anatomical summaries, while voxelwise maps are supplied as Supplementary Data.

**Interpretation.** Arbitrary partitions returned preferentially to anterior-insular and medial-frontal territory, but the strictest overlap location was not invariant and all consensus territory followed generic reporting topography. The result identifies preferential convergence zones rather than a consciousness-specific core.

##### Supplementary Note 13 | Pairwise differentiation among semantic-domain maps.

**Rationale and analysis.** A tiny strict intersection does not by itself show that the seven maps are collectively different, because arbitrary partitions can also have small all-way overlap. We therefore used the same complete study-label permutations to calibrate pairwise similarity across the map family. Dice between joint-FDR masks was primary. The confirmatory statistic was mean Dice across all 21 unordered pairs. For pair-specific inference, each observed lower-tail deficit was standardised by its pair-position null mean and standard deviation, and each draw's largest standardised deficit supplied single-step family-wise control. Jaccard coefficients were companion effect sizes. Pearson correlations among continuous z maps formed a separately corrected secondary family.

**Results.** Mean Dice across the observed pairs was 0.152559, compared with a null mean of 0.205627 and central limits of 0.1789–0.2272. No permuted mean was as small as the observed value, giving lower-tail $P = 0.000999$. Three pairwise deficits survived single-step family-wise correction. These involved disorders or states with bodily self or agency, disorders or states with affective awareness or fear, and affective awareness or fear with bodily self or agency. The largest observed overlap was between conscious perception or attention and bodily self or agency, but it received no upper-tail test and does not show unusually strong convergence.

Mean continuous-map correlation was also lower than its permutation reference, and more pair-specific deficits survived in that family. Continuous maps were more stable than thresholded masks in the split-half analysis, so this difference is interpreted as sensitivity to representation rather than independent replication. Supplementary Table 12 reports the calibrated omnibus and pairwise results, with complete Dice, Jaccard, Pearson and permutation tables supplied as Supplementary Data.

**Interpretation.** The semantic-domain maps were collectively more differentiated than arbitrary same-sized partitions of the same consciousness corpus. This within-corpus result does not identify semantic content as the sole cause of the differences, because task, modality, contrast and reporting composition covary with the literature domains.

##### Supplementary Note 14 | Measurement-source coverage and coordinate reporting volume.

**Rationale and analysis.** Domain differences could be exaggerated if some literatures contributed systematically more coordinates or contrasts per publication. We therefore joined the coordinate dataset, semantic assignments, retained NeuroStore records and independent publication units. The resulting inventory contained 1,252 coordinate-bearing contrasts nested in 579 publication units. Coordinates were treated as reported stereotactic peaks rather than source-image voxels retaining beta weights, entropy, centrality, metabolic values or other measurements.

Three fully observed publication-level quantities were tested. These were unique foci retained after pooling and deduplication, source contrasts and duplicate foci removed. For each quantity, a tie-corrected seven-group Kruskal-Wallis statistic and effect size were calculated. Null distributions used 10,000 permutations of domain labels across publication units, with a maximum statistic controlling the family of three tests.

**Results.** No domain difference was detected for unique retained foci, source contrasts or duplicate foci removed, and the rank-based effects were very small. Supplementary Table 13 reports the domain summaries and corrected probabilities. Contrast names were available for most records, but complete structured classifications of population, modality, task, source statistic, source measurement type, measurement derivation route, sample size and smoothing were unavailable. Text hints were retained only as search aids and were not used as inferential classifications.

No numeric source sample sizes were recovered for any of the 1,252 contrasts, and all stored values equalled the configured default. The ALE pipeline therefore imposed a common localisation-uncertainty kernel across domains. It did not establish equal enrolment or permit a comparison of genuine sample sizes. Sensitivity to the common-kernel assumption is reported in Supplementary Note 4.

**Interpretation.** The analysis provides no evidence that domain differences arose from unequal coordinate-reporting volume per publication. It cannot exclude differences in population, modality, task, preprocessing, source measurement or smoothing, which were not available as complete structured fields.

##### Supplementary Note 15 | Dependence among functional maps and paper-level profiles.

**Rationale and analysis.** Shapley values allocate predictive performance among correlated functional maps, so their ordering must be interpreted in light of predictor dependence. We described dependence at three levels. First, we calculated Pearson correlations among the four family-average spatial templates. Second, each topic map within one family was regressed on the 21 maps in the other families and the resulting multiple $R^2$ values were averaged within family. Third, contrast-level similarity scores were averaged within 577 independent paper or sample units and used to calculate Pearson, Spearman, partial Pearson and partial Spearman matrices. Percentile intervals used 5,000 paper-level bootstrap samples.

**Results.** The family-average templates formed a correlated executive-control, mnemonic/conceptual-social and perceptual/sensorimotor group. Organismic-affective was nearly orthogonal to executive-control and mnemonic/conceptual-social maps and negatively related to the perceptual/sensorimotor template. Paper-level profiles showed a different pattern, with the strongest association between perceptual/sensorimotor and organismic-affective scores.

Mean multiple $R^2$ was 0.714 for executive-control, 0.677 for mnemonic/conceptual-social, 0.709 for perceptual/sensorimotor and 0.605 for organismic-affective. Organismic-affective was therefore the least overlapping family and received the largest Shapley allocation. Lower collinearity may contribute to that conditional allocation, although it did not determine the full family ordering. Executive-control and perceptual/sensorimotor maps had similar subspace overlap but markedly different allocations.

Supplementary Table 15 reports the template correlations, principal paper-score associations and block-level overlap. Complete matrices, intervals and topic-level values are supplied as Supplementary Data. These summaries describe spatial and literature-level dependence rather than within-brain coactivation.

#### Supplementary Note 16 | Construction of the regional-locus predictors.

**Rationale and construction.** The regional-locus analysis compared the distributed functional maps with seven coarse anatomical predictors representing basal ganglia, brainstem, frontoparietal workspace, insula and operculum, prefrontal cortex, posterior hot zone and thalamus. Each mask was a deterministic union of parcels from the fixed Destrieux and FreeSurfer atlas on the common 2-mm grid. The masks served as predictors and did not restrict the analysis domain.

The predictors ranged from 1,789 to 30,537 voxels and differed substantially in extent and overlap. The frontoparietal mask contained the complete prefrontal mask, and other overlaps followed directly from the atlas definitions. At most 3 regional masks contained the same voxel. Main Figure 5b shows six of these loci. All seven binary masks, their parcel identifiers, voxel counts and overlap field are supplied as Supplementary Data.

**Predictive result.** The regional block explained non-zero residual variance when fitted alone. Adding all seven predictors to the complete functional-map model changed held-out residual prediction by $\Delta R^2 = -0.0012$ with a 95% interval of -0.0077–0.0057. The interval lay within the defined equivalence bounds, indicating no practically meaningful increment beyond the functional maps.

**Interpretation.** The masks are coarse operationalisations of theory-associated territories rather than complete implementations of the theories. The block-level result concerns additional prediction after the functional maps are present and does not imply that the constituent structures are irrelevant.

#### Supplementary Note 17 | Target information and partial information decomposition.

**Rationale and analysis.** The reconstruction analysis showed how well the functional maps predicted the continuous landscape, but not whether the information carried by the four families was redundant, synergistic or both. The four family-average templates and the whole-corpus ALE target were residualised against reporting density and binarised using a greater-than-or-equal quantile rule. The primary threshold defined 50% of voxels as high, with 40% and 60% thresholds as descriptive sensitivities. Total mutual information tested whether the joint four-family state carried more target information than matched reference landscapes.

Each of 1,000 reference targets was generated by relocating all 17,700 foci according to reporting density while preserving 579 studies, 1,252 experiments, experiment-level focus counts and common $N = 20$ kernels. The ALE and residualisation pipeline was rerun for every reference. Spatial-quality criteria based on local neighbour agreement and high-state and low-state component counts were applied before inference.

**Omnibus and allocation results.** The four templates jointly carried 0.0720 bits about whether voxels lay above or below the median of the residual landscape. Matched references had mean 0.0570 bits and central limits of 0.0518–0.0627. The excess was 0.0150 bits, with upper-tail $P = 0.000999$. Results at 40% and 60% high-state fractions were similar in direction.

All 15 non-empty source subsets were examined, together with Shapley allocations, fully conditioned increments and concentration summaries. Mnemonic/conceptual-social and organismic-affective maps retained additional above-reference information after conditioning on the other families. Fully conditioned increments include both unique information and synergy involving the family, so they provide upper bounds on unique contributions rather than pure unique information.

**Redundancy and synergy.** The minimum-mutual-information decomposition showed excess aggregate synergy of 0.0047 bits and excess mixed-route redundancy of 0.0114 bits. Mixed-route redundancy captured alternative routes to the same target information, with at least one route combining families. Individual-family redundancy, in which every alternative route was a single family, did not exceed the reference. Positive excess synergy arose at third order, while pairs contributed chiefly through alternative routes to the same target information. The decomposition contained 166 atoms, of which 78 had non-zero reference variance. Several atom-level reference distributions contained substantial numerical-zero mass, and no named synergistic combination was required for the main conclusion.

Supplementary Table 16 summarises the omnibus, allocation and information-type results. Complete subset, concentration, order-resolved and atom-level outputs are supplied as Supplementary Data. These are static spatial relations among functional maps and a meta-analytic target rather than measurements of temporal information exchange within individual brains.

##### Supplementary Note 18 | Parcel roles in higher-order coalitions.

**Rationale and enumeration.** The motif analysis established non-random coalition structure but did not identify where parcels tended to occupy core, intermediate or satellite positions. Within retained nested-expansion and shared-core-with-satellites motifs, parcel membership in three, two or one constituent hyperedges defined core, intermediate/bridge and satellite positions. Exact duplicate parcel sets were represented once within domain, matching the primary unweighted motif census. Role counts were divided by all same-domain instances of the corresponding motif family that contained the parcel.

**Dependence on reporting frequency.** Hyperdegree measures how often a parcel appeared across contrast hyperedges. Raw role counts were strongly associated with hyperdegree, and conditional role fractions reduced but did not remove this dependence. We therefore fitted each conditional role fraction as a function of natural-log hyperdegree. Positive residuals indicate greater role occupancy than predicted by that descriptive relation, and negative residuals indicate less. A minimum of 100 motif-family participations controlled display only.

**Results and interpretation.** The residual maps varied across domains and roles, but no parcel-level probabilities were calculated and no multiplicity correction was applied. Much of the apparent anatomy followed reporting frequency, so these maps should be read as descriptive localisation rather than evidence that a particular parcel preferentially occupies a coalition role. This limitation does not affect the motif-count inference, whose randomised-hypergraph null preserved parcel hyperdegree and hyperedge size.

Supplementary Figure 2 shows the residual role maps after adjustment for hyperdegree and application of the minimum-support display threshold. Complete role counts, conditional fractions, fitted values, degree diagnostics and atlas-space maps are supplied as Supplementary Data.

#### Supplementary Figures

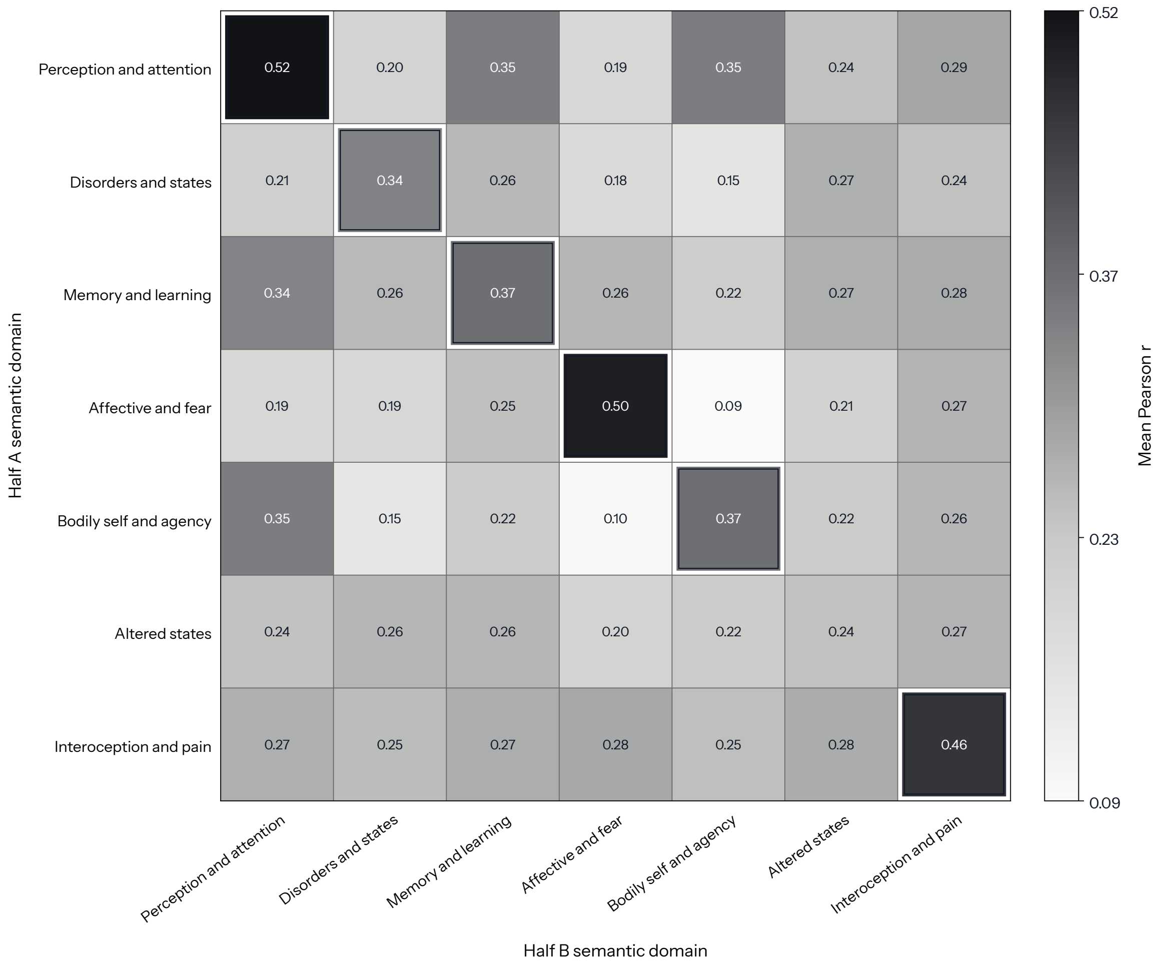


**Supplementary Figure 1 | Semantic-domain maps retain collective identity across study halves.**

Cells show mean Pearson correlations between continuous ALE maps from complementary study halves. For 6 of the seven domains, the outlined diagonal entry is the maximum in both its row and column, with altered states as the exception. The confirmatory statistic averaged diagonal values across domains and partitions and was evaluated against all $7! = 5,040$ global relabellings of the second-half domains. The identity arrangement was the unique maximum with exact $P = 0.000198$.

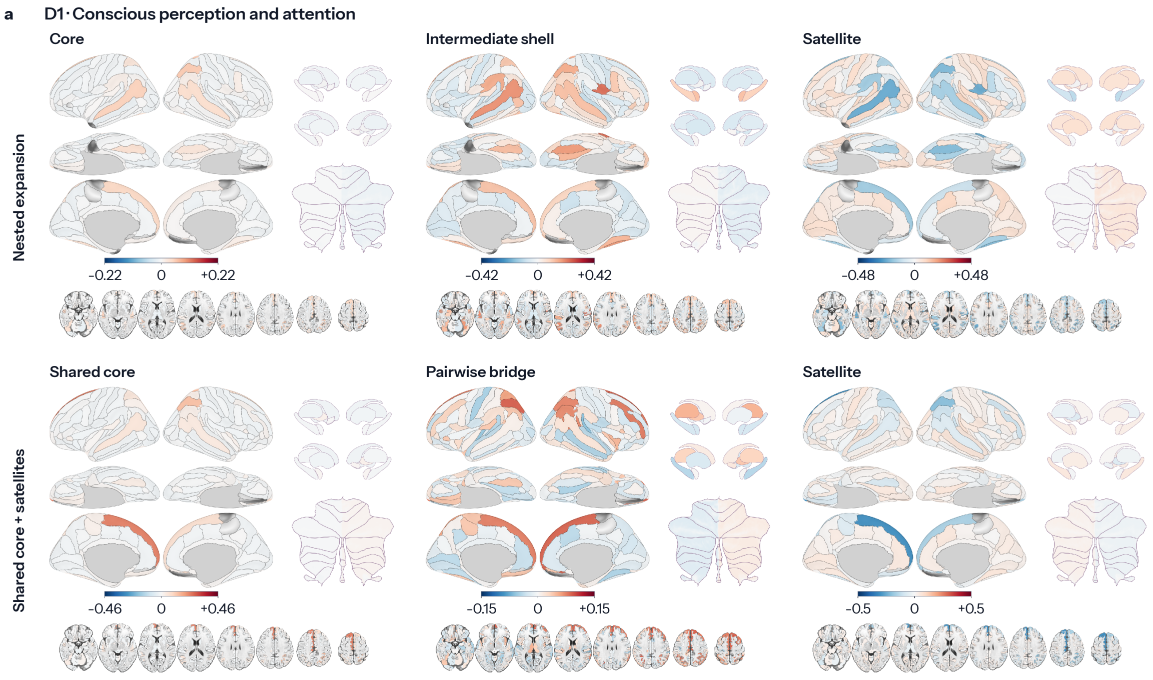
a
D1 · Conscious perception and attention
Core
Intermediate shell
Satellite
Nested expansion
−0.22
0
+0.22
−0.42
0
+0.42
−0.48
0
+0.48
Shared core
Pairwise bridge
Satellite
Shared core + satellites
−0.46
0
+0.46
−0.15
0
+0.15
−0.5
0
+0.5

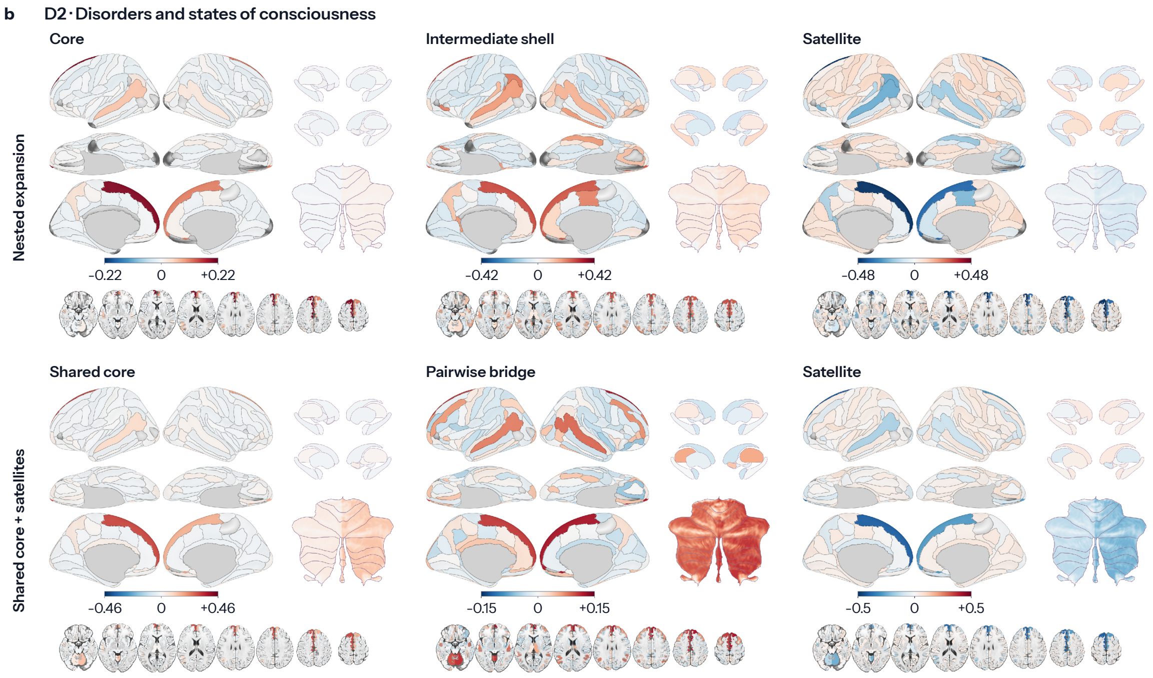
b
D2 · Disorders and states of consciousness
Core
Intermediate shell
Satellite
Nested expansion
−0.22
0
+0.22
−0.42
0
+0.42
−0.48
0
+0.48
Shared core
Pairwise bridge
Satellite
Shared core + satellites
−0.46
0
+0.46
−0.15
0
+0.15
−0.5
0
+0.5

c

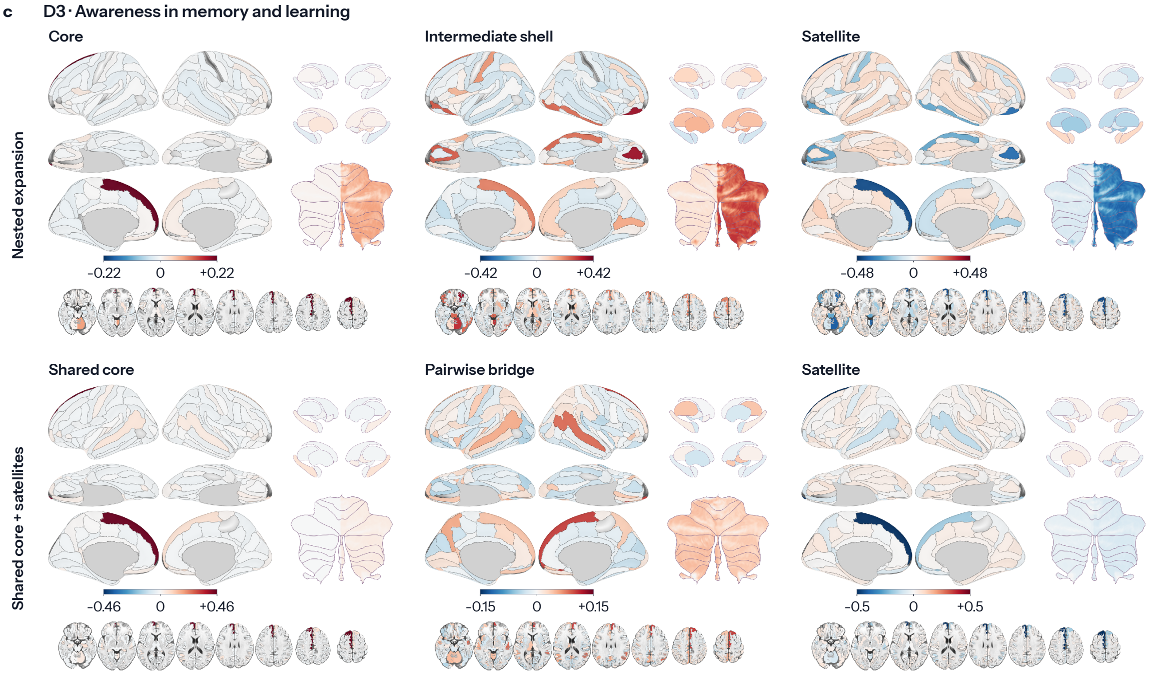
D3 · Awareness in memory and learning
Core
Intermediate shell
Satellite
Nested expansion
−0.22
0
+0.22
−0.42
0
+0.42
−0.48
0
+0.48
Shared core
Pairwise bridge
Satellite
Shared core + satellites
−0.46
0
+0.46
−0.15
0
+0.15
−0.5
0
+0.5

d

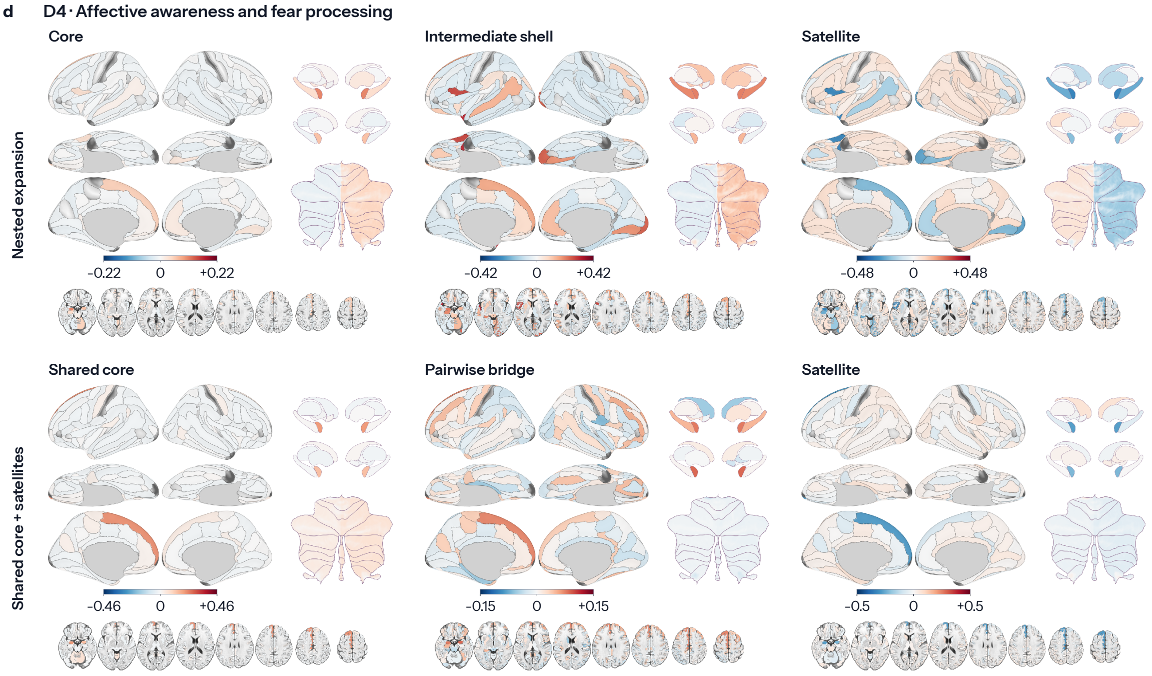
D4 · Affective awareness and fear processing
Core
Intermediate shell
Satellite
Nested expansion
−0.22
0
+0.22
−0.42
0
+0.42
−0.48
0
+0.48
Shared core
Pairwise bridge
Satellite
Shared core + satellites
−0.46
0
+0.46
−0.15
0
+0.15
−0.5
0
+0.5

e D5 · Bodily self-consciousness and agency

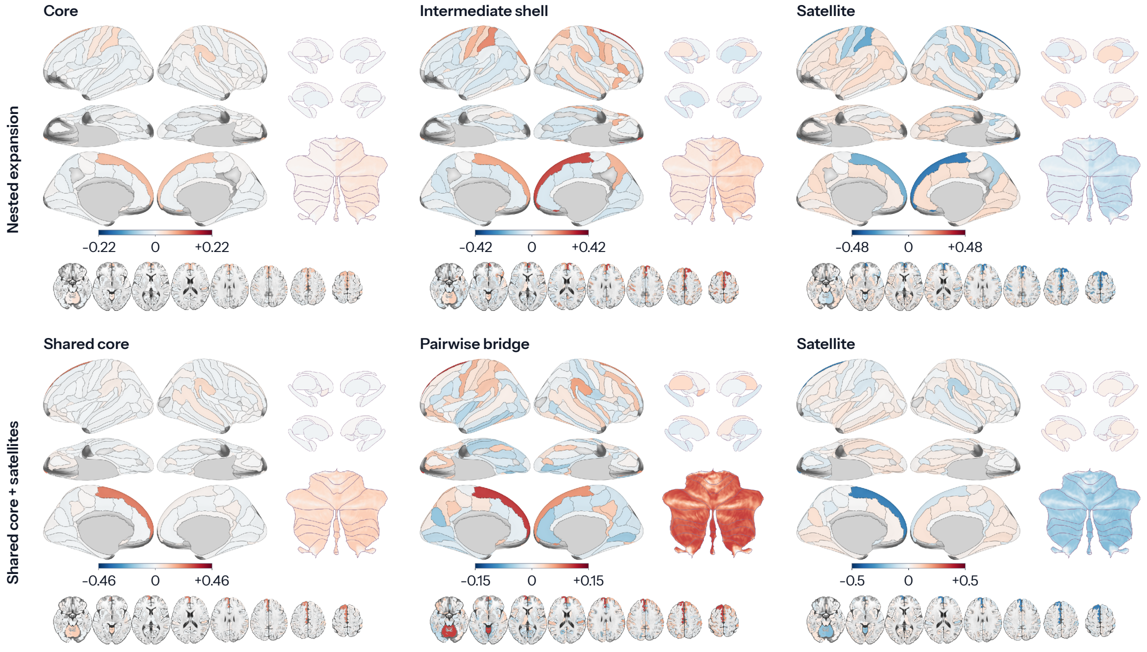


f D6 · Altered states of consciousness

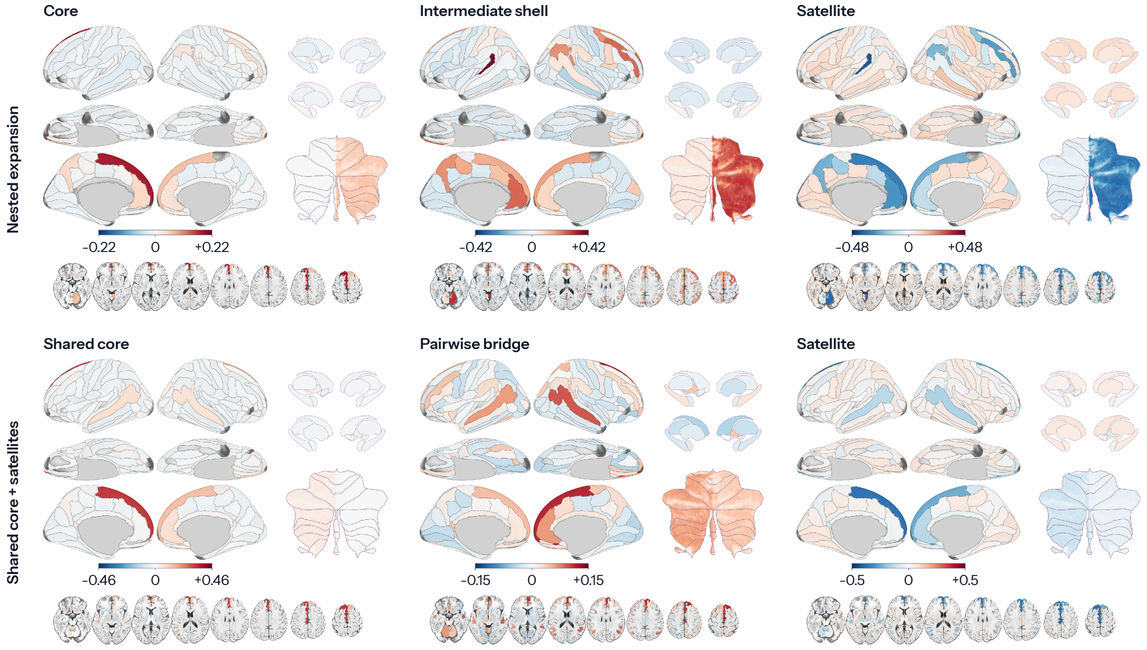

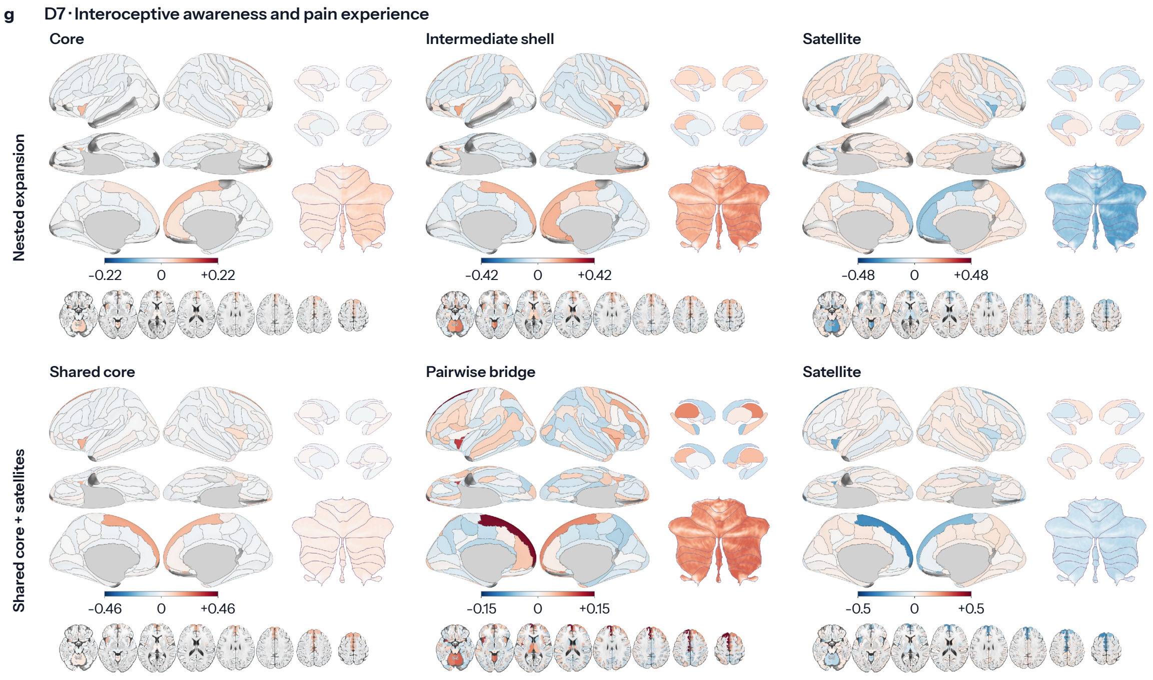


**Supplementary Figure 2 | Descriptive anatomical distribution of parcel roles in higher-order coalitions.**

Panels a–g follow D1–D7 in main Figure 1. Each domain plate shows nested-expansion core, intermediate-shell and satellite roles in the top row, and shared-core, pairwise-bridge and satellite roles for shared-core-with-satellites motifs in the bottom row. Maps are displayed on inflated cortical surfaces with subcortical, cerebellar and axial views. Colour shows the residual after regressing conditional role fraction on natural-log hyperdegree, using symmetric role-specific scales shared across domains; blue indicates negative and red positive residuals. Only parcels with at least 100 relevant motif-family participations are shown. The maps are descriptive; no parcel-level inference was performed and no multiplicity correction was applied.

### Supplementary Tables

**Supplementary Table 1 | Corpus construction, semantic stability and scope/coordinate sensitivities.**

| Section | Item | Value | Definition |
|---|---|---|---|
| Corpus flow | Base studies | 32,411 | NeuroStore/Neurosynth Compose local base |
| Corpus flow | Wide-net candidates | 3,388 | broad candidate screen |
| Corpus flow | Included studies | 579 | two-tier corpus |
| Corpus flow | Contrasts | 1,252 | coordinate-bearing NiMARE records |
| Corpus flow | Reported foci | 17,700 | reported coordinates |
| Study screen | Tier I unanimous inclusions | 695 | unanimous include among OpenAI, Claude and Gemini |
| Study screen | Tier II majority exclusions | 116 | removed when at least two models voted methods exclusion |
| Scope/coordinate sensitivity | Human-only scope | 576 studies, Dice 0.9936, $r = 0.9992$ | one-map whole-corpus FDR refit |
| Scope/coordinate sensitivity | Known coordinate space | 499 studies, Dice 0.9319, $r = 0.9803$ | one-map whole-corpus FDR refit |
| Scope/coordinate sensitivity | Coordinate provenance filter | 574 studies, Dice 0.9832, $r = 0.9969$ | one-map whole-corpus FDR refit |
| Semantic stability | Configured baseline | 8 | seed 42, 30 neighbours, resolution 1.0 |
| Semantic stability | Grid agreement with baseline | ARI 0.719, NMI 0.828 | 27 perturbation solutions |
| Semantic stability | Cluster-count modes | 7 and 11 | each occurred in 5/27 solutions, with the lower mode selected on tie |
| Semantic stability | Selected representative | 7 clusters, mean pairwise ARI 0.699 | seed 42, 50 neighbours, resolution 1.5 |
| Semantic stability | Direct embedding graph | ARI 0.655, NMI 0.686 | 7 clusters, with agreement measured against the configured baseline |
| Semantic domains | D1 Conscious perception and attention | 116 studies, 243 contrasts | 8,536 joint-FDR corrected voxels |
| Semantic domains | D2 Disorders and states of consciousness | 88 studies, 170 contrasts | 4,768 joint-FDR corrected voxels |
| Semantic domains | D3 Awareness in memory and learning | 87 studies, 191 contrasts | 5,165 joint-FDR corrected voxels |
| Semantic domains | D4 Affective awareness and fear processing | 82 studies, 161 contrasts | 5,224 joint-FDR corrected voxels |
| Semantic domains | D5 Bodily self-consciousness and agency | 76 studies, 168 contrasts | 4,713 joint-FDR corrected voxels |
| Semantic domains | D6 Altered states of consciousness | 69 studies, 173 contrasts | 2,973 joint-FDR corrected voxels |
| Semantic domains | D7 Interoceptive awareness and pain experience | 61 studies, 146 contrasts | 6,134 joint-FDR corrected voxels |

*Notes.* ARI is adjusted Rand index and NMI is normalised mutual information. Scope/coordinate-sensitivity Dice compares corrected masks and $r$ corrected z maps with the 579-study whole-corpus FDR map. Study inclusion used unanimous Tier I inclusion and no Tier II majority methods exclusion; the seven semantic domains are the scaffold selected by the stability analysis.

**Supplementary Table 2 | High-dimensional geometry of the semantic domains.**

| Semantic domain | Studies | Mean distance to assigned centroid | Mean within-domain pairwise distance | Mean nearest-centroid margin | Mean cosine silhouette | Non-positive nearest-centroid margin (%) | Negative cosine silhouette (%) |
|---|---|---|---|---|---|---|---|
| Conscious perception and attention | 116 | 0.1083 | 0.2066 | 0.0224 | 0.0748 | 7.8 | 12.9 |
| Disorders and states of consciousness | 88 | 0.1126 | 0.2150 | 0.0355 | 0.1138 | 2.3 | 3.4 |
| Awareness in memory and learning | 87 | 0.1072 | 0.2052 | 0.0197 | 0.0688 | 4.6 | 9.2 |
| Affective awareness and fear processing | 82 | 0.1002 | 0.1927 | 0.0278 | 0.1289 | 3.7 | 1.2 |
| Bodily self-consciousness and agency | 76 | 0.1054 | 0.2024 | 0.0259 | 0.0988 | 11.8 | 11.8 |
| Altered states of consciousness | 69 | 0.1141 | 0.2183 | 0.0190 | 0.0341 | 11.6 | 29.0 |
| Interoceptive awareness and pain experience | 61 | 0.1057 | 0.2036 | 0.0247 | 0.0838 | 9.8 | 13.1 |
| All domains | 579 | 0.1077 | 0.2062 | 0.0250 | 0.0867 | 7.1 | 11.1 |

*Notes.* All quantities use cosine geometry in the 1,024-dimensional BAAI/bge-large-en-v1.5 embedding space. A positive nearest-centroid margin means that a study is closer to its assigned centroid than to every alternative. Values describe weak, graded organisation rather than sharply separated classes; descriptive cluster-stratified bootstrap intervals are supplied as Supplementary Data.

**Supplementary Table 3 | Descriptive anatomy of the whole-corpus landscape.**

| Summary | Value |
|---|---|
| Corrected whole-corpus extent | 48,918 voxels (391,344 $mm^3$) |
| 26-neighbour components | 16 |
| Largest component | 48,523 voxels (99.19%) |
| Corrected whole-corpus peak | MNI [-34, 20, -2], corrected $z = 13.447$ |
| Corrected-z-weighted centre | MNI [-1.03, -14.72, 19.14] |
| Largest-component centre | MNI [-0.90, -14.92, 19.18] |
| Analysis-mask voxels assigned to atlas parcels | 95,846 voxels |
| Analysis-mask voxels outside the parcel atlas | 47,139 voxels |
| Eligible parcels without corrected convergence | 16 of 167 |

*Notes.* Weighted centres summarise the broad intensity distribution and are not anatomical convergence loci. Complete component and parcel inventories are supplied as Supplementary Data.

**Supplementary Table 4 | Functional topic decoding after reporting-density adjustment and spatial-null sensitivity.**

| Topic | Official NeuroSynth three-word label | Reconstruction family/status | Reporting-density-adjusted $r$ | Topic-specific spatial $P$ | FDR $q$ | Fourier max-$T$ $P$ | BrainSMASH max-$T$ $P$ | Strongest tier |
|---|---|---|---|---|---|---|---|---|
| 3 | cortex anterior cingulate | Not assigned | 0.286 | <0.001 | <0.001 | <0.001 | <0.001 | Both max-$T$ |
| 32 | pain somatosensory stimulation | Organismic-affective | 0.284 | <0.001 | <0.001 | <0.001 | <0.001 | Both max-$T$ |
| 13 | fear threat smokers | Organismic-affective | 0.221 | <0.001 | <0.001 | <0.001 | 0.014 | Both max-$T$ |
| 16 | response inhibition control | Executive-control | 0.218 | <0.001 | <0.001 | <0.001 | 0.017 | Both max-$T$ |
| 20 | control conflict task | Executive-control | 0.169 | <0.001 | <0.001 | <0.001 | 0.156 | Fourier max-$T$ |
| 37 | language reading word | Mnemonic/conceptual-social | -0.147 | <0.001 | <0.001 | 0.007 | 0.334 | Fourier max-$T$ |
| 36 | placebo treatment dopamine | Not assigned | 0.138 | <0.001 | <0.001 | 0.016 | 0.434 | Fourier max-$T$ |
| 27 | disorder adhd group | Not assigned | 0.132 | <0.001 | 0.002 | 0.027 | 0.502 | Fourier max-$T$ |
| 47 | attention attentional target | Executive-control | 0.131 | <0.001 | 0.002 | 0.030 | 0.515 | Fourier max-$T$ |
| 4 | stimulus time repetition | Consciousness-adjacent | 0.125 | <0.001 | <0.001 | 0.048 | 0.590 | Fourier max-$T$ |
| 12 | women men sex | Not assigned | 0.120 | <0.001 | 0.003 | 0.067 | 0.640 | FDR |
| 29 | stress ptsd trauma | Organismic-affective | 0.116 | 0.003 | 0.012 | 0.091 | 0.693 | FDR |
| 10 | food taste weight | Organismic-affective | 0.097 | 0.030 | 0.065 | 0.278 | 0.882 | Topic-specific |
| 38 | semantic category representations | Mnemonic/conceptual-social | -0.097 | 0.010 | 0.032 | 0.282 | 0.876 | FDR |
| 43 | magnetic mechanisms human | Not assigned | 0.097 | <0.001 | 0.003 | 0.287 | 0.894 | FDR |
| 17 | motor cortex hand | Perceptual/sensorimotor | 0.097 | 0.025 | 0.060 | 0.289 | 0.887 | Topic-specific |
| 30 | decision making risk | Organismic-affective | 0.095 | 0.006 | 0.022 | 0.313 | 0.901 | FDR |
| 15 | task performance cognitive | Executive-control | 0.094 | 0.008 | 0.028 | 0.330 | 0.911 | FDR |
| 8 | mpfc social medial | Mnemonic/conceptual-social | -0.094 | 0.020 | 0.056 | 0.333 | 0.905 | Topic-specific |
| 0 | network state resting | Not assigned | 0.092 | 0.019 | 0.055 | 0.365 | 0.922 | Topic-specific |
| 39 | stimulation tms bpd | Not assigned | 0.085 | 0.022 | 0.057 | 0.480 | 0.954 | Topic-specific |
| 1 | anxiety trait personality | Not assigned | 0.084 | 0.023 | 0.057 | 0.494 | 0.957 | Topic-specific |
| 2 | cerebellar cerebellum basal | Not assigned | 0.083 | 0.048 | 0.095 | 0.516 | 0.961 | Topic-specific |
| 26 | emotional amygdala negative | Organismic-affective | 0.073 | 0.076 | 0.141 | 0.708 | 0.988 | None |
| 14 | disease ad pd | Not assigned | 0.073 | 0.029 | 0.065 | 0.712 | 0.989 | Topic-specific |
| 6 | auditory speech temporal | Perceptual/sensorimotor | -0.072 | 0.128 | 0.216 | 0.734 | 0.989 | None |
| 34 | frequency hz ms | Not assigned | 0.070 | 0.047 | 0.095 | 0.758 | 0.992 | Topic-specific |
| 18 | number ips numerical | Mnemonic/conceptual-social | 0.069 | 0.087 | 0.156 | 0.775 | 0.993 | None |
| 41 | imagery mental events | Mnemonic/conceptual-social | -0.068 | 0.075 | 0.141 | 0.807 | 0.993 | None |
| 7 | reward feedback striatum | Organismic-affective | 0.055 | 0.166 | 0.245 | 0.950 | 0.999 | None |
| 28 | social empathy moral | Mnemonic/conceptual-social | 0.055 | 0.130 | 0.216 | 0.953 | 0.999 | None |
| 25 | spatial body human | Perceptual/sensorimotor | 0.053 | 0.192 | 0.274 | 0.963 | 0.999 | None |
| 44 | eye sleep gaze | Not assigned | 0.053 | 0.148 | 0.232 | 0.967 | 1.000 | None |
| 49 | depression mdd state | Not assigned | 0.053 | 0.155 | 0.235 | 0.967 | 1.000 | None |
| 35 | schizophrenia risk genetic | Not assigned | 0.049 | 0.146 | 0.232 | 0.984 | 1.000 | None |

| Topic | Official NeuroSynth three-word label | Reconstruction family/status | Reporting-density-adjusted $r$ | Topic-specific spatial $P$ | FDR $q$ | Fourier max-$T$ $P$ | BrainSMASH max-$T$ $P$ | Strongest tier |
|---|---|---|---|---|---|---|---|---|
| 33 | memory retrieval encoding | Mnemonic/conceptual-social | -0.046 | 0.229 | 0.317 | 0.992 | 1.000 | None |
| 9 | memory working wm | Executive-control | 0.041 | 0.297 | 0.401 | 0.998 | 1.000 | None |
| 40 | face faces facial | Perceptual/sensorimotor | -0.033 | 0.462 | 0.550 | 1.000 | 1.000 | None |
| 22 | method group approach | Not assigned | 0.031 | 0.345 | 0.453 | 1.000 | 1.000 | None |
| 48 | prefrontal cortex pfc | Executive-control | 0.029 | 0.434 | 0.530 | 1.000 | 1.000 | None |
| 46 | hemisphere language stroke | Not assigned | -0.027 | 0.418 | 0.523 | 1.000 | 1.000 | None |
| 23 | asd autism group | Not assigned | -0.025 | 0.384 | 0.492 | 1.000 | 1.000 | None |
| 5 | gyrus frontal inferior | Not assigned | 0.022 | 0.509 | 0.592 | 1.000 | 1.000 | None |
| 11 | learning training practice | Executive-control | 0.019 | 0.560 | 0.636 | 1.000 | 1.000 | None |
| 42 | visual cortex sensory | Perceptual/sensorimotor | -0.014 | 0.733 | 0.788 | 1.000 | 1.000 | None |
| 45 | motion perception visual | Perceptual/sensorimotor | 0.014 | 0.741 | 0.788 | 1.000 | 1.000 | None |
| 31 | model models prediction | Not assigned | 0.013 | 0.644 | 0.716 | 1.000 | 1.000 | None |
| 19 | action actions observation | Perceptual/sensorimotor | 0.011 | 0.798 | 0.832 | 1.000 | 1.000 | None |
| 21 | matter volume structural | Not assigned | 0.004 | 0.926 | 0.926 | 1.000 | 1.000 | None |
| 24 | age adults older | Not assigned | -0.003 | 0.916 | 0.926 | 1.000 | 1.000 | None |

*Notes.* All 50 topics are shown. The official three-word labels come from the NeuroSynth LDA vocabulary; the curated functional glosses in Supplementary Table 14d,e summarise these labels and the fuller topic-word profiles for the selected maps. E/M/S/O and consciousness-adjacent labels are analysis-specific reconstruction statuses; 'Not assigned' denotes exclusion from the prespecified EMSO grouping. Main Figure 2b displays the 13 E/M/S/O topics that passed the topic-specific spatial test; the consciousness-adjacent topic remains in this complete table. The strongest tier is both nulls, Fourier or BrainSMASH max-$T$, FDR, topic-specific, or none. Adjusted $r$, topic-specific spatial $P$, FDR $q$ and Fourier max-$T$ $P$ come from the primary Fourier analysis; the BrainSMASH column comes from the parallel null sensitivity. Totals were 25 topic-specific, 16 FDR, 10 Fourier max-$T$ and 4 under both nulls.

**Supplementary Table 5 | Fixed-kernel ALE sensitivity.**

**a, Joint FDR semantic profiles.**

| Profile | Assumed $N$ | FWHM (mm) | Union voxels | Union coverage change from $N = 20$ (pp) | Union Dice | Median domain Dice | Minimum domain Dice |
|---|---|---|---|---|---|---|---|
| Reported-experiment | 10 | 10.003 | 30,795 | +3.493 | 0.911 | 0.875 | 0.825 |
| Reported-experiment | 40 | 8.836 | 23,233 | -1.795 | 0.946 | 0.925 | 0.900 |
| Reported-experiment | 80 | 8.626 | 21,791 | -2.804 | 0.914 | 0.882 | 0.844 |
| Paper-collapsed | 10 | 10.003 | 23,021 | +2.967 | 0.897 | 0.864 | 0.813 |
| Paper-collapsed | 40 | 8.836 | 16,573 | -1.542 | 0.937 | 0.917 | 0.894 |
| Paper-collapsed | 80 | 8.626 | 15,383 | -2.374 | 0.900 | 0.870 | 0.835 |

**b, Strict cluster-FWE sensitivity.**

| Comparison | Assumed $N$ | FWHM (mm) | Union voxels | Union Dice | Median domain Dice | Minimum domain Dice |
|---|---|---|---|---|---|---|
| Same-kernel Monte Carlo repeat | 20 | 9.241 | 14,893 | 1.000 | 1.000 | 0.975 |
| Lower sample-size extreme | 10 | 10.003 | 17,358 | 0.922 | 0.899 | 0.883 |
| Higher sample-size extreme | 80 | 8.626 | 12,863 | 0.919 | 0.899 | 0.479 |

**c, Local altered-states sensitivity at the higher extreme.**

| Domain | Assumed $N$ | Voxels | Dice to $N = 20$ |
|---|---|---|---|
| Altered states of consciousness | 80 | 259 | 0.479 |

*Notes.* $N = 20$ defines the reference common localisation kernel; the other assumed sample sizes change kernel width rather than recover source enrolment. Joint FDR controls the complete domain-by-voxel family, whereas cluster-FWE is applied within domain with Bonferroni correction across seven domains. Panel c shows the only domain below the 0.70 descriptive stability threshold under strict inference.

**Supplementary Table 6 | Split-half reliability of semantic-domain maps.**

| Semantic domain | Studies in smaller half | Median continuous-map $r$ [2.5th–97.5th] | Median corrected-mask Dice | Median parcel Jaccard | Median peak displacement (mm) | Bidirectional identity |
|---|---|---|---|---|---|---|
| Conscious perception and attention | 58 | 0.519 [0.452–0.561] | 0.253 | 0.516 | 94.0 | 99.5% |
| Disorders and states of consciousness | 44 | 0.345 [0.283–0.384] | 0.082 | 0.353 | 52.2 | 91.0% |
| Awareness in memory and learning | 43 | 0.372 [0.302–0.417] | 0.169 | 0.396 | 58.7 | 66.0% |
| Affective awareness and fear processing | 41 | 0.496 [0.449–0.532] | 0.496 | 0.440 | 50.1 | 100.0% |
| Bodily self-consciousness and agency | 38 | 0.374 [0.320–0.415] | 0.077 | 0.397 | 64.9 | 76.5% |
| Altered states of consciousness | 34 | 0.244 [0.184–0.296] | 0.022 | 0.239 | 72.9 | 8.5% |
| Interoceptive awareness and pain experience | 30 | 0.460 [0.416–0.507] | 0.249 | 0.415 | 37.6 | 100.0% |

*Notes.* Values summarise 100 complementary within-domain partitions of the same corpus. Intervals are 2.5th–97.5th percentiles across dependent partitions, not confidence intervals from independent datasets. Bidirectional identity is the percentage of 200 half-direction comparisons in which the matched domain had the highest continuous-map correlation. Domain rows are descriptive; only the global exact relabelling test was confirmatory.

**Supplementary Table 7 | Leave-one-domain-out robustness of the six-way intersection and remaining-six union.**

**a, Position-matched six-way intersection.**

| Omitted semantic domain | Six-way intersection | Reference mean | Reference s.d. | Reference 95% limits | Standardised excess $z$ | Upper-tail $P$ | Family-wise $P$ |
|---|---|---|---|---|---|---|---|
| Conscious perception and attention | 19 | 68.2 | 39.7 | 4.0–155.0 | -1.241 | 0.906 | 0.998 |
| Disorders and states of consciousness | 71 | 78.0 | 43.9 | 5.0–169.0 | -0.159 | 0.532 | 0.930 |
| Awareness in memory and learning | 19 | 78.2 | 43.6 | 6.0–172.0 | -1.359 | 0.926 | 1.000 |
| Affective awareness and fear processing | 29 | 81.3 | 44.1 | 8.0–170.0 | -1.187 | 0.874 | 0.997 |
| Bodily self-consciousness and agency | 35 | 86.4 | 46.5 | 7.0–185.0 | -1.106 | 0.867 | 0.996 |
| Altered states of consciousness | 48 | 92.1 | 46.3 | 14.0–186.0 | -0.952 | 0.822 | 0.992 |
| Interoceptive awareness and pain experience | 24 | 104.4 | 47.9 | 19.0–200.0 | -1.677 | 0.961 | 1.000 |

**b, Remaining-six union and omitted-map summaries.**

| Omitted semantic domain | Remaining-six union | Retained (%) | Reference 95% limits | Unique to omitted map | Reference 95% limits | Omitted-map extent | Reference 95% limits |
|---|---|---|---|---|---|---|---|
| Conscious perception and attention | 21,625 | 83.8 | 11460.9–14561.1 | 4,175 | 1385.0–3258.3 | 8,536 | 3454.7–6628.8 |
| Disorders and states of consciousness | 23,219 | 90.0 | 12235.8–15117.2 | 2,581 | 884.9–2451.2 | 4,768 | 2287.9–5355.0 |
| Awareness in memory and learning | 23,989 | 93.0 | 12125.9–15138.3 | 1,811 | 846.8–2430.1 | 5,165 | 2217.7–5381.4 |
| Affective awareness and fear processing | 22,801 | 88.4 | 12347.6–15178.9 | 2,999 | 796.9–2316.2 | 5,224 | 1982.8–4973.6 |
| Bodily self-consciousness and agency | 23,550 | 91.3 | 12480.5–15306.4 | 2,250 | 743.9–2154.2 | 4,713 | 1871.0–4745.4 |
| Altered states of consciousness | 24,441 | 94.7 | 12600.0–15374.2 | 1,359 | 648.0–1996.2 | 2,973 | 1618.0–4444.2 |
| Interoceptive awareness and pain experience | 22,920 | 88.8 | 12764.9–15476.2 | 2,880 | 531.9–1836.2 | 6,134 | 1339.8–3989.2 |

*Notes.* The six-way intersection is calculated after omitting the named domain. References use 1,000 same-sized study-label partitions and are position-specific because observed domain sizes differ; family-wise probabilities use a single-step maximum statistic across the seven omissions. Panel b is descriptive, and retained percentages use the 25,800-voxel seven-domain union as denominator.

**Supplementary Table 8 | Randomised-coordinate distributedness control.**

| Measure | Analysis | Observed | Reference median | Reference central 95% | $P$ or status |
|---|---|---|---|---|---|
| Normalised parcel entropy | Primary, $K = 16{,}675$ | 0.88624 | 0.87105 | 0.86255–0.87887 | $P = 0.002$; +1.74% |
| Normalised parcel entropy | FDR-supported sensitivity, $K = 14{,}000$ | 0.87399 | 0.86126 | 0.85184–0.86993 | $P = 0.004$; +1.48% |
| Parcels with $\geq 20$ voxels | Full-union matched mass, $K = 25{,}800$ | 119 | 114 | 109.0–119.0 | Descriptive; no $P$ |
| Parcels per 1,000 voxels | Full-union matched mass, $K = 25{,}800$ | 4.612 | 4.419 | 4.225–4.613 | Descriptive; no $P$ |
| 26-neighbour components | Full-union matched mass, $K = 25{,}800$ | 88 | 76 | 61.0–92.0 | Descriptive; no $P$ |
| Largest-component share | Full-union matched mass, $K = 25{,}800$ | 0.375 | 0.511 | 0.262–0.728 | Descriptive; no $P$ |

*Notes.* Each of 1,000 draws relocated 17,700 foci according to reporting density, preserved study and reported-experiment focus counts and reran the seven domain-specific ALE models. Entropy used matched atlas-assigned mass at $K = 16{,}675$ and an FDR-supported $K = 14{,}000$ sensitivity; probabilities are two-sided plus-one and topology rows compare the observed full union with the strongest 25,800 voxels in each draw. Null fields were modestly smoother than observed. Separately, this control does not test whether distributedness is unique to consciousness.

**Supplementary Table 9 | Matched-literature distributedness control.**

| Metric | Analysis | Consciousness | Matched reference | | | Two-sided $P$ |
|---|---|---|---|---|---|---|
| | | | 2.5% | Median | 97.5% | |
| Normalised parcel entropy | Matched extent | 0.88688 | 0.87665 | 0.88559 | 0.89447 | 0.789 |
| Parcels occupied ≥1% | Matched extent | 148 | 140 | 145 | 150 | 0.370 |
| 26-neighbour components | Matched extent | 71 | 55 | 70 | 86 | 0.973 |
| Largest-component share | Matched extent | 0.290 | 0.182 | 0.359 | 0.669 | 0.611 |
| Normalised parcel entropy | FDR-supported extent | 0.86569 | 0.85541 | 0.86619 | 0.87636 | 0.933 |
| Parcels occupied ≥1% | FDR-supported extent | 133 | 127 | 134 | 141 | 0.789 |
| 26-neighbour components | FDR-supported extent | 70 | 63 | 77 | 92 | 0.452 |
| Largest-component share | FDR-supported extent | 0.157 | 0.139 | 0.197 | 0.403 | 0.228 |

*Notes.* Matched references are 1,000 NeuroSynth samples with the same paper count and complete foci-per-paper distribution, excluding overlapping records and records with direct consciousness or awareness title terms. The primary comparison used $K = 21{,}083$ atlas voxels; the $K = 16{,}000$ sensitivity lay below every map's FDR-supported extent. Entropy was primary, occupancy and topology were secondary, and no metric indicated that consciousness was unusually distributed relative to the matched heterogeneous literatures.

**Supplementary Table 10 | Magnitude and threshold sensitivity of seven-domain overlap.**

**a, Cumulative cross-domain overlap.**

| Domains containing a voxel | Observed voxels | Reference median | Reference 2.5% | Reference 97.5% |
|---|---|---|---|---|
| At least one | 25,800 | 15,193.0 | 13,869.6 | 16,515.3 |
| At least two | 7,745 | 4,986.5 | 4,446.0 | 5,523.0 |
| At least three | 2,464 | 2,027.0 | 1,786.9 | 2,293.0 |
| At least four | 956 | 938.5 | 793.0 | 1,065.1 |
| At least five | 398 | 483.5 | 340.9 | 578.0 |
| At least six | 131 | 223.0 | 116.0 | 316.0 |
| Seven-way intersection | 19 | 60.0 | 4.0 | 132.0 |

**b, Seven-way threshold sensitivity.**

| Threshold | Voxels | Components | Largest component |
|---|---|---|---|
| Joint FDR $q < 0.05$ | 19 | 3 | 9 |
| Joint FDR $q < 0.1$ | 66 | 5 | 36 |
| Uncorrected $p < 0.001$ | 14 | 3 | 9 |
| Uncorrected $p < 0.01$ | 139 | 4 | 78 |

*Notes.* Reference summaries come from 1,000 same-sized study-label partitions. The primary lower-tail test concerns the 19-voxel seven-way intersection; the 298-voxel meaningful-core reference is a prespecified descriptive magnitude bound. Other cumulative extents and threshold profiles are descriptive.

**Supplementary Table 11 | Spatial recurrence of high-order overlap.**

| Cumulative overlap | Peak recurrence (%) | Voxels at 50% recurrence | Voxels at 90% recurrence | Principal recurrent anatomy |
|---|---|---|---|---|
| At least four | 100.0% | 807 | 440 | Bilateral anterior insula or operculum, medial superior frontal or anterior cingulate, and small thalamic territory |
| At least five | 100.0% | 455 | 180 | Bilateral anterior insula or operculum and medial superior frontal or anterior cingulate |
| At least six | 98.2% | 192 | 33 | Left short-insular cortex at the 90% recurrence criterion |
| All seven | 72.1% | 24 | 0 | Left short-insular component at the 50% recurrence criterion |

*Notes.* Recurrence is the proportion of 1,000 same-sized study-label partitions in which a voxel entered each cumulative overlap. The 50% and 90% extents are descriptive recurrence criteria, not significance thresholds. Every non-empty recurrent territory lay in the top decile of whole-NeuroStore reporting density.

**Supplementary Table 12 | Pairwise differentiation among semantic-domain maps.**

| Comparison | Observed | Reference mean | Reference 95% limits | *P* |
|---|---|---|---|---|
| Mean pairwise Dice omnibus | 0.153 | 0.206 | 0.179–0.227 | <0.001 |
| Disorders and states of consciousness vs Affective awareness and fear processing | 0.076 | 0.219 | 0.108–0.318 | 0.038 |
| Disorders and states of consciousness vs Bodily self-consciousness and agency | 0.059 | 0.213 | 0.115–0.320 | 0.015 |
| Affective awareness and fear processing vs Bodily self-consciousness and agency | 0.060 | 0.204 | 0.097–0.305 | 0.041 |
| Mean pairwise continuous-map Pearson $r$ | 0.329 | 0.452 | 0.436–0.467 | <0.001 |

*Notes.* Observed values are compared with 1,000 same-sized study-label partitions. Omnibus values use lower-tail plus-one probabilities; pair-specific Dice values use single-step family-wise correction across all 21 pairs, and Pearson correlations form a separate corrected family. Complete matrices are supplied as Supplementary Data.

**Supplementary Table 13 | Coordinate-reporting volume and source-field availability.**

**a, Coordinate-reporting volume by semantic domain.**

| Semantic domain | Publication units | Unique retained foci | Source contrasts | Duplicate foci removed |
|---|---|---|---|---|
| Conscious perception and attention | 116 | 16.5 [9.0–35.0] | 1.0 [1.0–2.2] | 0.0 [0.0–0.0] |
| Disorders and states of consciousness | 88 | 18.5 [10.8–30.2] | 1.0 [1.0–2.0] | 0.0 [0.0–0.0] |
| Awareness in memory and learning | 87 | 21.0 [12.5–36.0] | 2.0 [1.0–3.0] | 0.0 [0.0–0.0] |
| Affective awareness and fear processing | 82 | 21.0 [12.0–34.0] | 2.0 [1.0–2.0] | 0.0 [0.0–0.0] |
| Bodily self-consciousness and agency | 76 | 20.0 [10.0–32.8] | 1.5 [1.0–2.2] | 0.0 [0.0–0.0] |
| Altered states of consciousness | 69 | 26.0 [11.0–41.0] | 2.0 [1.0–3.0] | 0.0 [0.0–0.0] |
| Interoceptive awareness and pain experience | 61 | 21.0 [13.0–50.0] | 1.0 [1.0–3.0] | 0.0 [0.0–1.0] |

**b, Key source-field availability.**

| Field | Available | Missing | Available (%) |
|---|---|---|---|
| Contrast name | 1,249 | 3 | 99.8 |
| Contrast description | 372 | 880 | 29.7 |
| Modality text hint | 1,077 | 175 | 86.0 |
| Measurement text hint | 1,047 | 205 | 83.6 |
| Numeric source sample size | 0 | 1,252 | 0.0 |
| Population, modality, task/paradigm, source statistic, source measurement type and measurement derivation route classifications | 0 | 1,252 | 0.0 |
| Smoothing FWHM | 0 | 1,252 | 0.0 |

*Notes.* Panel a gives median [interquartile range] per independent publication unit; no reporting-volume outcome differed after maximum-statistic family control (all adjusted $P \geq 0.342$). Panel b reports source-field availability across 1,252 contrasts; text hints were search aids, not classifications. Numeric source sample sizes were unavailable, so ALE used a common configured localisation-uncertainty kernel rather than actual source enrolment.

**Supplementary Table 14 | Functional-map reconstruction, regional comparison and self-related proxies.**

**a, Functional reconstruction and regional comparison.**

| Model or allocation | Estimate | Interval or companion |
|---|---|---|
| Reporting density only | Total held-out $R^2 = 0.7320$ | Nuisance baseline |
| Reporting density plus all 28 EMSO topic maps | Total held-out $R^2 = 0.7830$ | Absolute increment $\Delta R^2 = 0.0510$ |
| All 28 EMSO topic maps after foldwise reporting-density adjustment | Residual held-out $R^2 = 0.1902$ | 0.1400–0.2373 |
| Executive-control Shapley allocation | $\Delta R^2 = 0.0662$ | 0.0415–0.0900 |
| Mnemonic/conceptual-social Shapley allocation | $\Delta R^2 = 0.0297$ | 0.0151–0.0441 |
| Perceptual/sensorimotor Shapley allocation | $\Delta R^2 = 0.0035$ | -0.0078–0.0137 |
| Organismic-affective Shapley allocation | $\Delta R^2 = 0.0908$ | 0.0593–0.1217 |
| Theory-associated regional loci alone | Residual held-out $R^2 = 0.0296$ | Seven predictors |
| Increment of theory-associated regional loci beyond functional maps | $\Delta R^2 = -0.0012$ | -0.0077–0.0057 |

**b, Context-dependent functional profiles.**

| Test | Statistic | Multivariate $R^2$ | $P$ |
|---|---|---|---|
| Between-domain profile difference | pseudo-$F = 17.10$ | 0.153 | <0.0001 |
| Dispersion diagnostic | $F = 1.82$ | not applicable | 0.108 |

**c, Self-related proxy allocations.**

| Proxy | Shapley $\Delta R^2$ | 95% bootstrap interval |
|---|---|---|
| Embodied self-related proxy | 0.0185 | 0.0018–0.0340 |
| Narrative self-related proxy | 0.0074 | -0.0045–0.0189 |

**d, Functional-family topic membership.**

| Family | NeuroSynth topic index and curated functional gloss |
|---|---|
| Executive-control (E) | 9 working memory; 11 learning/practice; 15 task performance; 16 response inhibition; 20 control/conflict; 47 attention; 48 prefrontal/executive control |
| Mnemonic/conceptual-social (M) | 8 social cognition/mentalising; 18 numerical cognition; 28 social cognition/empathy; 33 memory retrieval; 37 language/reading; 38 semantic representation; 41 autobiographical scene construction |
| Perceptual/sensorimotor (S) | 6 auditory/speech perception; 17 motor processes; 19 action observation; 25 body/spatial representation; 40 face perception; 42 visual/multisensory processing; 45 visual motion perception |
| Organismic-affective (O) | 7 reward/feedback; 10 bodily need/food/taste; 13 fear/threat; 26 affect/emotion; 29 stress/trauma; 30 decision/risk/valuation; 32 pain/somatosensation |

**e, Self-related proxy topic membership.**

| Proxy | NeuroSynth topic index and curated functional gloss |
|---|---|
| Embodied self-related proxy | 10 bodily need/food/taste; 17 motor processes; 19 action observation; 25 body/spatial representation; 26 affect/emotion; 32 pain/somatosensation; 42 visual/multisensory processing |
| Narrative self-related proxy | 6 auditory/speech perception; 8 social cognition/mentalising; 28 social cognition/empathy; 33 memory retrieval; 37 language/reading; 38 semantic representation; 41 autobiographical scene construction |

*Notes.* Predictive estimates use spatially blocked cross-validation. Shapley values are conditional allocations among correlated families, not independent effects; the heat map is descriptive and the profile test is omnibus. Regional equivalence used ±0.020; its seven predictors were basal ganglia, brainstem, frontoparietal workspace, insula and operculum, prefrontal cortex, posterior hot zone and thalamus. Memberships in panels d and e come from canonical ledgers; curated glosses summarise the official three-word NeuroSynth labels in Supplementary Table 4 and fuller topic-word profiles. Embodied and narrative sets are proxies, not direct measures of selfhood.

**Supplementary Table 15 | Dependence among functional maps and paper-level profiles.**

**a, Family-average spatial-template correlations.**

| Pair | E–M | E–S | M–S | E–O | M–O | S–O |
|---|---|---|---|---|---|---|
| Pearson $r$ | 0.494 | 0.421 | 0.256 | 0.034 | -0.012 | -0.223 |

**b, Strongest paper-level pair across four dependence summaries.**

| Measure | Pair | Estimate | 95% bootstrap interval |
|---|---|---|---|
| Pearson | S–O | -0.623 | -0.669 to -0.575 |
| Spearman | S–O | -0.635 | -0.686 to -0.578 |
| Partial Pearson | S–O | -0.578 | -0.629 to -0.524 |
| Partial Spearman | S–O | -0.592 | -0.650 to -0.529 |

**c, Predictor-subspace overlap and Shapley allocation.**

| Family | Mean multiple $R^2$ on other families | Shapley $\Delta R^2$ |
|---|---|---|
| Executive-control | 0.714 | 0.0662 |
| Mnemonic/conceptual-social | 0.677 | 0.0297 |
| Perceptual/sensorimotor | 0.709 | 0.0035 |
| Organismic-affective | 0.605 | 0.0908 |

*Notes.* E, M, S and O denote executive-control, mnemonic/conceptual-social, perceptual/sensorimotor and organismic-affective families. Paper-level estimates use 577 independent paper or sample profiles with 5,000 bootstrap samples. Shapley values are conditional allocations, not independent effects. These descriptive dependencies do not represent within-brain coactivation; complete matrices are supplied as Supplementary Data.

**Supplementary Table 16 | Target information and partial information decomposition.**

**a, Total mutual information.**

| High-state fraction | Observed (bits) | Reference mean (bits) | Excess (bits) | $P$ or status |
|---|---|---|---|---|
| 40% | 0.069 | 0.046 | 0.022 | Descriptive |
| 50% | 0.072 | 0.057 | 0.015 | <0.001 |
| 60% | 0.046 | 0.038 | 0.008 | Descriptive |

**b, Source allocation.**

| Family | Shapley excess (bits) | FDR $q$ | Fully conditioned excess (bits) | FDR $q$ |
|---|---|---|---|---|
| Executive-control | -0.003 | 0.042 | -0.001 | 0.366 |
| Mnemonic/conceptual-social | 0.010 | 0.004 | 0.009 | 0.004 |
| Perceptual/sensorimotor | 0.005 | 0.015 | 0.004 | 0.136 |
| Organismic-affective | 0.003 | 0.014 | 0.005 | 0.032 |

**c, Partial-information types.**

| Type | Observed (bits) | Reference mean (bits) | Excess (bits) | Reference 95% limits (bits) | FDR $q$ |
|---|---|---|---|---|---|
| Synergy | 0.022 | 0.017 | 0.005 | 0.013–0.021 | 0.018 |
| Mixed-route redundancy | 0.043 | 0.032 | 0.011 | 0.026–0.038 | 0.003 |
| Individual-family redundancy | 0.007 | 0.009 | -0.001 | 0.005–0.013 | 0.564 |

*Notes.* References are 1,000 reporting-density-weighted ALE targets with study and reported-experiment focus counts and common $N = 20$ kernels preserved; EMSO templates were fixed. Synergy is available only from combined families. Mixed-route redundancy is available through alternative routes including at least one multi-family route, whereas individual-family redundancy is available separately from individual families. FDR is controlled within each named follow-up family. Fully conditioned increments include unique information and all synergy involving the focal family, so they are upper bounds on unique information.
